# Chromatin compartment dynamics in a haploinsufficient model of cardiac laminopathy

**DOI:** 10.1101/555250

**Authors:** Alessandro Bertero, Paul A. Fields, Alec S. T. Smith, Andrea Leonard, Kevin Beussman, Nathan J. Sniadecki, Deok-Ho Kim, Hung-Fat Tse, Lil Pabon, Jay Shendure, William S. Noble, Charles E. Murry

## Abstract

Pathogenic mutations in A-type nuclear lamins cause dilated cardiomyopathy, which is postulated to result from dysregulated gene expression due to changes in chromatin organization into active and inactive compartments. To test this, we performed genome-wide chromosome conformation analyses (Hi-C) in human induced pluripotent stem cell-derived cardiomyocytes (hiPSC-CMs) with a haploinsufficient mutation for lamin A/C. Compared to gene-corrected cells, mutant hiPSC-CMs have marked electrophysiological and contractile alterations, with modest gene expression changes. While large-scale changes in chromosomal topology are evident, differences in chromatin compartmentalization are limited to a few hotspots that escape inactivation during cardiogenesis. These regions exhibit upregulation of multiple non-cardiac genes including *CACNA1A*, encoding for neuronal P/Q-type calcium channels. Pharmacological inhibition of the resulting current partially mitigates the electrical alterations. On the other hand, A/B compartment changes do not explain most gene expression alterations in mutant hiPSC-CMs. We conclude that global errors in chromosomal compartmentation are not the primary pathogenic mechanism in heart failure due to lamin A/C haploinsufficiency.

**Summary:** Bertero et al. observe that lamin A/C haploinsufficiency in human cardiomyocytes markedly alters electrophysiology, contractility, gene expression, and chromosomal topology. Contrary to expectations, however, changes in chromatin compartments involve just few regions, and most dysregulated genes lie outside these hotspots.

**Condensed title:** Genomic effects of lamin A/C haploinsufficiency

## Introduction

Chromatin organization in the three-dimensional space has emerged as a key layer of mammalian gene expression control. The development of powerful technologies to map chromatin architecture, in particular genome-wide chromosome conformation capture (Hi-C; Lieberman-Aiden et al., 2009), has revealed that such 3D structure is complex, non-random, and hierarchical (Yu and Ren, 2017). Short-and long-range intra-chromosomal (*cis*) DNA interactions are generally constrained within topologically associating domains (TADs; Dixon et al., 2012; Nora et al., 2012; Sexton et al., 2012). TADs tend to interact based on their epigenetic status and transcriptional activity, thus dividing chromosomes into two types of large-scale compartments generally called active and inactive (A and B, respectively; Simonis et al., 2006; Lieberman-Aiden et al., 2009; Rao et al., 2014). Inter-chromosomal (*trans*) interactions predominantly involve the A compartment and are generally more limited, as chromosomes occupy distinct nuclear territories (Bolzer et al., 2005; Branco and Pombo, 2006; Kalhor et al., 2012).

Moving beyond the third dimension, there has been growing interest in understanding the functional changes in chromatin organization during development and in disease (Dekker et al., 2017). An emerging body of work has shown that TADs are largely invariant across cell types (Dixon et al., 2012, 2015; Fraser et al., 2015; Won et al., 2016; Schmitt et al., 2016; Fields et al., 2017). On the other hand, these same studies have established that pluripotent stem cell differentiation leads to a substantial degree of A/B compartment reorganization (e.g. ~20% of the genome), and that this is associated with important developmental changes in gene expression. Nevertheless, the precise mechanisms that regulate chromatin compartmentalization dynamics during development are still poorly understood (Adriaens et al., 2018). In the context of disease, disruption of TADs due to copy number variations or point mutations was shown to lead to congenital developmental disorders and cancers (Lupiáñez et al., 2015; Katainen et al., 2015; Redin et al., 2017; Sun et al., 2018). In contrast, whether dysregulation of A/B compartments plays a role in functional changes of gene expression leading to human disease is still unclear (Krumm and Duan, 2018).

The nuclear lamina has been proposed as a regulator of chromatin compartmentalization in development and disease (Buchwalter et al., 2018). The lamina lies along the inner nuclear membrane and is a complex mesh of nuclear intermediate filaments (A-and B-type lamins) and lamin-associated proteins. This structure provides key mechanical support to the nucleus, and is an important hub for the control of intracellular signaling (Dobrzynska et al., 2016). Moreover, the nuclear lamina interacts with large chromatin regions, aptly named lamin-associated domains (LADs; van Steensel and Belmont, 2017), which show heterochromatic features such as low gene density, enrichment for repressive histone marks, and poor transcriptional activity (Guelen et al., 2008). Several studies have established that such peripherally-located LADs strongly correlate with the B compartment, while the A compartment is predominantly in the nuclear interior (Kind et al., 2015; Stevens et al., 2017; Luperchio et al., 2017; Zheng et al., 2018). Both lamins and lamin-associated proteins, such as LBR (lamin B receptor), can directly interact with chromatin (Yuan et al., 1991; Taniura et al., 1995; Ye and Worman, 1994, 1996). Based on this and on the results of loss-of-function studies it has been proposed that the nuclear lamina tethers LADs at the nuclear periphery (Solovei et al., 2013; Harr et al., 2015; Luperchio et al., 2017). However, the precise role of A-and B-type lamins in chromatin compartmentalization is still being debated (Amendola and van Steensel, 2015; Zheng et al., 2015, 2018; Adriaens et al., 2018).

Elucidating the function of A-type lamins (lamin A and lamin C, which result from alternative splicing of the *LMNA* gene) is particularly important not only from a cell and developmental biology perspective, but also because of their involvement in human disease. *LMNA* mutations lead to a wide spectrum of conditions collectively referred to as laminopathies (Capell and Collins, 2006). Notably, nearly 80% of all reported *LMNA* mutations lead to isolated striated muscle disease, with another 10% leading to overlapping phenotypes that also affect striated muscles (Bertrand et al., 2011). The majority of patients with striated muscle laminopathies develop dilated cardiomyopathy (DCM), with variable skeletal muscle manifestations (Captur et al., 2018). Mutations in *LMNA* are generally considered the second-most most common cause of familial DCM, depending on the ethnicity of the population (Akinrinade et al., 2015; Haas et al., 2015; Tobita et al., 2018). Compared to other types of DCM, *LMNA*-DCM is quite atypical as it is characterized by early onset of life-threatening cardiac electrical abnormalities such as severe conduction system disease and/or atrial and ventricular arrhythmias (Van Rijsingen et al., 2012; Hasselberg et al., 2018; Kumar et al., 2016). Another peculiar aspect of *LMNA*-DCM is that not all patients go on to develop left ventricular dilatation and reduced contractile function, which are typical hallmarks of DCM (Van Berlo et al., 2005; Kumar et al., 2016; Tobita et al., 2018). Nevertheless, patients with *LMNA*-DCM are poorly responsive to medical treatments and show minimal beneficial left ventricular reverse remodeling (Tobita et al., 2018). This may be explained by the higher incidence of cardiac fibrosis in *LMNA*-DCM patients (van Tintelen et al., 2007b; Fontana et al., 2013; Tobita et al., 2018). Overall, *LMNA*-DCM (which throughout the rest of the manuscript we will refer to as “cardiac laminopathy”) is a more severe disease compared to other genetic DCM, with young onset, high penetrance, poor prognosis, and frequent need for heart transplantation as the only current available therapy (Hasselberg et al., 2018; Tobita et al., 2018).

Over the nearly 20 years since the first report linking mutations in *LMNA* to human disease (Bonne et al., 1999), three central non-mutually exclusive mechanisms have been hypothesized to underpin the pathogenesis of cardiac laminopathy: (1) impaired nuclear mechanoresistance *via* the nucleo-cytoplasmic network, or “mechanical hypothesis”; (2) alteration of lamin A/C-controlled intracellular signaling pathways, or “signaling hypothesis”; and (3) dysregulation of heterochromatin organization leading to gene expression alterations, or “chromatin hypothesis” (Worman and Courvalin, 2004; Cattin et al., 2013). While evidence supporting the first two hypotheses has accumulated over the years, and therapies targeting intracellular signaling alterations are being pre-clinically developed (Cattin et al., 2013; Captur et al., 2018), the possible involvement of chromatin dysregulation in cardiac laminopathy is still far from established (Adriaens et al., 2018). Indeed, while there have been reports of changes in the nuclear positioning of selected loci in patients with cardiac laminopathy (Meaburn et al., 2007; Mewborn et al., 2010), the functional consequences of such alterations on the disease pathogenesis are unclear. Moreover, these studies have relied on fibroblasts instead of cardiomyocytes, the primary cell type involved in cardiac laminopathy. Most importantly, to the best of our knowledge the 3D chromatin organization changes associated with cardiac laminopathy have not yet, been tested at a genome-wide level.

To address these limitations, we performed genome-wide chromosome conformation capture (Hi-C) and gene expression (RNA-seq) analyses to examine the changes in 3D chromatin architecture induced by a haploinsufficient *LMNA* mutation in cardiomyocytes derived from human induced pluripotent stem cells (hiPSC-CMs). We hypothesized that decreased expression of A-type lamins would lead to broad functional alterations in A/B compartmentalization leading to aberrant gene expression. We observed that mutant hiPSC-CMs show marked alterations in their electrophysiological and contractile phenotypes, broad dysregulation of gene expression, and alterations in large-scale chromosomal topology (strengthened separation between chromosome territories and between chromatin compartments). However, functional changes in A/B compartmentalization are limited to a few genomic hotspots that normally transition from the A to B compartment during cardiogenesis, but remain in A in lamin A/C haploinsufficient hiPSC-CMs. These hotspots exhibit upregulation of multiple non-cardiac genes, including the neuronal gene *CACNA1A*, which encodes for P/Q-type calcium channels. Of note, however, changes in A/B compartments do not explain the majority of gene expression changes, including the upregulation of the cardiac gene *CACNA1C*, which encodes for the L-type calcium channel. Pharmacological inhibition of P/Q-type calcium currents partially mitigates the field potential duration elongation observed in mutant hiPSC-CMs, while inhibition of L-type calcium currents has a more powerful effect. We conclude that, while *LMNA* haploinsufficient mutations functionally affect selected aspects of 3D chromatin organization in human cardiomyocytes, altered A/B compartmentalization does not represent the primary mechanism directly leading to gene expression changes and disease pathogenesis.

## Results

### Generation of an *in vitro* model of cardiac lamin A/C haploinsufficiency

To investigate the role of chromatin dynamics in cardiac laminopathy, we took advantage of hiPSCs bearing a heterozygous nonsense mutation in *LMNA* predicted to cause premature truncation of both lamin A and lamin C splicing isoforms (c.672C>T, resulting in p.Arg225*, which we will refer to as R225X; Fig. 1A). This hiPSC line was previously derived from a 56-year-old male patient who developed severe cardiac conduction disease evolving into heart failure, a condition that segregated within the family with autosomal dominant inheritance of the R225X mutation (Siu et al., 2012). This same mutation has been reported in multiple other cohorts with similar symptoms (van Tintelen et al., 2007a; Saga et al., 2009; Jakobs et al., 2001), establishing it as a *bona fide* genetic cause of cardiac laminopathy.

**Figure 1.**
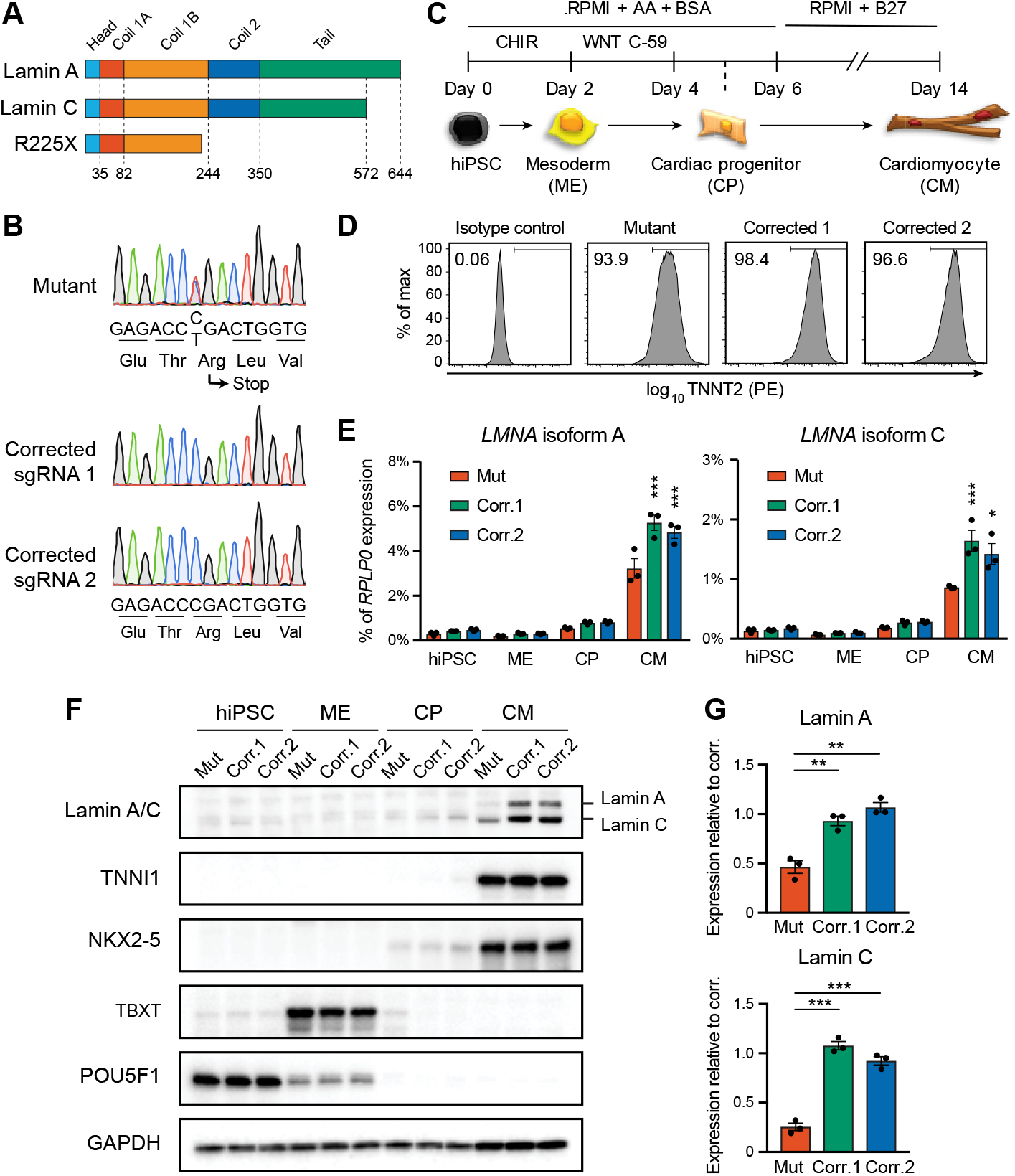
Generation of lamin A/C haploinsufficient hiPSC-CMs. (**A**) Predicted effect of the *LMNA* R225X mutation on the two splicing products lamin A and C. Key protein domains and their location along the amino acid sequence are indicated. (**B**) Sanger sequencing of *LMNA* exon 4 in hiPSCs with heterozygous R225X mutation (top), or in hiPSCs obtained after CRISPR/Cas9-based scarless correction of the mutation (bottom). (**C**) Schematic of the protocol for step-wise directed differentiation of hiPSC-CMs. CHIR: CHIR99021 (WNT activator); WNT C-59: WNT inhibitor; AA: ascorbic acid. (**D**) Quantification of cardiac differentiation efficiency by flow cytometry for cardiac troponin T (TNNT2) on hiPSC-CMs at day 14 of differentiation. The percentage of TNNT2+ cells is reported. (**E**) RT-qPCR analyses at the indicated stages of hiPSC-CM differentiation (see panel C). Differences *versus* mutant were calculated by two-way ANOVA with post-hoc Holm-Sidak binary comparisons (* = p < 0.05, *** = p < 0.001; n = 3 differentiations; average ± SEM). (**F**) Representative western blot for lamin A/C and differentiation markers during iPSC-CM differentiation. (**G**) Quantification of lamin A/C expression in hiPSC-CMs from western blot densitometries (n = 3 differentiations; average ± SEM). Differences versus mutant were calculated by one-way ANOVA with post-hoc Holm-Sidak binary comparisons (** = p < 0.01, *** = p < 0.001; n = 3 differentiations; average ± SEM). Throughout the figure (and in all other figures), Mut or Mutant indicates *LMNA* R225X hiPSCs, and Corr.1/2 or Corrected 1/2 indicate the two isogenic corrected control *LMNA* R225R hiPSC lines.

It is well established that variability among hiPSC lines can profoundly influence both molecular and cellular phenotypes (Ortmann and Vallier, 2017), which is particularly evident when assessing the complex electrophysiology and contractile properties of hiPSC-CMs (Sala et al., 2017). Thus, we decided to generate isogenic control hiPSCs by correcting the R225X mutation back to the wild-type allele, a strategy currently considered the gold standard to determine the association between genotype and phenotype in hiPSC-CMs (Bellin et al., 2013; Kodo et al., 2016; Mosqueira et al., 2018). By leveraging existing methods (Yusa, 2013), we designed a two-step scarless gene editing strategy relying on CRISPR/Cas9-facilitated homologous recombination of a targeting vector containing the wild-type allele in the 3’ homology arm, and an excisable piggyBac drug resistance cassette (Fig. S1A-B). To control for the potential variability between hiPSC sublines, we obtained two isogenic control hiPSCs using distinct sgRNA sequences (Fig. 1B and S1C-D). The resulting hiPSCs, which we will refer to as Corr.1 and Corr.2 (short for Corrected 1 and Corrected 2) were karyotypically normal (Fig. S1E).

Lamin A/C is expressed at very low levels in human pluripotent stem cells, and is upregulated during differentiation (Constantinescu et al., 2006; Liu et al., 2011). Given that hiPSC-CMs are quite immature (Yang et al., 2014), we sought to first confirm whether hiPSC-CMs express substantial levels of lamin A/C. Further, we tested the consequence of the R225X mutation on lamin A/C expression at the transcript and protein level. For this, we differentiated hiPSCs into hiPSC-CMs using an established protocol based on the temporal modulation of WNT signaling using small molecules (Burridge et al., 2014; Fields et al., 2017; Fig. 1C). Both mutant and corrected hiPSCs could be differentiated with high efficiency, as measured by flow-cytometry for TNNT2 (Fig. 1D, Video S1, and Video S2; 91.6% ± 3.2%, 94.4% ± 2.5%, and 91.3% ± 3.1% for Mutant, Corr.1, and Corr.2, respectively; average ± SEM of n = 4 independent differentiations). Western blot and quantitative reverse transcription PCR (RT-qPCR) confirmed that all hiPSC lines underwent the expected developmental progression through mesoderm and cardiac progenitors before reaching a cardiomyocyte phenotype (Fig. 1F and S2A). Lamin A/C was upregulated specifically in hiPSC-CMs, and mutant lines showed significantly reduced levels of both mRNA and protein compared to both corrected controls (Fig. 1E-G). Of note, no detectable protein truncation was detected by western blot (Fig. S2B), and lamin A/C expression in mutant hiPSC-CMs also proved to be reduced when compared to cardiomyocytes generated from an unrelated hiPSC line derived from a healthy individual (Fig. S2C). Finally, other minor *LMNA* isoforms were undetectable or nearly undetectable by RT-qPCR in both control and mutant hiPSC-CMs, excluding possible compensatory mechanisms (data not shown). These gene expression data agree with earlier findings from analysis of skin fibroblasts bearing the R225X heterozygous mutation (Siu et al., 2012), and indicate that such premature nonsense mutation leads to lamin A/C haploinsufficiency presumably due to nonsense-mediated decay of both the lamin A and lamin C transcripts. Collectively, we established a robust *in vitro* model to study cardiac laminopathy due to lamin A/C haploinsufficiency in developing cardiomyocytes.

### Lamin A/C haploinsufficiency alters hiPSC-CM automaticity and prolongs membrane depolarization

Before exploring the effect of lamin A/C haploinsufficiency on chromatin dynamics, we wished to confirm a phenotypic effect on cardiac physiology in developing hiPSC-CMs. Indeed, cardiac laminopathy is a disease that manifests in the third to fourth decade of life, and previous efforts to model the disease in immature cardiomyocytes have been focused on studying nuclear dysmorphology, cellular senescence, and susceptibility to apoptosis (Siu et al., 2012; Lee et al., 2017). Since electrical abnormalities are the primary and most characteristic manifestations of cardiac laminopathy (Van Rijsingen et al., 2012; Hasselberg et al., 2018; Kumar et al., 2016), we began by assessing the electrophysiological properties of mutant hiPSC-CMs. For this, we first used multi-electrode arrays (MEAs) to measure the extracellular electric field potential elicited by spontaneously contracting monolayers of hiPSC-CMs at day 30 of differentiation. Mutant cells showed a number of alterations compared to both corrected controls. Most notably, the beat rate proved highly erratic and prone to prolonged pauses (Fig. 2A). Moreover, even when the analysis was focused on the periods showing the highest consistency, the beat rate was still more irregular and reduced (Fig. 2B). The amplitude of field potential changes was elevated more than 2-fold compared to controls, which is indicative of stronger depolarizing ion currents (Fig. 2B). Along the same lines, the field potential duration (FPD), which indicates the interval between depolarization and repolarization, was prolonged by ~60% (Fig. 2B). This finding held true even when the FPD was corrected for the beat period (FPDc; Fig. 2B), an established way to robustly compare the FPD across multiple conditions (Rast et al., 2016; Asakura et al., 2015). Of note, despite all of these alterations, the conduction velocity across the monolayer was not affected (Fig. 2B), indicating that intercellullar electrical coupling was preserved in mutant cells.

**Figure 2.**
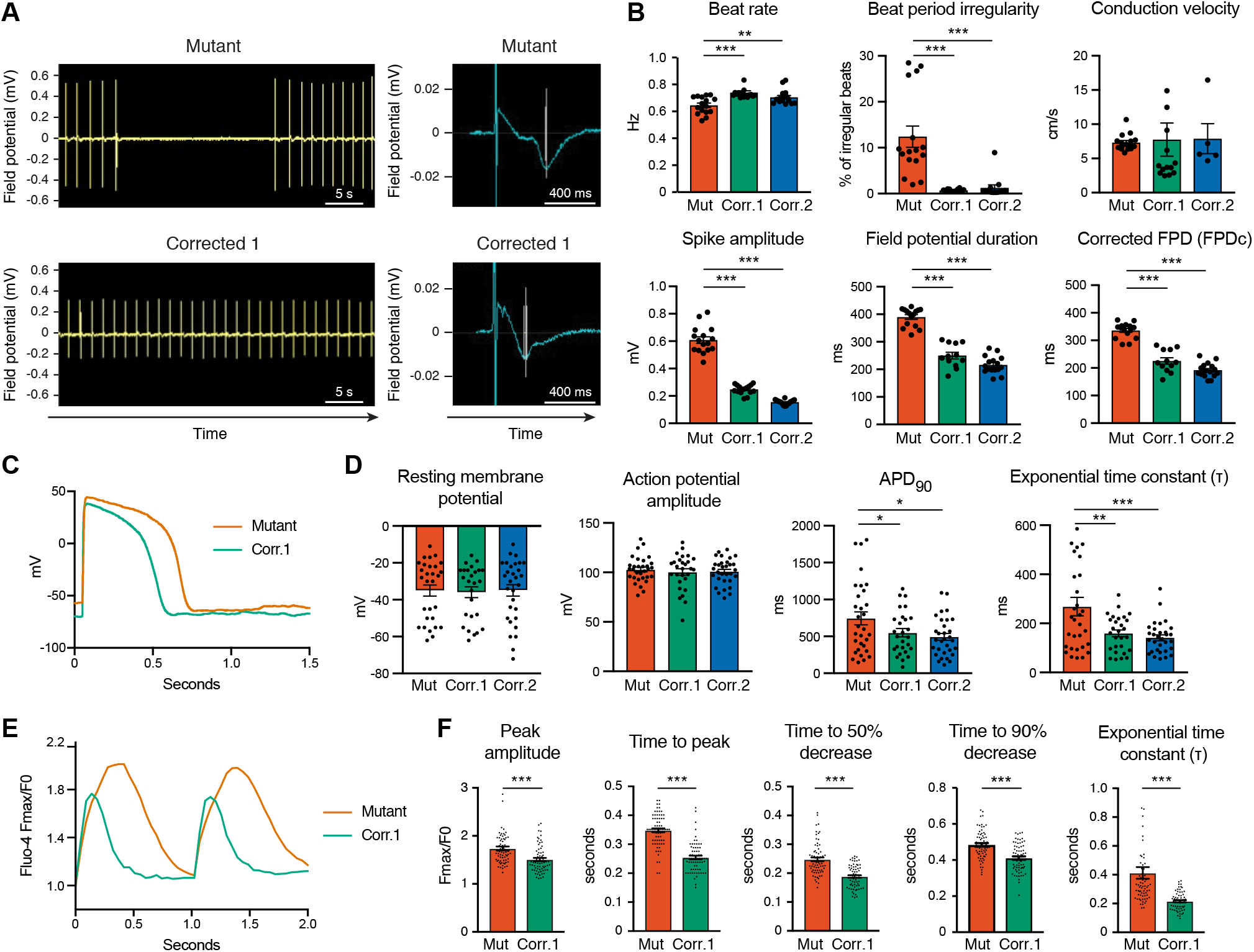
Electrophysiological properties of lamin A/C haploinsufficient hiPSC-CMs. (**A**) Representative traces from MEA recordings of spontaneous electrical activity in hiPSC-CM monolayers. On the right, the average field potential changes during an individual beat are reported, and depolarization and repolarization timings are indicated by vertical lines. (**B**) Representative quantifications of electrophysiological properties from MEA analyses. Differences *versus* mutant were calculated by one-way ANOVA with post-hoc Holm-Sidak binary comparisons (** = p < 0.01, *** = p < 0.001; n = 5-16 wells; average ± SEM). (**C**) Representative voltage recordings by whole-cell patch clamp during evoked action potentials in individual hiPSC-CMs. (**D**) Quantifications of electrophysiological properties from whole-cell patch clamp analyses. Differences *versus* mutant were calculated by one-way ANOVA with post-hoc Holm-Sidak binary comparisons (* = p < 0.05, ** = p < 0.01, *** = p < 0.001; n = 26-30 cells from two differentiations; average ± SEM). (**E**) Representative optical recordings of calcium fluxes with Fluo-4 in hiPSC-CM monolayers electrically paced at 1 Hz. (**F**) Representative quantifications of calcium fluxes properties. Differences *versus* mutant were calculated by unpaired t-test (*** = p < 0.001; n = 69-70 cells; average ± SEM).

To further characterize the electrophysiological properties of mutant hiPSC-CMs, we performed whole-cell patch clamp recordings of voltage changes occurring during action potentials firing in individual cardiomyocytes. To promote a mature phenotype and enhance cell viability during the invasive patch procedure, we cultured hiPSC-CMs onto biomimetic anisotropic nanopatterns (Carson et al., 2016; Macadangdang et al., 2015). While a number of parameters were unaffected in mutant cells (such as maximum diastolic potential, action potential amplitude, mean diastolic potential, and repolarization amplitude; Fig. 2D and data not shown), the action potential duration and exponential time constant for the action potential decay were increased (Fig. 2C-D). This observation provided a cell-autonomous explanation for the increased FPDc in cell monolayers, and indicated that this phenotype is maintained in maturing hiPSC-CMs.

Finally, we measured the effects of excitation abnormalities in mutant cells on their intracellular calcium dynamics using the fluorescent calcium reporter Fluo-4. Electrically-paced mutant cell monolayers showed stronger and longer calcium peaks (Fig. 2E-F, Video S3, and Video S4), in agreement with MEA and patch clamp data. Overall, these findings indicate that lamin A/C haploinsufficiency in developing cardiomyocytes leads to altered automaticity and prolonged membrane depolarization leading to more robust calcium transients.

### Systolic hyperfunction and diastolic dysfunction in lamin A/C haploinsufficient hiPSC-CMs

Since changes in intracellular calcium concentrations are the primary determinant of hiPSC-CM contractility, we examined the effect of lamin A/C haploinsufficiency on this process. First, we performed correlation-based contraction quantification (CCQ) analyses to measure the cellular displacement associated with cardiac contraction in electrically-paced hiPSC-CM monolayers (Macadangdang et al., 2015). Mutant cells showed ~50% stronger contractions and a delayed relaxation time (Fig. 3A-B, Video S5, and Video S6), consistent with their stronger and prolonged calcium fluxes.

**Figure 3.**
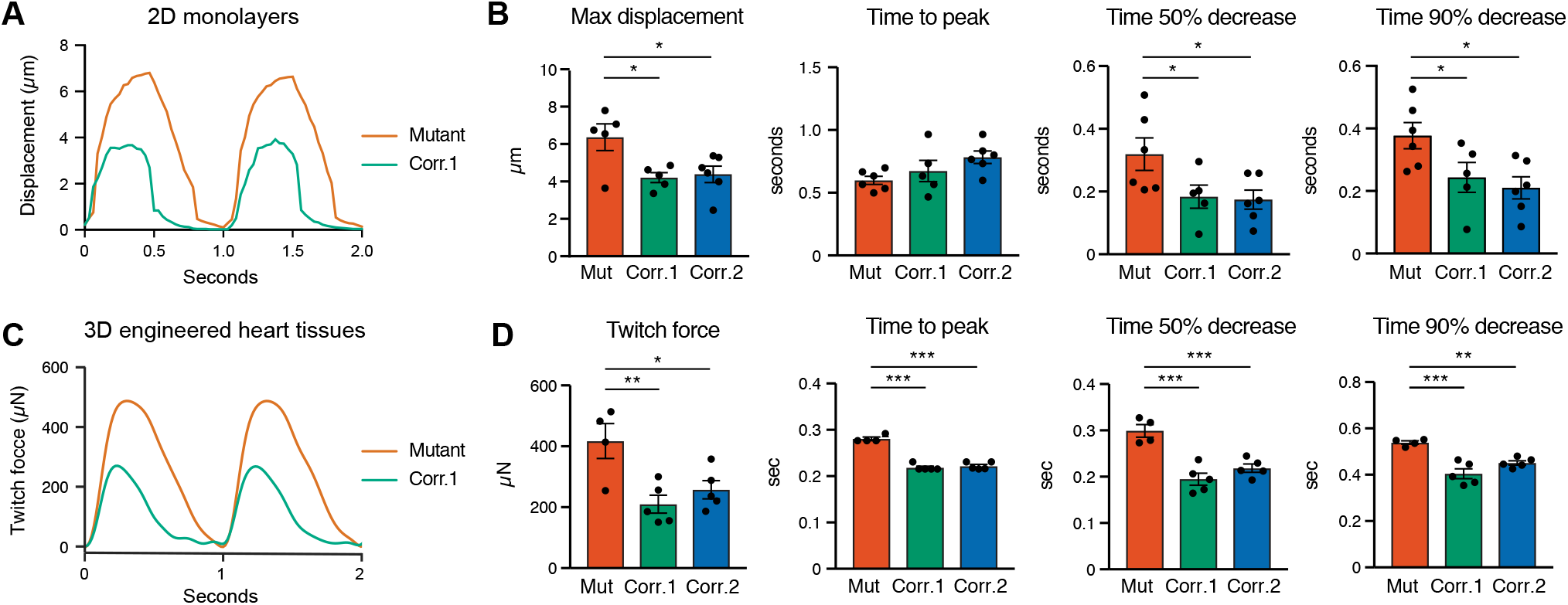
Contractile properties of lamin A/C haploinsufficient hiPSC-CMs. (**A**) Representative measurements of cellular displacement during contraction of hiPSC-CM monolayers electrically paced at 1 Hz. (**B**) Representative quantifications of cell contractility from analyses of optical recordings. Differences *versus* mutant were calculated by one-way ANOVA with post-hoc Holm-Sidak binary comparisons (* = p < 0.05; n = 5-6 field of views; average ± SEM). (**C**) Representative measurements of twitch force during contraction of 3D-EHTs electrically paced at 1 Hz. (**D**) Representative quantifications of tissue contractility from analyses of optical recordings. Differences *versus* mutant were calculated by one-way ANOVA with post-hoc Holm-Sidak binary comparisons (* = p < 0.05, ** = p < 0.01, *** = p < 0.001; n = 4-5 3D-EHTs; average ± SEM).

To validate these findings in a more physiologically-relevant model, we performed biomechanical assessments of contractility in electrically-paced three-dimensional engineered heart tissues (3D-EHTs), an established model to promote cardiac maturation by providing topological, mechanical, and multicellular cues (Leonard et al., 2018; Ruan et al., 2016). In agreement with observations in 2D monolayers, mutant 3D-EHTs showed ~2-fold stronger and prolonged contractions with a markedly impaired relaxation kinetic (Fig. 3C-D, Video S7, and Video S8). Collectively, we concluded that lamin A/C haploinsufficiency leads to systolic hyperfunction and diastolic dysfunction in both early and maturing cardiomyocytes.

### Lamin A/C haploinsufficiency causes dysregulation of specific ion channel genes and broad gene expression changes

To explore the molecular mechanisms that might explain the electrophysiological and contractile phenotypes observed in lamin A/C haploinsufficient hiPSC-CMs, we first monitored the expression of the genes encoding key ion-handling proteins involved in the generation of action potentials and in excitation-contraction coupling (Amin et al., 2010; Eisner et al., 2017). RT-qPCR analyses excluded a dysregulation of a large number of such genes, including the voltage-gated sodium channel *SCN5A*, the sodium-calcium exchanger *NCX*, and the sarcoplasmic calcium ATPase *ATP2A2* (also known as SERCA2a; Fig. S2D). On the other hand, mutant cells showed a significant upregulation of *CACNA1C*, and downregulation of *KCNQ1* (Fig. 4A). *CACNA1C* encodes for the pore-forming subunit of L-type calcium channels which mediate I_Ca,L_, the main source of inward depolarizing current during phase 2 (plateau) of the action potential (Bodi et al., 2005). On the other hand, I_KS_, the outward potassium current resulting from the potassium channel encoded by *KCNQ1*, antagonizes I_Ca,L_ in phase 2 by initiating membrane repolarization (Peroz et al., 2008). Thus, the combined effects of *CACNA1C* upregulation and *KCNQ1* downregulation could explain the prolonged membrane depolarization observed in mutant cells.

**Figure 4.**
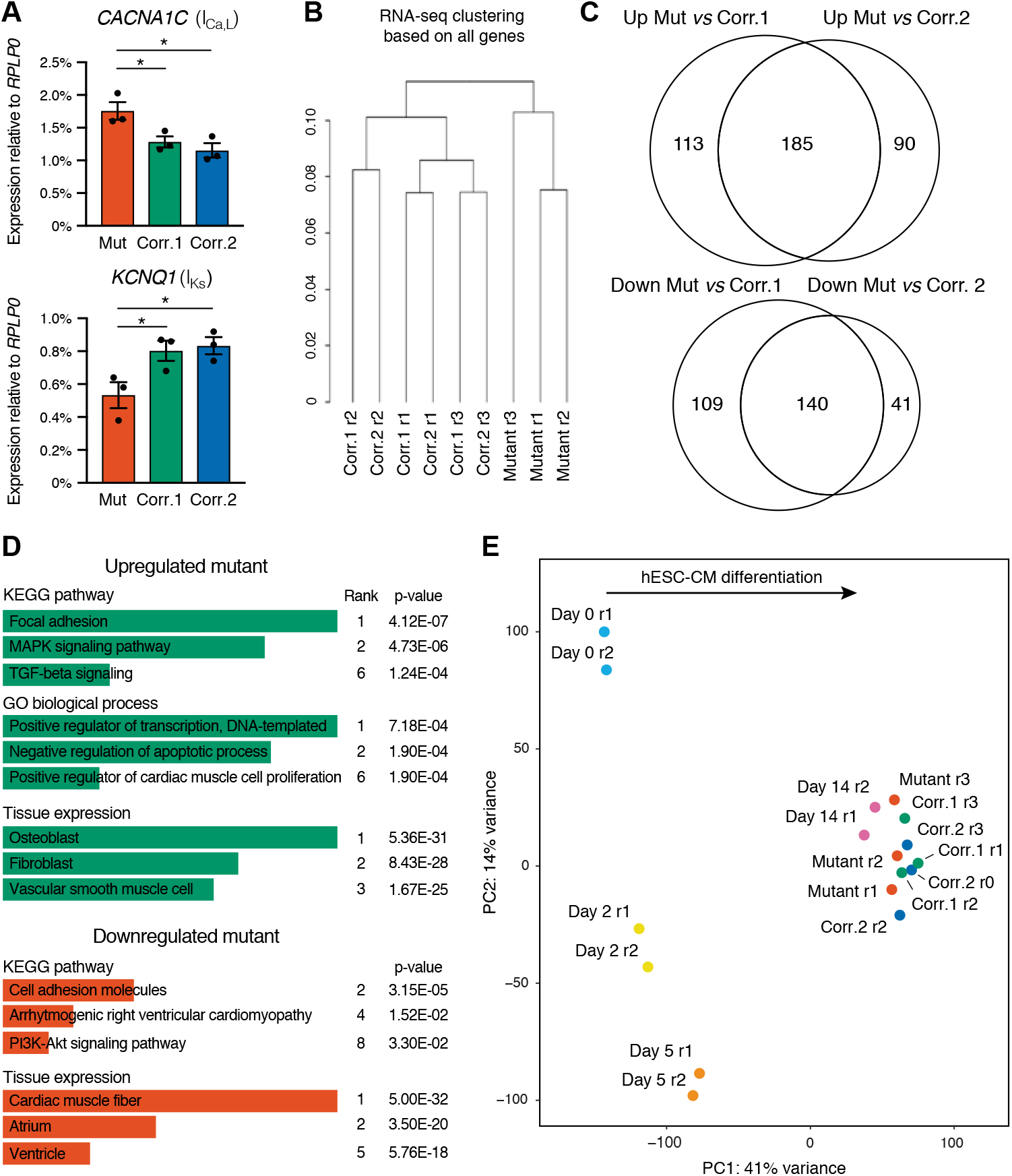
Gene expression changes in lamin A/C haploinsufficient hiPSC-CMs. (**A**) RT-qPCR analyses in hiPSC-CMs at day 14 of differentiation. Differences *versus* mutant were calculated by one-way ANOVA with post-hoc Holm-Sidak binary comparisons (* = p < 0.05, n = 3 differentiations; average ± SEM). (**B**) Hierarchical clustering of hiPSC-CMs analyzed by RNA-seq based on all expressed genes. Biological replicates from 3 independent differentiations were analyzed (r1, r2, r3). (**C**) Overlap in genes up-or downregulated in mutant hiPSC-CMs *versus* hiPSC-CMs from the two corrected control lines (fold-change > 2 and q-value < 0.05; Table S1). (**D**) Selected results from ontology and pathway enrichment analyses of genes consistently up-or downregulation in mutant hiPSC-CMs. For each term, the rank and the corresponding p-value are reported (terms ranked by combined score; Table S2). (**E**) Linear dimensionality reduction by principal component analysis of RNA-seq data of mutant and corrected hiPSC-CMs, and hESC-CMs sampled at different time points of differentiation (Fields et al., 2017). The amount of variance captured by each of the two main principal components (PC) is reported, and the biological interpretation for the PC1 axis is indicated.

We then expanded these gene expression analyses genome-wide by performing RNA sequencing of hiPSC-CM monolayers at day 14 of differentiation on biological triplicates (RNA-seq; Table S1). While cells from the two corrected control lines clustered closely and showed remarkably similar gene expression profiles (29 upregulated and 71 downregulated genes between Corr.1 and Corr.2; fold-change > 2 and q-value < 0.05; Fig. 4B and Table S1), mutant cells clustered separately due to substantial gene up-and downregulation (Fig. 4B-C and Table S1). To increase the robustness of our subsequent analyses, we only considered genes as dysregulated if they had significant expression changes in mutant hiPSC-CMs *versus* both corrected controls (185 upregulated and 140 downregulated; Fig. 4C and Table S1). Ontology and pathway enrichment analyses on these gene lists revealed that upregulation in mutant cells was significantly associated with: (1) focal adhesion, MAPK, and TGFß pathways; (2) transcriptional activation, positive regulation of cardiac differentiation, and inhibition of apoptosis; (3) non-cardiac lineage expression (such as fibroblast and smooth muscle; Fig. 4D and Table S2). Notable examples of genes within these categories include: (1) *PDGFA*, *EGF*, and *GDF7*; (2) *TBX3*, *TBX20*, and *HAND1*, (3) *CTGF/CCN2*, *MYH9*, and *ACTA2*. In contrast, downregulated genes were enriched in cardiac transcripts and factors involved in cardiomyopathy and PI3K pathways, such as *TNNT3*, *TNNI3*, *SGCA*, *NGFR*, and *PDGFD* (Fig. 4D and Table S2). We also compared these RNA-seq data with those we recently obtained from human embryonic stem cells (hESCs) profiled at different time points of cardiac differentiation (Fields et al., 2017). Linear dimensionality reduction by principal component analysis showed that both mutant and corrected hiPSC-CMs clustered closely to hESC-CMs (Fig. 4E). This indicated that despite dysregulation of >300 genes, mutant hiPSC-CMs were not globally developmentally delayed from a transcriptional standpoint. These data suggest that lamin A/C haploinsufficiency in developing hiPSC-CMs leads to dysregulation of multiple signaling pathways, in agreement with earlier findings from mouse models (Arimura et al., 2005; Muchir et al., 2007; Ramos et al., 2012; Choi et al., 2012). Further, it prevents efficient silencing of some cardiac developmental regulators and alternative lineage genes, while specific cardiac genes are not fully activated.

### Lamin A/C haploinsufficiency strengthens the separation between chromosome territories and between chromatin compartments

Having established that lamin A/C haploinsufficiency results in marked changes in both gene expression and cellular physiology in developing hiPSC-CMs, we tested whether some of these phenotypes could be explained by changes in chromatin topology. In order to explore this aspect at a genome-wide level, we took advantage of *in situ* DNase Hi-C to capture all pairwise interactions between any two genomic regions (Ramani et al., 2016). We analyzed two independent batches of hiPSC-CMs at day 14 of differentiation from the mutant and two corrected lines, and we confirmed that the resulting Hi-C data were of high quality and reproducible across biological replicates, with samples clustering separately based on their genotype (Fig. 5A and Table S3). We then began exploring the global properties of chromatin topology and noticed that mutant cells showed an increased ratio of genomic interactions with the same chromosome (in *cis*) over those involving different chromosomes (in *trans*; Fig. 5B). Moreover, *trans* interactions were distributed differently: while all hiPSC-CMs showed the expected pattern of preferential self-association between small, gene-rich chromosomes (especially chromosomes 16, 17, 19, 20, and 22) and between large, gene-poor chromosomes (such as chromosomes 1 to 8; Fig. S3A; Lieberman-Aiden et al., 2009), this property was even more striking in mutant cells (Fig. 5C). On the contrary, interactions between small and large chromosomes were less frequent (Fig. 5C). Thus, lamin A/C haploinsufficiency reinforces the separation between chromosome territories and their segregation based on size and gene density.

**Figure 5.**
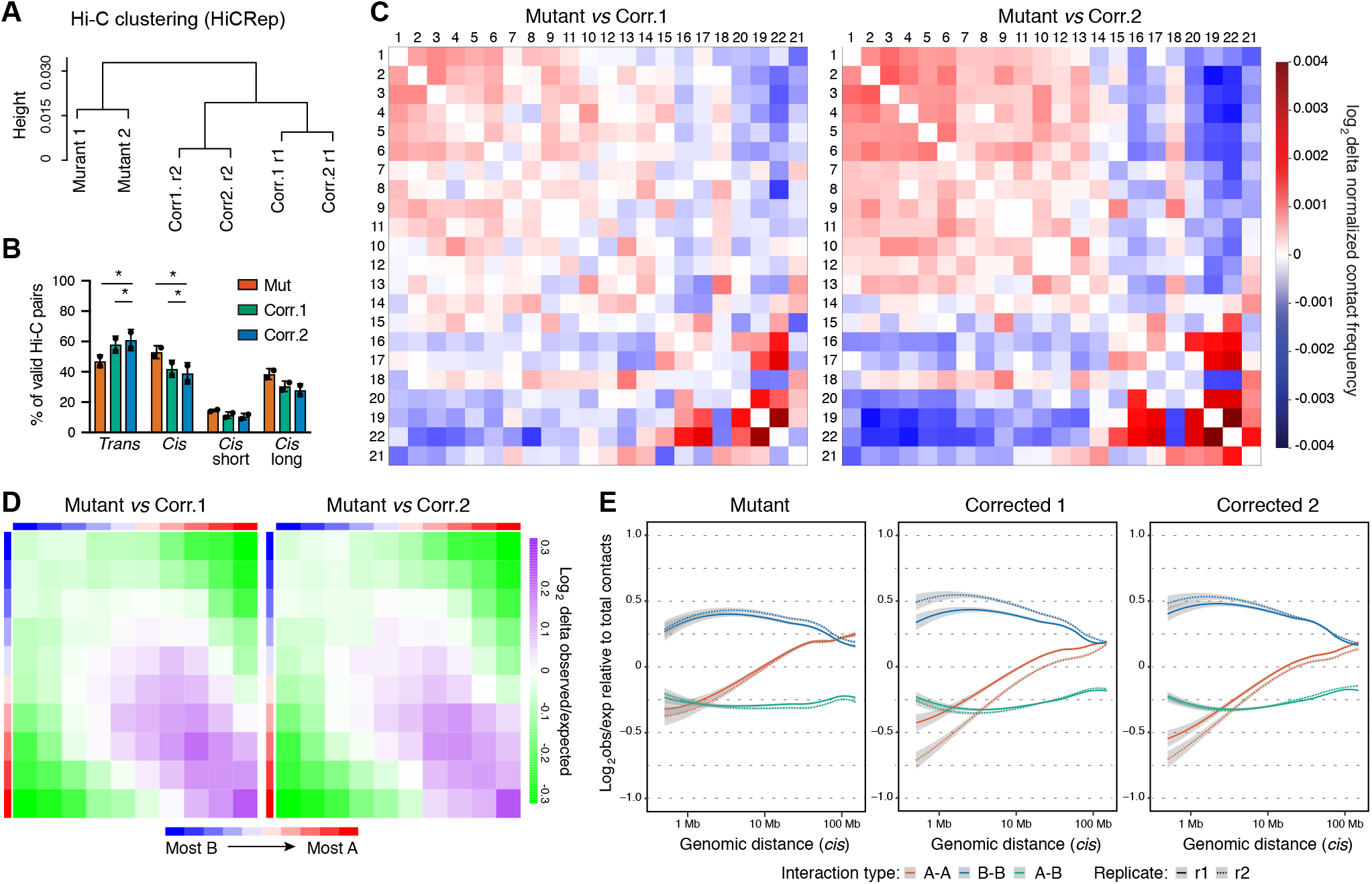
Global properties of chromatin topology in lamin A/C haploinsufficient hiPSC-CMs. (**A**) Hierarchical clustering of hiPSC-CMs analyzed by *in situ* DNase Hi-C based on similarity scores between the genomic contact matrices calculated with HiCRep. Biological replicates from 2 independent differentiations were analyzed (r1, r2). (**B**) Proportion of genomic interactions between different chromosomes (*trans*) or within the same chromosome (*cis*) involving distances < 20 Kb (*cis* short) or > 20 Kb (*cis* long; Table S3). Differences *versus* mutant were calculated by two-way ANOVA with post-hoc Holm-Sidak binary comparisons (* = p < 0.05; n = 2 differentiations; average ± SEM). (**C**) Representative heatmaps of differential contact matrices between chromosomes. Autosomes are ranked based on their size from left to right and top to bottom. (**D**) Representative heatmaps of differential *cis* interactions between active (A) and inactive (B) chromatin compartments. 500 Kb genomic bins were assigned to ten deciles based on their PC1 score from the linear dimensionality reduction of the Hi-C matrix (from most B to most A; Table S4), and average observed/expected distance normalized scores for each pair of deciles were calculated. (**E**) Probability of *cis* genomic contacts over increasing genomic distance for regions in homotypic (A-A or B-B) or heterotypic (A-B) chromatin compartments. Values are normalized to all contacts observed at a given distance, and LOESS curves are shown (gray background: 95% confidence bands).

We then analyzed the separation of chromatin domains into the active (A) or inactive (B) compartment. For this, we computed the first principal component (PC1) from the contact matrix using bins of 500 Kb (Table S4), a well-established method to determine chromatin compartments based on their preferential association (Imakaev et al., 2012; Lieberman-Aiden et al., 2009). In agreement with our previous observations in hESC-CMs (Fields et al., 2017), *trans* interactions between chromatin domains favored A-A compartments, and this was not affected by lamin A/C haploinsufficiency (Fig. S3B). In *cis*, mutant cells showed stronger interactions between A-A compartments (Fig. 5D and Fig. S3B), particularly due to increased short-range contacts within 0.5-1 Mb (Fig. 5E). In contrast, interactions between heterotypic regions (A-B) were reduced in mutant cells (Fig. 5D and Fig. S3B), particularly for long-range contacts >10 Mb (Fig. 5E). Collectively, lamin A/C haploinsufficiency reinforces the separation between the active and inactive chromatin compartments.

### Incomplete transitions from the active to inactive chromatin compartment in lamin A/C haploinsufficient hiPSC-CMs

To assess the effect of lamin A/C haploinsufficiency on chromatin compartmentalization, we identified genomic bins with significantly different A/B compartment scores and switching from active to inactive or *vice versa* between at least two conditions (Table S4). We noticed that the vast majority of compartment transitions were observed for mutant hiPSC-CMs *versus* each corrected control (Fig. 6A), and that B to A inversions were more common than A to B ones (42 and 27, respectively; Table S4). We observed that, 63% of A to B transitions in mutant cells involved the X chromosome, while B to A changes showed a notable concentration on chromosome 19 but were otherwise evenly spread across 13 additional chromosomes (Fig. 6B, Fig. S4A, and Table S4). Overall, compartment changes involved approximately 1.2% of the genome, indicating that chromatin compartment dysregulation in mutant cells is not widespread but is actually highly restricted.

**Figure 6.**
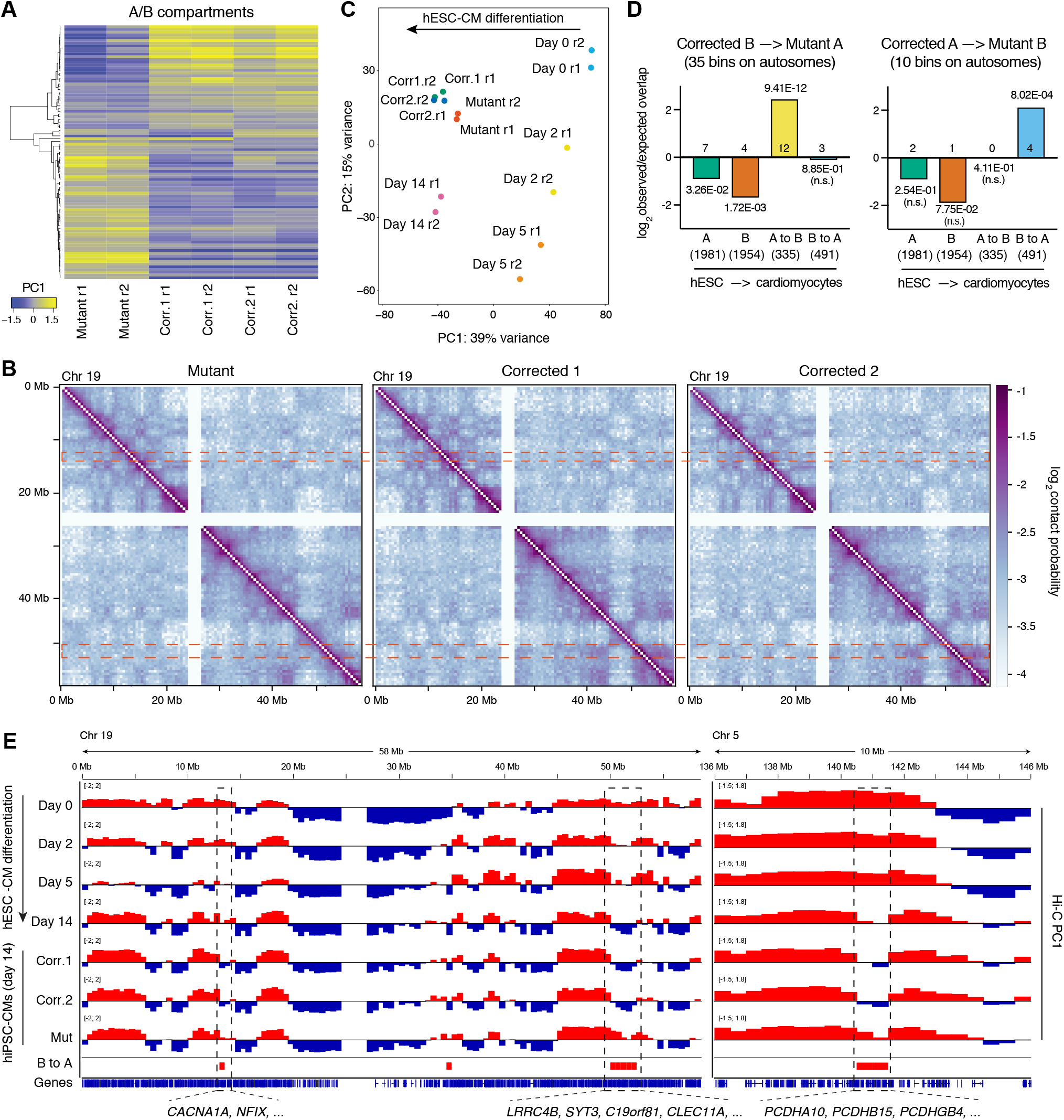
Chromatin compartment transitions in lamin A/C haploinsufficient hiPSC-CMs. (**A**) Heatmap of all significantly different A/B compartment scores (Hi-C matrix PC1; p < 0.05 by one-way ANOVA; n = 2 differentiations; Table S4) in 500 Kb bins that changed PC1 sign between two or more conditions. Positive and negative PC1 indicate A and B compartmentalization, respectively. (**B**) Representative log-transformed contact probability maps for chromosome 19. Topologically associating domains (TADs) are visible as squares along the diagonal. TADs within the same compartment interact off the diagonal as indicated by the symmetrical checkerboard patterns. Two genomic regions which show different compartmentalization in mutant hiPSC-CMs are indicated by dashed boxes to highlight the differences in contact probabilities with other genomic regions off the diagonal. (**C**) Linear dimensionality reduction by principal component analysis of A/B compartment scores from Hi-C data of mutant and corrected hiPSC-CMs, and hESC-CMs sampled at different time points of differentiation (Fields et al., 2017). The amount of variance captured by each of the two main principal components (PC) is reported, and the biological interpretation for the PC1 axis is indicated. (**D**) Significance of the overlap between changes in A/B compartments in mutant hiPSC-CMs and those occourring during hESC-CM differentiation. The number of genomic bins within each of the categories is indicated, and p-values were calculated by chi-squared tests. Note that only autosomes were considered. (**E**) Representative genomic tracks of chromatin compartmentalization for chromosome 19 and a section of chromosome 5. Positive and negative Hi-C matrix PC1 scores are shown in red and blue, and indicate 500 Kb genomic bins in the A and B compartments, respectively. Genomic regions that transition from A to B during hESC-CM differentiation but remain in A in mutant hiPSC-CMs (noted as B to A) are indicated by dashed boxes. Selected genes found within such regions are listed (refer to Fig. 7 and Fig. S4).

To gain insight into the relationship between dysregulated regions and normal chromatin compartment dynamics during cardiomyocyte differentiation, we integrated Hi-C data from mutant and corrected hiPSC-CMs with those we previously generated at different time points throughout hESC-CM differentiation (Fields et al., 2017). Linear dimensionality reduction of A/B compartment scores for all samples confirmed that mutant hiPSC-CMs cluster separately from both corrected controls (Fig. 6C). Moreover, this analysis revealed that based on the first principal component (which explained 39% of the variance and ordered hESC samples based on their developmental progression), mutant cells were mildly developmentally delayed from a chromatin compartmentalization standpoint (Fig. 6C). Accordingly, we observed a strong and significant enrichment for domains that normally transition from A to B during cardiac differentiation but remain in A in mutant cells (Fig. 6D). Notable examples of such behavior involved two 0.5 and 2.5 Mb-long portions of chromosome 19 (corresponding to 19p13.13 and 19q13.33, respectively), and a 1 Mb region on chromosome 5 (5q31.3; Fig. 6B and 6E). We also observed a weaker enrichment for the opposite dynamic (impaired B to A transitions; Fig. 6D), but we note that this analysis was limited to autosomes since compartment transitions of the X chromosome could not be assessed in female hESC-CMs due to the confounding factor of X inactivation (Fields et al., 2017). In sum, lamin A/C haploinsufficiency in developing hiPSC-CMs results in highly selective dysregulation of chromatin compartmentalization, particularly for a handful of genomic hotspots that fail to transition from the active to inactive compartment. We will refer to these as lamin A/C-sensitive B domains.

### Dysregulation of lamin A/C-sensitive B domains leads to ectopic expression of non-cardiac genes

We then assessed the functional consequences of compartment dysregulation due to lamin A/C haploinsufficiency. Strikingly, we observed almost no overlap between genes within lamin A/C-sensitive domains and genes significantly and strongly up-or downregulated in mutant cells (Fig. S4B). Accordingly, there were no significant changes in the average expression of genes found in lamin A/C-sensitive domains in mutant *versus* corrected controls (Fig. 7A). Nevertheless, we noticed a small number of genes located in lamin A/C-sensitive B domains that were expressed at very low levels in corrected hiPSC-CMs and upregulated in mutant cells (29 genes with an average fold-change > 2; Fig. 7A and Fig. S4C). These genes were significantly enriched for three chromosome locations (Fig. 7B), two of which corresponded to the lamin A/C-sensitive hotspots 5q31.3 and 19q13.33 (Fig. 6E), and were associated with neuronal development (Fig. 7B, and Table S5).

**Figure 7.**
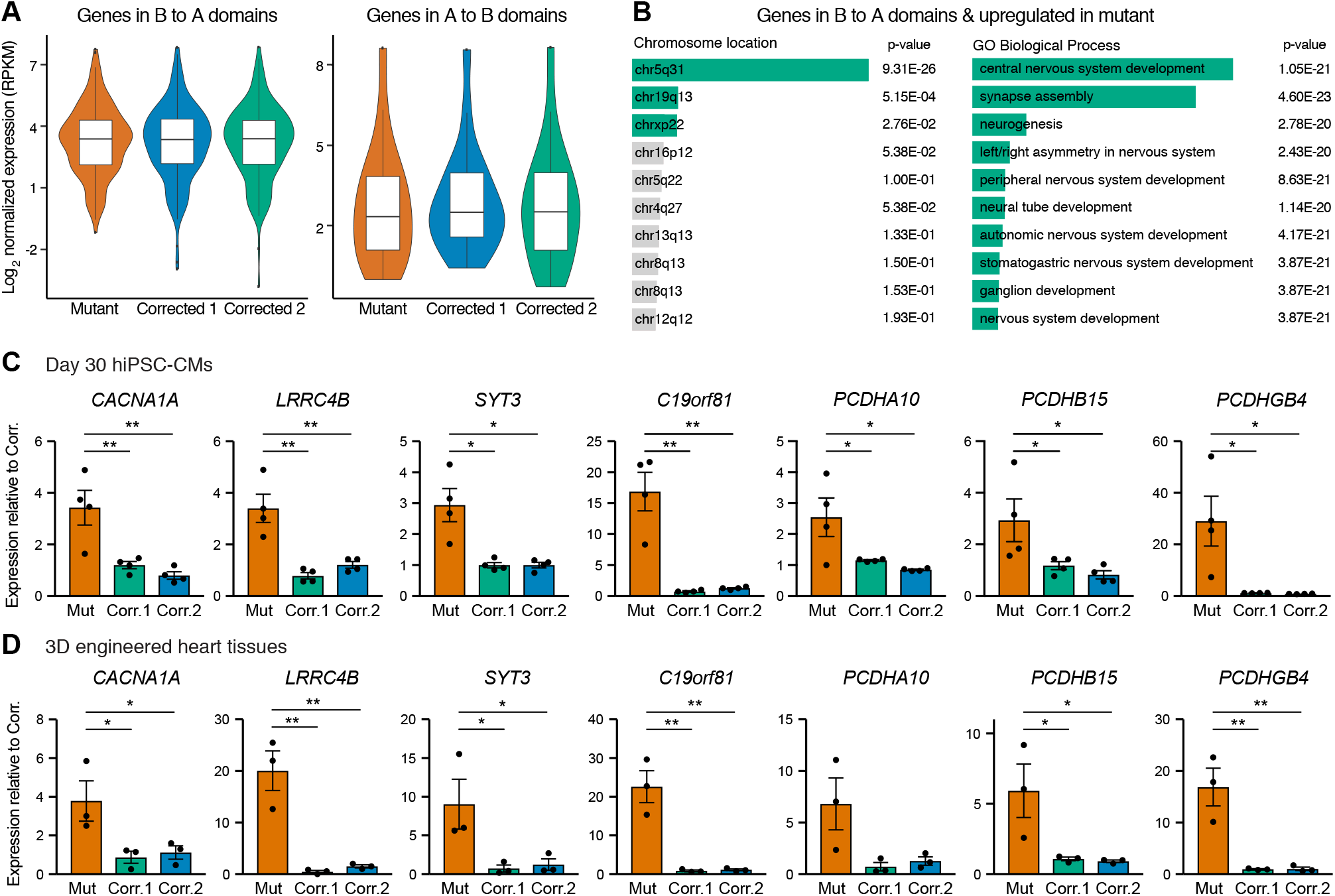
Correlation between altered chromatin compartmentalization and gene expression changes in lamin A/C haploinsufficient hiPSC-CMs. (**A**) Violin plots showing the expression of genes found with lamin A/C-sensitive domains (average expression in RNA-seq data from 3 differentiations). Boxplots indicate the first quartile, median, and third quartile, while whiskers are from the 5^th^ to 95^th^ percentile. In the left panel, note that the tail of genes expressed at very low levels in corrected hiPSC-CMs is less pronounced in mutant cells. (**B**) Selected results from ontology enrichment analyses of upregulated genes in domains aberrantly found in the A compartment in mutant hiPSC-CMs (average fold-change > 2; Fig. S3C). Term as are listed by their rank based on the combined score, and the p-values are reported (Table S5). (**C-D**) RT-qPCR validation of gene expression changes in hiPSC-CMs matured by culture *in vitro* for 30 days (C) or by generation of 3D-EHTs (D). Differences *versus* mutant were calculated by one-way ANOVA with post-hoc Holm-Sidak binary comparisons (* = p < 0.05, ** = p < 0.01; n = 4 differentiations for panel C, and n = 3 3D-EHT batches for panel D; average ± SEM).

Of note, most genes in the group just described had not been determined as differentially expressed based on RNA-seq analysis using Cufflinks since lowly-expressed genes are subject to strong negative penalization when calculating the q score due to the challenges in robustly assessing their expression (Trapnell et al., 2012, 2010). This explains their absence in the lists used for the overlap shown in Supplemental Figure 4B. To increase our confidence with these results we validated the expression of several genes within this class by RT-qPCR. Moreover, to exclude that such gene expression differences were simply explained by a differentiation delay in mutant hiPSC-CMs, we analyzed hiPSC-CMs matured either by longer 2D culture or by the generation of 3D-EHTs. Remarkably, nearly all genes tested showed consistent significant upregulation in mutant samples (Fig. 7C-D and Fig. S4D-E). These findings confirmed that impaired transition to the B compartment of selected lamin A/C-sensitive domains leads to upregulation of multiple non-cardiac genes that would otherwise be silenced during cardiomyocyte differentiation.

### Ectopic P/Q-type and potentiated L-type calcium currents contribute to electrophysiological abnormalities of lamin A/C haploinsufficient hiPSC-CMs

Genes found in lamin A/C-sensitive B domains and upregulated in mutant hiPSC-CMs included multiple factors with either unknown function, such as the putative uncharacterized protein C19orf81, or with established roles in the nervous system but not normally expressed in the heart (Fig. 7C-D and Fig. S4D-E). This group included genes from all of the three protocadherin clusters on chromosome 5 (alpha, beta, and gamma, exemplified by *PCDHA10*, *PCDHB15*, and *PCHDGB4*; Chen and Maniatis, 2013), *LRRC4B* (encoding for the postsynaptic cell adhesion molecule NGL-3; Maruo et al., 2017), *SYT3* (involved in postsynaptic endocytosis; Awasthi et al., 2018), and *CACNA1A* (which encodes for the pore-forming subunit of neuronal P/Q-type calcium channels; Rajakulendran et al., 2012).

*CACNA1A* appeared particularly interesting given the prolonged action potential duration observed in mutant hiPSC-CM populations. Indeed, the depolarizing I_Ca,P_ and I_Ca,Q_ currents resulting from the protein product of *CACNA1A* are known to be strong and long-lasting, even more so than I_Ca,L_ currents typical of hiPSC-CMs (Catterall et al., 2005; Nimmrich and Gross, 2012). Thus, we tested whether I_Ca,P_ and I_Ca,Q_ currents contributed to the electrophysiological abnormalities of mutant cells by inhibiting such currents using two structurally unrelated highly-specific inhibitors derived from spider venoms: ω-Conotoxin MVIIC and ω-Agatoxin TK (Adams et al., 1993; McDonough et al., 1996; Nimmrich and Gross, 2012). MEA experiments demonstrated that both toxins led to a modest but significant decrease in the FPDc in monolayers of mutant hiPSC-CMs, while they did not affect their depolarization amplitude (Fig. 8A-B). On the other hand, neither toxin had a significant effect on the FPDc in corrected controls (Fig. 8A-B), confirming that I_Ca,P_ and I_Ca,Q_ currents do not play a role in cardiac depolarization in physiological conditions, and establishing that the toxins had no overt non-specific effects on hiPSC-CM electrophysiology at the doses tested. These experiments indicated that ectopic expression of *CACNA1A* in mutant cells and the resulting P/Q-type calcium currents contribute to the prolonged depolarization observed in lamin A/C haploinsufficient hiPSC-CMs.

**Figure 8.**
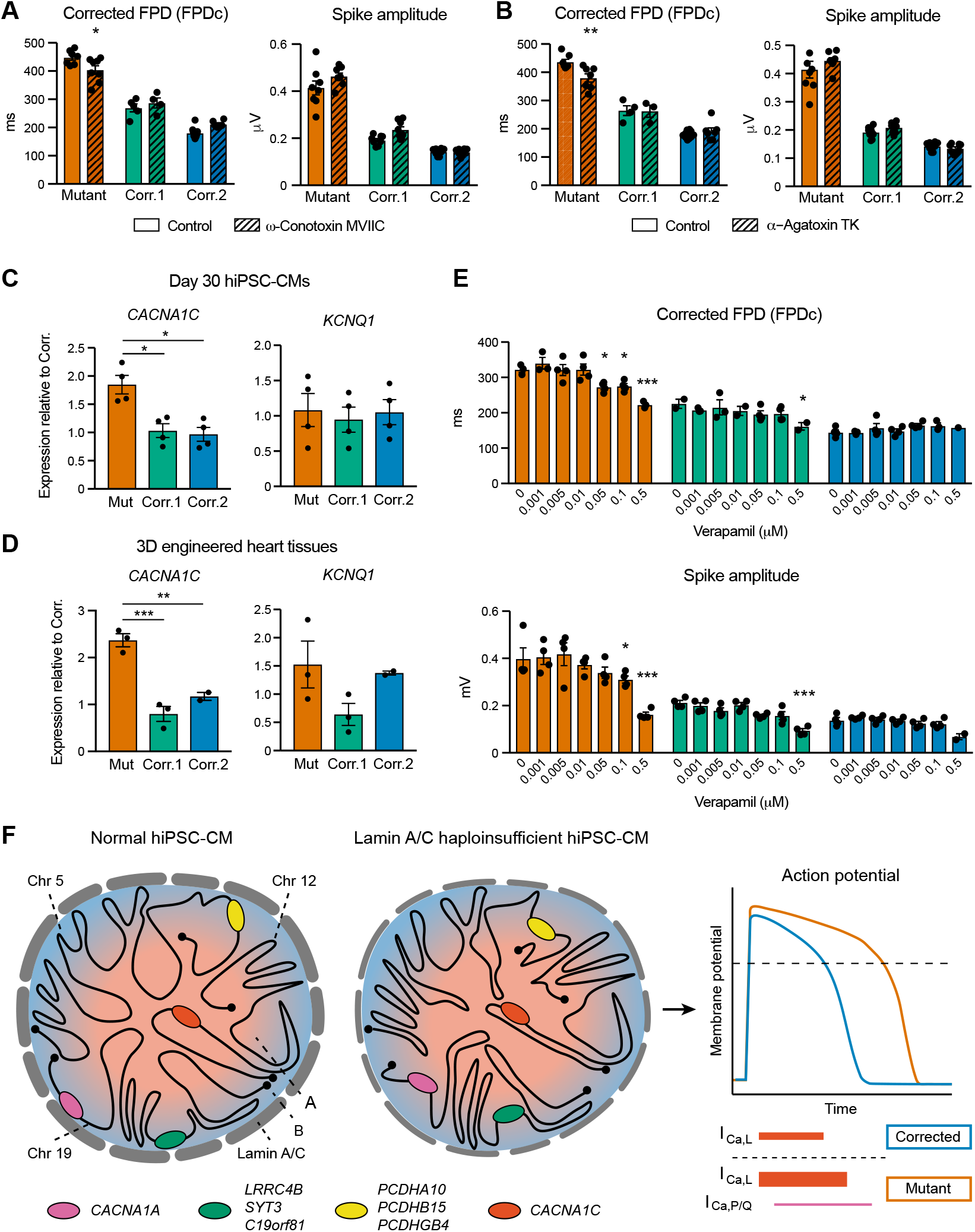
Role of P/Q-and L-type calcium currents in electrophysiological abnormalities of lamin A/C haploinsufficient hiPSC-CMs. (**A-B**) Representative quantifications of electrophysiological properties from MEA analyses. hiPSC-CMs were maintained in standard culture conditions, or pre-treated for 20 min with the indicated inhibitors for P/Q-type calcium channels (ω-Conotoxin MVIIC: 2 µM; ω-Agatoxin TK: 0.5 µM). Differences *versus* mutant were calculated by one-way ANOVA with post-hoc Holm-Sidak binary comparisons (* = p < 0.05, n = 3-8 wells; average ± SEM). (**C-D**) RT-qPCR validation of gene expression changes in hiPSC-CMs matured by culture *in vitro* for 30 days (C) or by generation of 3D-EHTs (D). Differences *versus* mutant were calculated by one-way ANOVA with post-hoc Holm-Sidak binary comparisons (* = p < 0.05, ** = p < 0.01, *** = p < 0.001; n = 4 differentiations for panel C, and n = 3 3D-EHT batches for panel D; average ± SEM). (**E**) As in panels A-B, but hiPSC cm were pre-treated for 10 min with increasing doses of the L-type calcium channel blocker verapamil. (**F**) Proposed model for the chromatin compartmentalization-dependent and independent effects of lamin A/C haploinsufficieny in developing hiPSC-CMs. Mutant cells have stronger interactions within the A compartment and decreased intermixing of A and B compartments. *Trans* interactions are globally reduced, while large chromosomes (exemplified by chromosomes 5 and 12) and small-chromosomes (exemplified by chromosome 19) self-associate more. Compartmentalization of selected genomic hotspots is dysregulated, in particular for compartments that developmentally transition from A to B in normal hiPSC-CMs but remain in A in mutant cells (exemplified by the magenta, green, and yellow loci). A large number of genes are dysregulated independently of chromatin compartment changes (exemplified by the red locus). These combined effects drive stronger L-type calcium currents (thick red line) and ectopic P/Q type calcium currents (thin magenta line), leading to prolonged action potentials, stronger calcium fluxes, systolic hyperfunction, and diastolic dysfunction.

As mentioned above, we also observed *CACNA1C* upregulation and *KCNQ1* downregulation in mutant hiPSC-CMs at day 14 of differentiation (Fig. 4A). RT-qPCR analyses in hiPSC-CMs matured for a longer period in 2D monolayers or in 3D-EHTs indicated that *KCNQ1* downregulation was specific to early hiPSC-CMs (Fig. 8C-D). On the other hand, upregulation of *CACNA1C* was maintained in more mature hiPSC-CMs (Fig. 8C-D). Of note, *CACNA1C* is a gene always found in the A chromatin compartment throughout hiPSC-CM differentiation (Fields et al., 2017), and such localization was unaltered in mutant hiPSC-CMs (Table S4). Given the established role of I_Ca,L_ in the development and maintenance of the cardiac action potential (Catterall et al., 2005), we tested if its inhibition with the L-type calcium blocker verapamil could revert the electrophysiological abnormalities of mutant hiPSC-CMs. Remarkably, low concentrations of verapamil markedly reduced both the FPDc and the spike amplitude of mutant hiPSC-CM monolayers, while they had little or no effect on corrected controls (Fig. 8E). Overall, we conclude that the combination of ectopic P/Q-type calcium currents due to aberrant chromatin compartment dynamics, and of enhanced L-type calcium currents *via* other epigenetic mechanisms lead to the electrophysiological abnormalities of lamin A/C haploinsufficient hiPSC-CMs (Fig. 8F).

## Discussion

### Disease modeling of cardiac laminopathy in developing cardiomyocytes

Previous efforts to study cardiac laminopathy using patient-derived hiPSCs have relied on hiPSCs from unrelated healthy subjects as controls (Siu et al., 2012; Lee et al., 2017), a strategy which has known limitations (Sala et al., 2017). By developing a method to generate isogenic control hiPSCs through entirely scarless gene correction we provide a more rigorous *in vitro* model of cardiac laminopathy due to lamin A/C haploinsufficiency, which we apply to broaden our understanding of the disease with respect to cardiac electrophysiology and contractility.

Because the earliest manifestation of cardiac laminopathy is severe electrical abnormalities in the myocardium (Van Rijsingen et al., 2012; Hasselberg et al., 2018; Kumar et al., 2016), our observations of aberrant electrophysiological properties in mutant hiPSC-CMs may be noteworthy. While alterations in cardiac rhythm could have been anticipated given the clinical manifestations of this disease, we were surprised to note increased FPDc and action potential duration. The FPD is the *in vitro* analogue of the QT interval measured by electrocardiogram (which indicates the interval between ventricular depolarization and repolarization). Genetic or acquired prolongation of QT interval is a strong risk factor for development of severe arrhythmias (Vandael et al., 2017), and QT interval prolongation has been reported in patients with *LMNA* mutations causing cardiac laminopathy (Pan et al., 2009), Emery-Dreifuss muscular dystrophy (Russo et al., 2012), and Hutchinson-Gilford progeria syndrome (Merideth et al., 2008). Although prolonged QT has not been widely described as a hallmark of cardiac laminopathy, we speculate that this could be an underappreciated early clinical phenotype. Prolonged QTc and subsequent increase in calcium influx might predispose patients to cardiac arrhythmias and/or could be an early marker of a broader conduction system disease.

A striking phenotype of the laminopathic cardiomyocytes was their enhanced contractility in monolayers and in engineered heart tissues. Increased contractility can be explained by the prolonged action potentials and stronger calcium fluxes that these myocytes exhibit. At a first glance, greater contractility might appear to conflict with the clinical phenotype of some laminopathy patients who develop systolic heart failure. However, in contrast to the early-onset and highly penetrant malignant conduction disease, left ventricular dilatation and systolic failure is experienced by just a fraction of cardiac laminopathy patients and only after years after the initial diagnosis (Van Berlo et al., 2005; Kumar et al., 2016; Tobita et al., 2018). Moreover, heart failure is a complex disease that involves multiple cell types as well as the extracellular matrix (Metra and Teerlink, 2017). Thus, we propose that the systolic hyperfunction and diastolic dysfunction of lamin A/C haploinsufficient hiPSC-CMs might reflect an early and cell-autonomous phenotype of cardiac laminopathy, which over the years can evolve into an organ-level disease characterized by decreased systolic function. A possible mechanism for this could be the chronic calcium overload in laminopathic cardiomyocytes, as this is a well-established stimulus that can lead to dilated cardiomyopathy due to activation of signaling pathways such as calcineurin/NFAT (Molkentin et al., 1998; Nakamura et al., 2008; Zhang et al., 2016). Additionally, it was previously shown that laminopathic cardiomyocytes are highly sensitive to apoptosis (Ho et al., 2011; Lee et al., 2017), which could over the years contribute to the development of severe cardiac fibrosis that is typical in cardiac laminopathy patients (van Tintelen et al., 2007b; Fontana et al., 2013; Tobita et al., 2018). A corollary of these intriguing hypotheses is that medical interventions during the early phases of the disease, such as the use of calcium antagonists and/or anti-apoptotic agents (Lee et al., 2014; Nie et al., 2018), may be able to prevent the development of heart failure in cardiac laminopathy patients.

### Chromatin architecture changes in lamin A/C haploinsufficient cardiomyocytes

Our study’s primary goal was to study the pathogenesis of cardiac laminopathy. Nevertheless, by integrating Hi-C results from lamin A/C haploinsufficient and control hiPSC-CMs with our previous Hi-C dataset at different stages of human cardiogenesis (Fields et al., 2017), we were also able to shed light on the physiological roles of lamin A/C during cardiac specification. We observed that lamin A/C haploinsufficiency reduces inter-chromosomal interactions, reinforces the separation between small and large chromosomes, increases the long-range segregation of A and B chromatin compartments, and strengthens short-range homotypic interactions within the A compartment. Intriguingly, most of these effects antagonize the chromatin organization dynamics that normally occur during human cardiogenesis (Fields et al., 2017). These data demonstrate that lamin A/C is an important mediator of physiological changes in nuclear architecture during differentiation, in agreement with its gradual upregulation as cells exit the pluripotent state and commit to the cardiac lineage.

The major hypothesis going into this study was that lamin A/C haploinsufficiency would induce widespread gene dysregulation due to inappropriate A/B compartmentalization, the so-called “chromatin hypothesis” for the pathogenesis of cardiac laminopathy (Worman and Courvalin, 2004; Cattin et al., 2013). This hypothesis was not robustly supported. RNA-seq analysis showed that only ~325 genes were up-or down-regulated in laminopathic cardiomyocytes. Analysis of A/B compartment changes revealed that only ~1.2% of the genome changed compartments in *LMNA* mutants, and these aberrations were concentrated in hotspots on chromosomes 5 and 19. Surprisingly, the overlap between strong dysregulation in gene expression and compartment aberrations was minimal (less than 2%; Fig. S4B). As discussed below, the *CACNA1A* gene is ectopically expressed and resides in an A compartment that fails to silence during differentiation. Thus, while examples can be found that may contribute to disease, most of the transcriptional dysregulation appears to result from factors other than errors in compartmentalization. Overall, these findings do not support the “chromatin hypothesis” for the pathogenesis of cardiac laminopathy. On the other hand, our results agree with previous findings from mouse embryonic stem cells (mESCs), in which depletion of B-type nuclear lamins results in minimal changes in A/B compartmentalization (Amendola and van Steensel, 2015; Zheng et al., 2015, 2018). Our results establish that while A-type lamins seem to participate in chromatin organization in developing cardiomyocytes, lamin A/C haploinsufficiency leads to only modest alterations in A/B compartmentalization even in mechanically active cells.

Interestingly, we identify *CACNA1A* as a disease-associated gene linked to alterations of A/B compartmentalization. *CACNA1A* is normally expressed throughout the nervous system with the highest expression on Purkinje neurons in the cerebellum and in cerebellar granular cells (Nimmrich and Gross, 2012). Its ectopic expression and the modest contribution of the resulting P/Q-type calcium currents on FPDc prolongation in lamin A/C haploinsufficient hiPSC-CMs suggests that this may represent a therapeutic target to ameliorate the electrical abnormalities in the myocardium of cardiac laminopathy patients. Further assessments of *CACNA1A* expression in patient-derived primary samples will be of high interest. We also identified the locus containing the three protocadherin clusters as another lamin A/C-sensitive B domain associated to ectopic gene expression. Protocadherin genes are well-established mediators of accurate neuronal connectivity, and are not normally expressed in the heart (Hayashi and Takeichi, 2015). It is an intriguing possibility that ectopic protocadherin expression may change cellular adhesion patterns in the cardiac conduction system, which may contribute to conduction disease in cardiac laminopathy. Other interesting targets of chromatin compartment dysregulation in lamin A/C haploinsufficient hiPSC-CMs are GPC4, whose upregulation might explain the incomplete silencing of cardiac progenitor genes (Strate et al., 2015), and CLEC11A (also known as stem cell growth factor), a cytokine normally involved in bone marrow homeostasis but upregulated in cardiac disease (Wang et al., 2013).

If chromatin compartment alterations do not directly explain the majority of gene expression changes observed in lamin A/C haploinsufficient hiPSC-CMs, what is driving these changes? An important example of ths behaviour is *CACNA1C*, which is upregulated in mutant cells while not changing compartmentalization. At least two non-exclusive possibilities come to mind. First, it remains possible that alterations in chromatin organization play a primary role, albeit not in the form of outright changes in A/B compartmentalization. For instance, it was reported that B-type lamins can indirectly affect the expression of genes within the nuclear interior by affecting the interaction between TADs as a consequence of distal alterations in LAD compaction (Zheng et al., 2018). It is thus possible that A-type lamins might have a similar role, which will be interesting to test in detail in future studies. A second possibility for compartment-independent gene expression changes might be the result of changes in intracellular signaling pathways such as MAPK and mTOR. Indeed, such pathways have well-established links with the nuclear lamina (Dobrzynska et al., 2016), and are upregulated in animal models of cardiac laminopathy (Muchir et al., 2007, 2012; Choi et al., 2012).

Emerging evidence suggests that nonsense/haploinsufficency mutations in *LMNA* may have a different pathogenesis than missense mutations. We have recently collaborated to study a previously described heterozygous K219T missense mutation in *LMNA* (Roncarati et al., 2013). Interestingly, this mutation leads to distinct electrophysiological abnormalities, namely reduced peak sodium current and diminished conduction velocity, which are caused by downregulation of *SCN5A* as result of closer proximity to the nuclear lamina and increased H3K27me3 (Salvarani, Crasto et al., manuscript in preparation). In the current study we have not observed a similar reduction of *SCN5A* expression in lamin A/C haploinsufficient hiPSC-CMs (Fig. S2D). Furthermore, our Hi-C data indicate that the *SCN5A* gene is found in a chromatin domain which is always part of the A compartment both throughout normal cardiac differentiation (Fields et al., 2017) and in lamin A/C haploinsufficient hiPSC-CMs (Table S4). These observations prompt the intriguing hypothesis that haploinsufficient and missense mutations in *LMNA* might lead to cardiac laminopathy *via* distinct molecular mechanisms.

In conclusion, our work establishes that while lamin A/C haploinsufficient hiPSC-CMs show marked alterations in electrophysiology, contractility, and chromosomal topology, phenotypic changes cannot, for the most part, be directly explained by alterations in chromatin compartmentalization. With this in mind, it is important to mention that this study does not come without limitations. We acknowledge that modeling diseases of the adult heart in immature hiPSC-CMs suffers from inherent drawbacks. It is possible that functional chromatin dysregulation could be more important in adult myocytes subjected to high levels of mechanical stress *in vivo*. Addressing this aspect will require substantial advances in our ability to mature hiPSC-CMs and/or improvements of genome-wide chromatin conformation capture technologies in order to reliably measure chromatin architecture from small numbers of myocytes isolated from precious primary samples. Furthermore, future studies will be required to test whether other types of mutations (such as missense changes like the aforementioned K219T mutation) result in more substantial genome-wide alterations in chromatin topology than what was observed following lamin A/C haploinsufficiency. All considered, our work provides a stepping stone towards understanding the relevance of the “chromatin hypothesis” in the pathogenesis of cardiac laminopathy, while it also sheds light on the physiological roles of A-type lamins in the chromatin dynamics that accompany human cardiogenesis.

## Materials and methods

### hiPSC culture and differentiation

Human induced pluripotent stem cells (hiPSCs) were cultured and differentiated into cardiomyocytes (hiPSC-CMs) with minor modifications of previously described methods (Burridge et al., 2014; Fields et al., 2017). hiPSCs were cultured on plates pre-coated with 0.5 μg/cm^2^ recombinant human Laminin-521 matrix (rhLaminin521; Biolamina) with daily changes of antibiotic-free Essential 8 (E8) media (ThermoFisher). Cells were passaged as small clumps with Versene (ThermoFisher), and 10 μM Y-27632 (Tocris) was added for the first 16 hr.

For hiPSC-CM generation, cells were passaged as single cells with Versene, and seeded at a density of 2.5 × 10^5^ per well of a 12-well plate pre-coated with 2 μg/cm^2^ rhLaminin521 (denoted day −2). After 24 hr media was changed to E8 with 1 μM CHIR-99021 (Cayman), denoted day −1. On day 0 media was changed to RBA media (RPMI 1640 with glutamine [ThermoFisher] supplemented with 500 μg/ml BSA and 213 μg/ml ascorbic acid [both from Sigma-Aldrich]) supplemented with 4 μM CHIR-99021. At day 2, media was changed to RBA with 2 μM WNT-C59 (Selleckchem). On day 4, media was changed to RBA. On day 6, media was changed to RPMI-B27 media (RPMI with 1x B-27 supplement, both from Thermo Fisher), with further media changes every other day. Beating was first observed between day 7 and day 9, and cells were cultured until day 14 before collection (unless otherwise indicated). hiPSC-CMs to be used for functional assays of cardiac electrophysiology or contractility were preconditioned with a 30 min heat-shock at 42º on day 13, and cryopreserved at day 14 following single-cell dissociation using 0.25% w/vol Trypsin (ThermoFisher) in Versene.

Frozen hiPSC-CM stocks were thawed and seeded at a density of 2 × 10^5^ cells/cm^2^ onto rhLamin521 pre-coated dishes (2 μg/ml) in RPMI-B27, which was supplemented with 10 μM Y-27632 and 5% FBS (ThermoFisher) for the first 16 hr. hiPSC-CMs were then cultured in RPMI-B27 with media changes every other day. After one week (day 21 of differentiation), hiPSC-CMs were dissociated to single cell using 0.05% Trypsin-EDTA (ThermoFisher), and seeded at the desired density the downstream assays.

### Gene editing

hiPSCs with a heterozygous c.672C>T mutation in the *LMNA* gene (resulting in p.Arg225*, or R225X) were generated through lentiviral reprogramming of dermal fibroblasts from a 56 year-old male patient with severe cardiac laminopathy (Siu et al., 2012). Cells were obtained at passage 29, adapted to culture in E8/rhLaminin521, and banked at passage 34. These hiPSC stocks were confirmed to be Mycoplasma negative (MycoAlert Detection Kit, Lonza), and proved euploid by conventional G-banding karyotyping (Diagnostic Cytogenetics Inc, Seattle). Cell identity was tested by Sanger sequencing of a genomic PCR product for exon 4 of *LMNA* (PCR primers: 5’-GGCTGGGTGATGACAGACTT-3’ and 5’-TACTGCTCCACCTGGTCCTC-3’; sequencing primer 5’-GCCCTAGTGGACAGGGAGTT-3’), which confirmed the expected c.672C>T heterozygous mutation.

The mutation was corrected into the wild-type allele by adapting a previously described two-step method for scarless genome editing relying on CRISPR/Cas9-facilitated homologous recombination of a targeting vector containing the wild-type allele in one homology arm, and an excisable piggyBac drug resistance cassette (Fig. S1A; Yusa, 2013; Yusa et al., 2011). Since the c.672C>T mutation lies close to the 3’ splice acceptor site of exon 4 (Fig. S1B), we reasoned that any intergenic mutation could have poorly predictable effects on *LMNA* splicing. To avoid any kind of genomic scar, we identified a suitable endogenous “TTAA” site in the third intron of *LMNA* and located 151 bp upstream to the c.672C>T mutation in exon 4 (Fig. S1B). Further, we designed single guide RNAs (sgRNAs) spanning such TTAA site, so that only the endogenous allele could be cut by CRISPR/Cas9. This strategy allowed us to avoid inserting any additional mutation onto one of homology arms in the targeting vector (such as those classically used to disrupt the PAM site). Two sgRNAs were designed using mit.crispr.edu, and had a score higher than 75%, indicating a very high *in-silico* predicted specificity (sgRNA 1: 5’-CTACCAGCCCCACTTTAACC-’3 and sgRNA2 5’-TCAGCTCCCAGGTTAAAGTG-3’, sequences without PAM site). To further decrease the risk of CRISPR/Cas9 off-target activity, we adopted the enhanced specificity *Streptococcus Pyogenes* Cas9 (eSpCas9) developed by Dr. Feng Zhang and colleagues (Slaymaker et al., 2015). The sgRNA was cloned into the eSpCas9(1.1) plasmid (Addgene #71784) using a standard method based on restriction digestion with BbsI followed by ligation of a double-stranded oligo (Ran et al., 2013). The resulting plasmids were named eSpCas9(1.1)_LMNA_sgRNA1 and eSpCas9(1.1)_LMNA_sgRNA2. The sequences were confirmed by Sanger sequencing, and the sgRNAs were validated to have a high on-target activity as measured by T7E1 assay in HEK293 cells (which was comparable to that observed using wild-type SpCas9).

The *LMNA* targeting vector was constructed starting from the MV-PGK-Puro-TK_SGK-005 plasmid (Transposagen), which contains a piggyBac transposon encoding for a PGK-EM7 promoter-driven dual positive/negative selection cassette (puromycin N-acetlytransferase, ensuring resistance to puromycin, and truncated *Herpes simplex* virus thymidine kinase, conferring sensitivity to ganciclovir or its analog fialuridine). First, the piggyBac cassette was excised using NsiI and BsiWI and isolated. Then, a backbone with ends suitable for the subsequent overlap-based assembly was obtained from this same plasmid after removal of the piggyBac cassette using NotI and AscI. Finally, these two fragments were re-assembled together with two PCR products representing the 5’ and 3’ homology arms to the *LMNA* gene. The two homology arms were approximately 1 Kb long, and were amplified from genomic DNA of RUES2 human embryonic stem cells (hESCs) using primers: 5’-GGTCCCGGCATCCGATACCCAATG-GCGCGCCCGTACTTCAGGCTTCAGCAGT-3’ and 5’-AAAGAGAGAGCAATATTTCAAGAATGCATG-CGTCAATTTTACGCAGACTATCTTTCTAGGGTTAACCTGGGAGCTGAGTGC-3’ (for the 5’ homology arm); 5’-AATTTTACGCATGATTATCTTTAACGTACGTCACAATATGATTATCTTTCTAGGGTTAAAGT-GGGGCTGGTAGTG-3’ and 5’-CGAATGCGTCGAGATATTGGGTCGCGGCCGCCCTGTCACAAATAG-CACAGCC-3’ (for the 3’ homology arm), and Q5 High-Fidelity DNA Polymerase (New England Biolabs) according to the manufacturer’s instructions. The four-way assembly reaction was performed using NEBuilder HiFi DNA Assembly Kit (New England Biolabs) according to the manufacturer’s instructions, and the resulting targeting plasmid was named pbLMNA_R225R. Sanger sequencing confirmed that the 3’ homology arm contained the wild-type R225R allele, while the remaining genomic sequence of both homology arms was identical to that of the R225X hiPSC line as no SNPs were identified. The cloning strategy was designed so that during PCR the “TTAA” site was inserted both at the end of the 5’ homology arm and at the start of the 3’ homology arm, ensuring that the piggyBac cassette contained within could be excised using transposase while leaving behind a single “TTAA” matching the original genomic sequence (Fig. S1A).

For the first gene targeting step, 7.5 × 10^4^ hiPSCs were seeded in each well of 6-well plate and immediately transfected using GeneJuice (Millipore) according to the manufacturer’s instructions. Briefly, for each well 3 µl of GeneJuice was mixed with 100 µl of Opti-MEM (ThermoFisher Scientific) and incubated for 5 min at room temperature. 1 µg of DNA was added to the transfection solution (equally divided between pbLMNA_R225R and either eSpCas9(1.1)_LMNA_sgRNA1 or eSpCas9(1.1)_LMNA_sgRNA2), which was further incubated for 15 min at room temperature and finally added to the cell suspension. After 16 hr from transfection, cells were washed with DPBS and cultured for another 3 days. Gene targeted cells were selected by adding 1 µg/ml puromycin to the media for 4 days, after which the dose was reduced to 0.5 µg/ml. 10 µM Y-27362 was added for the first 48 hr of selection. Puromycin was then maintained at all times until the second gene targeting step to prevent silencing of the piggyBac transgene. After 7 days from the transfection, 10-15 individual and well-separated colonies could be identified in each well of 6-well plate, indicating that they likely arose from clonal expansion of a single gene-edited hiPSC. Colonies were manually picked following gentle treatment with Versene to facilitate their detachment from the matrix, and individually expanded as individual lines. Clones were screened by genomic PCR using LongAmp Taq Polymerase (New England Biolabs) according to manufacturer’s instructions, except that all reactions were performed using an annealing temperature of 63 °C and an extension time of 2 min. The primer sequences are reported in Supplemental Table 6, and the genotyping strategies are illustrated in Supplemental Figure 1. Briefly, junctional PCRs for both the 5’ and 3’ integration site (5’-and 3’-INT) were used to confirm site-specific integration, while locus PCRs were used to monitor the presence of residual wild-type alleles. This allowed to discriminate homozygous clones from heterozygous ones or mixed cell populations. Finally, PCRs of the targeting vector backbone (5’-and 3’-BB) were performed to exclude random integration of the plasmid elsewhere in the genome. Homozygous clones with only on-target integration events were selected (3 out of 18 and 2 out of 24 for sgRNA 1 and sgRNA 2, respectively). These positive clones were further characterized by Sanger sequencing of the 5’-and 3’-INT PCR products to confirm the presence of the wild-type R225R allele in homozygosity (found in all of the 5 lines) and exclude other unwanted mutations elsewhere in the locus (absent in all 5 lines). Two clones (one for each sgRNA) were karyotyped by standard G-banding, which confirmed their euploidy, and were therefore selected for the second gene targeting step. These clones were named pb R225R g1 and pb R225R g2 (Fig. S1C).

To remove the piggyBac and restore the *LMNA* locus to its original form, pb R225R g1 and pb R225R g2 hiPSCs were transfected as described above but using 1 µg of excision-only piggyBac transposase expression vector (PBx; Transposagen). Puromycin was removed from the media the day before transfection, and subsequently omitted. After 3 days from the transfection, the populations were passaged as single cells, and 1 × 10^4^ cells were seeded per 10 cm plate in the presence of 10 µM Y-27362. On the next day, negative selection of cells still possessing the piggyBac cassette was initiated by adding 200 nM fialuridine. 10 µM Y-27362 was added for the subsequent 48 hr. Selection was complete after 5 days, at which point 10-50 individual and well-separated colonies could be identified in each 10 cm dish. Individual colonies were isolated, clonally expanded, and screened by genomic PCR as described above to identify those with homozygous reconstitution of the wild-type allele (5 out of 30 and 6 out of 39 for sgRNA 1 and sgRNA2, respectively). These were further characterized by sequencing to ensure that the sequence surrounding the “TTAA” site was faithfully reconstituted upon piggyBac excision (confirmed in a subset of 4 lines, 2 for each sgRNA; Fig. S1C-D). Two clones (one for each sgRNA), were karyotyped by standard G-banding, which confirmed their euploidy, and therefore selected for subsequent functional experiments. These clones were named R225R g1-15 and R225R g1-38, and are referred to in the text and figures as Corrected 1 and Corrected 2 (or Corr.1 and Corr.2).

Parental R225X hiPSCs (referred to in the text and figures as Mutant or Mut) were cultured in parallel throughout the whole gene editing procedure to provide a passage-matched control, and were re-banked at passage 49 together with Corrected 1 and Corrected 2. These cells were confirmed to be euploid by G-banding karyotyping. Mutant and Corrected hiPSCs to be used for derivation of hiPSC-CMs were cultured between passage 50 to 60 before resorting to a new frozen stock.

### MEA

Multi-electrode array (MEA) analyses were performed on hiPSC-CM monolayers at day 30 of differentiation. hiPSC-CMs at day 21 of differentiation were seeded at a density of 5 × 10^4^ cells per well of 48-well MEA plates (CytoView MEA 48; Axion Byosystems) pre-coated with rhLaminin521 (2 μg/ml). Cells were cultured in RPMI-B27 with media changes every other day. After 9 days (day 30 of differentiation), cells were prepared for MEA analysis by changing the culture media to Tyrode’s buffer (140 mM NaCl, 5.4 mM KCl, 1.8 mM CaCl2, 1 mM MgCl2, 0.33 mM NaH_2_PO_4_, 5 mM D-glucose, and 10 mM HEPES; pH adjusted to 7.36) pre-warmed at 37 ºC. After 10 min of equilibration in Tyrode’s buffer at 37 ºC, MEA data were acquired for 5 min using the Maestro MEA system (Axion Biosystems) using standard recording settings for spontaneous cardiac field potentials. Data acquisition and automated data analysis was performed using Axis software, version 2.1. Standard acquisition settings have 130 × gain, and record from 1 to 25 000 Hz, with a low-pass digital filter of 2 kHz for noise reduction. Automated data analysis was focused on the 30 most stable beats within the recording period. The beat detection threshold was 100 µV, and the field potential duration (FPD) detection utilized an inflection search algorithm with the threshold set at 1.5 × noise to detect the T wave. The FPD was corrected for the beat period (FPDc) according to the Fridericia’s formula: FPDc = FPD / (beat period)^1/3^ (Rast et al., 2016; Asakura et al., 2015). Reported results for individual wells were calculated by averaging all of the electrodes. In certain instances, poor signal quality and/or irregularity of field potential behavior prevented the calculation of certain parameters (such as FPD). The presented data constitutes all recorded values that could be reliably measured by the software based on automatic quality-control thresholds.

For pharmacological studies of P/Q-and L-type calcium current inhibition, Tyrode’s buffer was supplemented with 2 µM ω-Conotoxin MVIIC, 0.5 µM ω-Agatoxin TK, or 0.001-0.5 µM verapamil (all from Tocris). hiPSC-CMs were incubated at 37 ºC for 20 min or 10 min (for ω-Conotoxin and ω-Agatoxin, or verapamil, respectively) before MEA data acquisition.

### Whole cell patch clamp

Whole-cell patch clamp recordings were obtained from individual hiPSC-CMs at day 30 of differentiation. hiPSC-CMs at day 21 of differentiation were seeded at a density of 4.5 × 10^5^ cells per well of 35-mm glass-bottom FluoroDish with nanopatterned surfaces pre-coated with rhLaminin521 (2 μg/ml). Anisotropically nanofabricated substrata (ANFS) with 800 nm topographic features were fabricated via UV-assisted capillary force lithography as previously described (Macadangdang et al., 2015). First, liquid polyurethane acrylate (PUA) prepolymer was drop dispensed onto a silicon master mold. A transparent polyester film (PET) was then placed on top of the dispensed PUA. After exposure to UV radiation (λ = 250–400 nm), the film was peeled away from the silicon master, creating a PUA mold. A polyurethane-based prepolymer (NOA76, Norland Products Inc.) was then drop dispensed onto standard glass coverslips and the PUA mold was placed on top. The mold was then exposed to UV radiation for curing. After curing, the PUA mold was peeled off, leaving behind an ANFS for cell culture. Dishes were sterilized and activated by gas plasma treatment before coating with rhLaminin521. After 9 days, cells were assayed by whole-cell patch clamp on the 37 °C heated stage of an inverted DIC microscope (Nikon) connected to an EPC10 patch clamp amplifier and computer running Patchmaster software version 2×73.2 (HEKA). Cells on patterned coverslips were loaded onto the stage and bathed in Tyrode’s buffer. An intracellular recording solution (120 mM L-aspartic acid, 20 mM KCl, 5 mM NaCl, 1 mM MgCl2, 3 mM Mg2+-ATP, 5 mM EGTA, and 10 mM HEPES) was used in conjunction with borosilicate glass patch pipettes (World Precision Instruments) with a resistance in the range of 2–6 MΩ. Offset potentials were nulled before formation of a gigaΩ seal and fast and slow capacitance was compensated for in all recordings. Membrane potentials were corrected by subtraction of a 15 mV liquid junction potential calculated by the HEKA software. Current injection was controlled by the software and used to hold patched cells at an artificial resting membrane potential of −70 mV. Cells that required more than 100 pA of current to achieve a −70 mV resting membrane potential were excluded from analysis as excessive application of current was taken as indication of poor patch quality and/or membrane integrity. To generate a single action potential, a 5 ms depolarizing current pulse of 50 nA was then applied and the resulting voltage change recorded in current clamp mode. Action potential rise times were calculated as the time taken to reach 90% maximum action potential amplitude from 10% of the maximum amplitude. The exponential time constant (τ) was calculated from 90% to 10% repolarization of the action potential. Action potential duration was calculated as the time delay between 10% of the maximum depolarization and 90% repolarization from the maximum action potential amplitude. Gap-free recordings of spontaneous cardiomyocyte activity were then collected for 30 seconds with 0 pA current injection to provide a measure of the maximum diastolic membrane potential held by the cell without current input.

### Assessment of intracellular calcium fluxes

Calcium fluxes were assessed in hiPSC-CM monolayers at day 30 of differentiation. hiPSC-CMs at day 21 of differentiation were seeded at a density of 5 × 10^5^ cells per well of 6-well plate pre-coated with rhLaminin521 (2 μg/ml). After 9 days, cells were prepared for imaging by incubation for 30 min at 37 ºC with 1 μM Fluro-4, AM (ThermoFisher) diluted in culture media. Cells were rinsed in fresh media for 30 min at 37 ºC, and equilibrated in Tyrode’s buffer pre-warmed at 37 ºC for 10 min. hiPSC-CMs were paced at 1 Hz using a C-Dish for 6-well plate connected to a C-Pace EM cell stimulator (both from IonOptix) providing biphasic field stimulation (pulses of 10 V/cm for 20 ms). Videos of Fluo-4 fluorescence (excitation/emission of 494 and 516) were recorded at 20 frames per second (FPS) for at least 5 contractions using a Nikon Ti-E epi-fluorescent microscope with a 20x-objective and 1x coupler between the microscope and Hamamatsu flash V3 camera. Videos were obtained for at least 20 random fields of view. A custom Matlab program was used to define the region of interest (ROI, containing an individual hiPSC-CM), threshold the Fluo-4 intensity based on the surrounding non-fluorescent background, and track the average ROI_Fluo-4_ fluorescence (F) over time. The relaxation time constant, τ, was determined by fitting the formula F(t) = A*e*^-t/τ^ + B to the decay phase of the Fluo-4 transient profile, where t is time in seconds and A and B are fitted constants.

### CCQ analysis of cardiac contractility in hiPSC-CM monolayers

Contraction correlation quantification (CCQ) analysis was performed on hiPSC-CM monolayers at day 30 of differentiation. hiPSC-CMs at day 21 of differentiation were seeded at a density of 1 × 10^6^ cells per well of 6-well plate pre-coated with rhLaminin521 (2 μg/ml). After 9 days, cells were paced at 1 Hz as just described for the measurement of calcium fluxes. Bright-field videos of at least ten contractions in multiple random field of views were recorded at 30 fps using a Nikon TS100 microscope with a 20x-objective and 1x coupler between the microscope and a Canon VIXIA HF S20 camera. Videos were analyzed by CCQ using a custom Matlab script, as previously described (Macadangdang et al., 2015). Briefly, this method utilizes particle image velocimetry and digital image correlation algorithms to provide relevant contractile endpoints from bright field video recordings. A reference video frame is divided into a grid of windows of a set size. Each window is run through a correlation scheme with a second frame, providing the new location for that window in the second frame. This displacement is converted into a vector map, which provides contraction angles and, when spatially averaged, contraction magnitudes. The correlation equation used provides a Gaussian correlation peak with a probabilistic nature that provides sub-pixel accuracy.

### Generation and biomechanical characterization of 3D-EHTs

3D engineered heart tissues (3D-EHTs) were generated and characterized with minor changes to a previously described method (Leonard et al., 2018). Racks of polydimethylsiloxane (PDMS) posts were fabricated by pouring uncured PDMS (Sylgard 184 mixed at a 1:10 curing agent to base ratio) into a custom acrylic mold. Glass capillary tubes (1.1 mm diameter; Drummond) were cut to length and inserted into the holes on one side of the mold before curing to render one post in each pair rigid. Post racks were baked overnight at 65 °C before being peeled from the molds. Racks consisted of six pairs of posts that were evenly spaced to fit along one row of a standard 24-well plate. Fabricated posts were 12.5 mm long and 1.5 mm in diameter with a cap structure (2.0 mm diameter for the topmost 0.5 mm) to aide in the attachment of 3D-EHTs. The center-to-center post spacing (corresponding to pre-compacted 3D-EHT length) was 8 mm. Prior to casting 3D-EHTs, all 3D printed parts and PDMS posts were sterilized in a UVO Cleaner (Jetlight, No. 342) for 7 min, submerged in 70% ethanol, and rinsed with sterile deionized water. Rectangular 2% w/vol agarose/PBS casting troughs (12 mm in length, 4 mm in width, and ~4 mm in depth) were generated in the bottom of 24-well plates by using custom 3D printed spacers (12 mm × 4 mm in cross section and 13 mm long) as negative molds. PDMS posts racks were positioned upside down with one rigid-flexible post pair centered in each trough (leaving a 0.5 mm gap between the tip of the post and the bottom of the casting trough). Each tissue consisted of a 97 μL fibrinogen-media solution (89 μL of RPMI-B-27; 5.5 μL of 2X DMEM with 20% FBS; and 2.5 μL of 200 mg/mL bovine fibrinogen, Sigma-Aldrich) containing 5 × 10^5^ hiPSC-CMs and 5 × 10^4^ supporting HS27a human bone marrow stromal cells (ATCC), which was chilled and mixed with 3 μL of cold thrombin (at 100 U/mL, Sigma-Aldrich) just before pipetting into the agarose casting troughs. The 3D-EHT mixtures were incubated for 90 min at 37 °C, at which point the fibrin gels were sufficiently polymerized around the posts to be lubricated in media and transferred from the casting troughs into a 24-well plate with fresh 3D-EHT media (RPMI-B-27 with penicillin/streptomycin, and 5 mg/mL aminocaproic acid, Sigma-Aldrich). 3D-EHTs were supplied with 2.5 mL/well of fresh 3D-EHT media three times per week.

*In situ* force measurements were performed after 4 weeks from 3D-EHT casting. In order to pace 3D-EHTs, post racks were transferred to a custom-built 24-well plate with carbon electrodes connected through an electrical stimulator (Astro Med Grass Stimulator, Model S88X) to provide biphasic field stimulation (5 V/cm for 20 ms durations) during imaging (Leonard et al., 2018). 3D-EHTs were equilibrated in Tyrode’s buffer (containing 1.8 mM Ca^2+^) pre-heated to 37 °C and paced at 1 Hz, which was greater than the average spontaneous twitch frequency of the tissues. Videos of at least ten contractions were recorded inside a 37 °C heated chamber using a monochrome CMOS camera (Mightex, SMN-B050-U) with a board lens (The Imaging Source, TBL 8.4-2 5MP). The camera-lens configuration allowed for a capture rate of 65 frames per second (FPS) with 8.3 μm/pixel resolution and a field of view of 1536 × 400 pixels, which was sufficient to capture images of the whole 3D-EHT from rigid to flexible post. A custom Matlab program was used to threshold the images and track the centroid of the flexible post relative to the centroid of the rigid post. The twitch force profile, F_twitch_(t) = k_post_*Δ_post_(t), was calculated from the bending stiffness, k_post_, and deflection of the flexible post, Δ_post_, at all time points (t), where k_post_ = 0.95 μN/μm was determined from beam bending theory using the dimensions of the posts and taking the Young’s modulus of PDMS to be 2.5 MPa (Sniadecki and Chen, 2007). The twitch force and twitch kinetics were calculated from the twitch force profiles using a custom Matlab program.

### RT-qPCR

Quantitative reverse-transcription PCR (RT-qPCR) was performed as previously described for 2D cell monolayers (Fields et al., 2017) or 3D-EHTs (Leonard et al., 2018). 2D monolayers were lysed in RLT buffer supplemented with 1% 2-Mercaptoethanol before RNA purification using the RNeasy Mini Kit (QIAGEN) according to the manufacturer’s instructions and including the on-column DNase digestion step. cDNA was synthetized by reverse transcription of 500 ng of total RNA using M-MLV RT (Invitrogen) and random hexamer priming according to the manufacturer’s protocol and including RNase OUT (Invitrogen). RT-qPCR was performed in technical duplicate with SYBR Select Master Mix (Applied Biosystems) using 10 ng of cDNA and 400 nM forward and reverse primers. Reactions were run for 40 cycles on a 7900HT Fast Real-Time PCR System (Applied Biosystem, 4329001), all according to the manufacturer’s instructions, Gene expression relative to the housekeeping gene *RPLP0* was calculated using the ∆Ct method as 2^(Ct gene - Ct housekeeping)^. Where indicated, gene expression was further normalized to a control condition (which was set at the arbitrary value of 1). Primers were designed using PrimerBlast, and confirmed to amplify a single product. A complete list can be found in Supplemental Table 7.

For 3D-EHTs, RNA was extracted using RNeasy Plus Micro Kit (QIAGEN) according to the manufacturer’s instructions except for the following modifications. Individual 3D-EHTs were pre-digested using 2 mg/ml Proteinase K in 100 µl DPBS for 10 min at 56 °C in agitation. Cells were then lysed by adding 350 µl of RLT Plus Buffer supplemented with 1% 2-Mercaptoethanol, and cleared through the gDNA Eliminator Mini Spin Column. The RNA lysate was finally prepared for binding to the RNeasy MinElute Spin Column by adding 250 µl of 200 proof ethanol. All subsequent steps were performed according to the supplier’s recommendations. 10 µl of eluted RNA (corresponding to 50-100 ng) were subjected to reverse transcription, and 2 ng of cDNA were used as template for RT-qPCR, all as just described for 2D monolayers.

### Western blot

Protein lysates were obtained using ice-cold 1X RIPA buffer containing protease and phosphatase inhibitors (Cell Signaling 9806), and freshly supplemented with 1 mM phenylmethylsulfonyl fluoride (PMSF). After incubation for 30 min on ice, the lysate was clarified from insoluble material by centrifugation at 16,000 g for 10 minutes at 4 °C. The protein concentration was assessed using the Pierce BCA Protein Assay Kit (ThermoFisher) according to the manufacturer’s instructions. After addition of Laemmli Sample Buffer (Bio-Rad) to a final concentration of 1x, and 2-mercaptoethanol to a final concentration of 2.5%, the samples were denatured by heating at 95 °C for 5 minutes. For electrophoretic separation, 20 µg of protein for each sample was loaded onto 7.5% Mini-PROTEAN TGX Precast Protein Gels (Bio-Rad) and run at 100V for 60 minutes using 1x Tris/Glycine/SDS running buffer (Bio-Rad). Proteins were transferred onto Immobilion-P PVDF membranes (Millipore Sigma) by means of tank blotting in 1X Tris/Glycine (Bio-Rad) supplemented with 20% methanol; transfer was performed at 100 V for 60 minutes at 4 °C. Membranes were blocked in PBS supplemented with 0.1% Tween-20 (hereafter PBST) and 4% Blotting-Grade Blocker (Bio-Rad) for 1 hr at room temperature. Primary antibody incubation of whole membranes was performed overnight at 4 °C under agitation, and antibodies were diluted in the same blocking buffer. The following antibodies were used: goat polyclonal anti-lamin A/C (Santa Cruz sc-6215), 1:500 dilution; mouse monoclonal anti-TNNI (clone OTI8H8, Novus Biologicals NBP2-46170), dilution 1:500; goat polyclonal anti-NKX2-5 (R&D Systems AF2444), dilution 1:500; goat polyclonal anti-TBXT (R&D Systems AF2085), dilution 1:500; rabbit polyclonal anti-POU5F1 (Abcam 19857), dilution 1:1000; mouse monoclonal anti-GAPDH (clone 6C5, Abcam ab8245), dilution 1:2000. Membranes were washed three times in PBST for 10 minutes at room temperature, incubated for 1 hr at room temperature with species-appropriate HRP-conjugated secondary antibodies (ThermoFisher) diluted 1:10,000 in blocking buffer, and washed three times in PBST for 10 minutes at room temperature. Chemiluminescent reaction was initiated by incubation with SuperSignal West Pico PLUS Chemiluminescent Substrate (ThermoFisher), and images were acquired using a ChemiDoc Imaging System (Bio-Rad) in “high resolution” mode. Before re-probing with a new antibody, membranes were treated with Restore Plus western blot stripping buffer (ThermoFisher), washed three times, and re-blocked. Densitometric quantification of Western blots was performed using ImageJ, and protein abundance estimation was normalized on the levels of GAPDH within each lysate.

### RNA-seq

RNA sequencing (RNA-seq) was performed on 2-3 × 10^6^ hiPSC-CMs at day 14 of differentiation on three biological replicates (independent differentiations) per cell line. RNA-seq libraries were prepared from 100 ng of total RNA (obtained as described above for RT-qPCR) using the TruSeq Stranded Total RNA LT Kit with Ribo-Zero H/M/R (all Illumina), according to the manufacturer’s instructions. The analysis RNA-seq libraries were paired-end sequenced on a Illumina NextSeq 500 in a high output run with 150 cycles (75 for each end), achieving approximately 40 million paired-end reads per sample. Reads were mapped to hg38 using STAR (Dobin et al., 2013), and then quantified and processed through the Cufflinks suite (Trapnell et al., 2012), all using default parameters. Differential expressed genes exhibited a q-value < 0.05 for a pairwise comparison and were expressed at least 1 RPKM (read kilobase per million mapped reads) in one time point. Hierarchical clustering was performed with the CummeRbund suite in R. Ontology enrichment analysis was done using EnrichR (Chen et al., 2013). For comparision with previously described RNA-seq data of hESC-CM differentiation (Fields et al., 2017), all datasets were collectively re-normalized and -quantified using Cufflinks. Principal component analysis (PCA) was performed using the R function “prcomp”.

### *In situ* DNase Hi-C

Genome-wide chromosome conformation capture based on *in situ* proximity ligation of DNase I-digested nuclei (DNase Hi-C) was performed with minor modifications of a previously described method (Ramani et al., 2016; Fields et al., 2017). The assay was performed on ~2 × 10^6^ hiPSC-CMs at day 14 of differentiation on two biological replicates (independent differentiations) per cell line. Unless otherwise indicated, all molecular biology reagents were from ThermoFisher and reactions were performed at room temperature (RT). Cells were fixed in the dish with fresh RPMI-1640 supplemented with 2% formaldehyde while in gentle orbital rotation for 10 min, and subsequently quenched with 25 mM Glycine for 5 min at room temperature followed by 15 min at 4 °C. Cells were then treated with 0.05% Trypsin for 10 min at 37 °C, rinsed in RPMI-1640 with 10% FBS, and scraped off the plate. Cells were washed once with PBS, flash frozen in liquid nitrogen, and stored at −80 °C. After rapid thawing, samples were resuspended in 500 µL of ice-cold cell lysis buffer (10 mM Tris-HCl pH 8.0, 10 mM NaCl, 0.5% Igepal CA-630, supplemented with the protease inhibitor cocktail from Sigma-Aldrich), incubated for 20 min on ice, and dounce-homogenized 60-80 times with a tight pestle. Extracted nuclei were centrifuged at 2,500 g for 1 min at RT (standard spinning protocol), resuspended in 300 µL 0.5x DNase I digestion buffer with 0.2% SDS and 20 mM MnCl2, and incubated at 37 °C for 60 min with periodic gentle vortexing. Then, 300 µL 0.5x DNase I digestion buffer with 2% Triton X-100, 20 mM MnCl2, and 0.4 µg/µL RNase A were added and incubated for another 10 min. Chromatin was finally digested with 7 U of DNase I during a 7 min incubation at RT. The reaction was stopped with 30 µL of 0.5M EDTA and 15 µL 10% SDS. The efficiency of DNase I digestion was confirmed by comparing the DNA shearing patterns of small samples of undigested and digested nuclei using a 6% PAGE gel (following proteinase K digestion). Nuclei were spun down and resuspended in 150 µl RNase and DNase-free water. 300 µl of AMPure XP beads (Beckman) were added to irreversibly bind nuclei; going forward nuclei were therefore cleaned up by magnetic purification. *In situ* reactions were performed first for end repair (15 U of T4 DNA Polymerase and 30 U of Klenow Fragment for 1 hr at RT) and dA-tailing (75 U of Klenow exo-minus for 1 hr at 37 °C), each in 200 µL, and followed by inactivation with 5 µL of 10% SDS and cleanup. Nuclei were then subjected to overnight ligation of custom T-and blunt-bridge biotin-tagged adapters (each at 8 µM; sequences detailed in Ramani et al., 2016) using 25 U of T4 DNA ligase and 5% polyethylene glycol (PEG) in a 100 µl reaction incubated at 16 °C. The reaction was stopped with 5 µL of 10% SDS, and nuclei were washed twice with AMPure buffer (20% PEG in 2.5 M NaCl) followed by 2 washes with 80% ethanol to remove un-ligated adapter. Nuclei were then treated for 1 hr at 37 °C with 100 U of T4 PNK in a 100 µL reaction to phosphorylate the adapters. Proximity ligation of DNA ends was performed for 4 hours at RT using 30 U of T4 DNA ligase in a 1 mL reaction maintained in gentle agitation. Nuclei were resuspended in 1x NEBuffer 2 (New England Biolabs) with 1% SDS, and digested with 800 µg of Proteinase K overnight at 62 °C. DNA was precipitated by adding 60 µg glycogen, 50 µL 3M Na-acetate (ph5.2) and 500 µL isopropanol followed by incubation for 2 hours at −80 °C and centrifugation at 16,000 g for 30 min at 4 °C. DNA was resuspended in 100 µL water and purified with 100 µL AMPure beads according to the manufacturer’s instructions, and resuspended in 100 µL elution buffer (EB; 10 mM Tris-HCl pH 8.5). Biotin pull-down was performed on the purified DNA to isolate ligation products containing the biotin adapters. 100 µL Myone C1 beads were mixed with the DNA for 30 min at RT under gentle rotation. Samples were washed 4 times with B&W buffer (5 mM Tris-HCl pH 8.0, 0.5 mM EDTA, 1 M NaCl, 0.05% Tween-20), and twice with EB. DNA was then treated on the beads to perform end-repair (200 µL reaction with the Fast DNA End Repair Kit for 10 min at 18 °C) and dA-Tailing (50 µL reaction with 25 U of Klenow exo-minus in NEBuffer 2 and 1.2 mM dATP for 30 min at 37 °C). Custom Y-adapters for Illumina sequencing (each at 2 µM; sequences detailed in Ramani et al., 2016) were then ligated using 20 U of T4 DNA ligase for 1 hour at RT in a 50 µL reaction using the Rapid Ligation Buffer. Beads were washed 4 times with B&W buffer and twice with EB after each of these reactions. Finally, libraries were amplified by 12 PCR cycles with Kapa HiFi ReadyStart Master Mix with custom barcode-containing primers (sequences detailed in Ramani et al., 2016). Libraries were purified with 0.8x Ampure XP beads according to the manufacturer’s instructions, quantified with a Qubit and the DNA high-sensitivity reagent, and pooled at equimolar ratios in preparation for next-generation sequencing. Samples were paired-end sequenced on three runs with an Illumina NextSeq 500 at high output with 150 cycles (75 for each end), resulting in approximately 150 M paired-end reads per sample.

### Hi-C data analysis

Fastq files were mapped to the hg38 genome using BWA-MEM with default parameters, mapping each end of the read pairs individually. The mapped files were processed through HiC-Pro (Servant et al., 2015), filtering for MAPQ score greater than 30 and excluding pairs less than 1 Kb apart, to generate valid pairs and ICE balanced matrices at 500 Kb resolution. Hi-C QC metrics from HiC-Pro are reported in Supplemental Table 3. Samples were clustered based on HiC-Rep scores calculated using a resolution of 500 Kb with a max distance of 5 Mb and h = 1 (Yang et al., 2017). Hi-C contact matrix heatmaps for *cis* or *trans* interactions were generated with Cooler (https://github.com/mirnylab/cooler) using default parameters and logarithmic interaction probabilities, without diagonals. A/B compartmentalization was computed by eigenvalue decomposition of the contact maps using HOMER (Heinz et al., 2010) with 500 Kb resolution and no additional windowing (super-resolution also set at 500 Kb). The sign of the first eigenvector (PC1) was selected based on the expression ~5000 genes constitutively expressed across hESC-CM differentiation (Fields et al., 2017), so that positive and negative values indicate A and B compartmentalization, respectively. Saddle plots of inter-compartment interaction enrichment were generated by assigning each genomic bin to its corresponding percentile value based on PC1, and dividing the genome into 10 deciles. Each interaction was normalized to the average score at the corresponding distance (for *cis* interactions) or to the average of all contacts (for *trans* interactions), and assigned to a pair of deciles based on the PC1 scores of the two bins. The data was plotted in heatmaps representing the log_2_ average value for pairs of deciles, while the change between mutant and corrected hiPSC-CMs is the log_2_ value of the difference in such values. To calculate the interaction probability at varying genomic distances based on compartmentalization, each interaction was assigned to A-A, B-B, or A-B based on the pairs of bins involved, and then the average interaction score for a given distance was normalized to the average interaction score for all pairs of contacts at that distance. The data was plotted on a logarithmic scale and loess-smoothed using the R function “geom_smooth”. Visualization of sample similarity by PC1 scores was performed with the R function “prcomp”. Gene tracks were generated using IGV (Thorvaldsdóttir et al., 2013). Changes in A/B compartmentalization were determined by a one-way ANOVA of PC1 scores across the two replicates for the three cell lines, using a significance cutoff of p-value < 0.05 combined by the need for the average PC1 to change sign across at least one pair of condition. Consistent changes in A/B compartmentalization were further selected if the average PC1 score for mutant hiPSC-CMs changed sign compared the average PC1 score of each corrected hiPSC-CMs.

### Statistical analyses

Unless specifically described elsewhere in the methods, all statistical analyses were performed using Prism 7 (GraphPad). The type and number of replicates, the statistics plotted, the statistical test used, and the test results are described in the figure legends. All statistical tests employed were two-tailed. No experimental samples were excluded from the statistical analyses. Sample size was not pre-determined through power calculations, and no randomization or investigator blinding approaches were implemented during the experiments and data analyses. When a representative experiment is reported, this exemplifies the results obtained in at least two independent biological replications.

### Data and code availability

Hi-C and RNA-seq data is available on Gene Expression Omnibus accession number GSE126460. Custom code for Hi-C analyses was previously described (Fields et al., 2017) and is available on github (https://github.com/pfields8/Fields_et_al_2018/). All other raw data and custom code is available from the corresponding author upon reasonable request.

## Supplemental material

The Supplemental material linked to this article includes Supplemental Figures 1 to 4 and their matching Figure Legends (presented in the same document), Supplemental Tables 1 to 5 (presented as individual files), Supplemental Tables 6 and 7 (presented in the same document as the Supplemental Figures), and Supplemental Videos 1 to 8 (presented as individual files).

## Acknowledgments

We thank the members of the Murry lab and of the UW-CNOF, especially Choli Lee for assistance with NGS, Galip Gurkan Yardimci for advice on Hi-C analysis, and Katie Mitzelfelt, Kai-Chun Yang, Elaheh Karbassi, Xiulan Yang, and Hans Reinecke for helpful discussions during the development of this project. We are also grateful to Jesse Macadangdang for assistance with CCQ analysis, and to the UW Cell Analysis Facility for support with flow cytometry analyses. A.B. is funded by an EMBO Long-Term Fellowship, ALTF 448-2017. P.F. is funded through Experimental Pathology of Cardiovascular Disease 1073 training grant, NIH T32 HL007312. This work is part of the NIH 4D Nucleome consortium (NIH U54 DK107979, to C.E.M, W.S.N. and J.S.), with additional support from National Science Foundation Grants CBET-1509106 and CMMI-1661730 (N.J.S.), F32 HL126332 (A.L.), R01 HL135143 (D-H.K. and C.E.M.), P01 GM081619 (C.E.M.), R01 HL128362 (C.E.M.), and the Foundation Leducq Transatlantic Network of Excellence (C.E.M.). The authors declare no competing financial interests.

## Author contributions

A.B. lead methodology, investigation, formal analysis, data curation, visualization, writing -original draft, and writing -review and editing; supporting conceptualization. P.F. supporting software, and formal analysis (Hi-C and RNA-seq); supporting writing – review and editing. A.S.T.S: supporting investigation and formal analysis (patch clamp); supporting writing – review and editing. A.L.: supporting methodology, software, and formal analysis (3D-EHT contractility). K.B: supporting software and formal analysis (Fluo-4). H-F.T: supporting resources (hiPSCs). N.J.S, D-H.K., J.S. W.S.N: supporting supervision, funding acquisition, and resources. L.P.: supporting supervision, project administration, and writing – review and editing. C.E.M. lead conceptualization, supervision, project administration, resources, and writing – review & editing. All authors have read, commented, and approved the final manuscript.

## Abbreviations

3D-EHT(s): three-dimensional engineered heart tissue(s)
A/B: active/inactive (chromatin compartment)
CCQ: contraction correlation quantification
FPD: field potential duration
DCM: dilated cardiomyopathy
Hi-C: genome-wide chromosome conformation capture
hESC(s): human embryonic stem cell(s)
hESC-CM(s): human embryonic stem cell-derived cardiomyocyte(s)
hiPSC(s): human induced pluripotent stem cell(s)
hiPSC-CM(s): human induced pluripotent stem cell-derived cardiomyocyte(s)
LAD(s): lamin-associated domains
MEA(s): multi-electrode array(s)
PC: principal component
RNA-seq: ribonucleic acid sequencing
RT-qPCR: quantitative reverse transcription polymerase chain reaction
TAD(s): topologically associating domain(s)

## References

Adams, M.E., I.M. Mintz, M.D. Reily, V. Thanabal, and B.P. Bean. 1993. Structure and properties of omega-agatoxin IVB, a new antagonist of P-type calcium channels. Mol. Pharmacol. 44:681–688.

Adriaens, C., L.A. Serebryannyy, M. Feric, A. Schibler, K.J. Meaburn, N. Kubben, P. Trzaskoma, S. Shachar, S. Vidak, E.H. Finn, V. Sood, G. Pegoraro, and T. Misteli. 2018. Blank spots on the map: some current questions on nuclear organization and genome architecture. Histochem. Cell Biol. 150:579–592. doi:10.1007/s00418-018-1726-1.

Akinrinade, O., L. Ollila, S. Vattulainen, J. Tallila, M. Gentile, P. Salmenperä, H. Koillinen, M. Kaartinen, M.S. Nieminen, S. Myllykangas, T.-P. Alastalo, J.W. Koskenvuo, and T. Heliö. 2015. Genetics and genotype–phenotype correlations in Finnish patients with dilated cardiomyopathy. Eur. Heart J. 36:2327–2337. doi:10.1093/eurheartj/ehv253.

Amendola, M., and B. van Steensel. 2015. Nuclear lamins are not required for lamina-associated domain organization in mouse embryonic stem cells. EMBO Rep. 16:610–7. doi:10.15252/embr.201439789.

Amin, A.S., H.L. Tan, and A.A.M. Wilde. 2010. Cardiac ion channels in health and disease. Hear. Rhythm. 7:117–126. doi:10.1016/j.hrthm.2009.08.005.

Arimura, T., A. Helbling-Leclerc, C. Massart, S. Varnous, F. Niel, E. Lacène, Y. Fromes, M. Toussaint, A.-M. Mura, D.I. Keller, H. Amthor, R. Isnard, M. Malissen, K. Schwartz, and G. Bonne. 2005. Mouse model carrying H222P- Lmna mutation develops muscular dystrophy and dilated cardiomyopathy similar to human striated muscle laminopathies. Hum. Mol. Genet. 14:155–169. doi:10.1093/hmg/ddi017.

Asakura, K., S. Hayashi, A. Ojima, T. Taniguchi, N. Miyamoto, C. Nakamori, C. Nagasawa, T. Kitamura, T. Osada, Y. Honda, C. Kasai, H. Ando, Y. Kanda, Y. Sekino, and K. Sawada. 2015. Improvement of acquisition and analysis methods in multi-electrode array experiments with iPS cell-derived cardiomyocytes. J. Pharmacol. Toxicol. Methods. 75:17–26. doi:10.1016/j.vascn.2015.04.002.

Awasthi, A., B. Ramachandran, S. Ahmed, E. Benito, Y. Shinoda, N. Nitzan, A. Heukamp, S. Rannio, H. Martens, J. Barth, K. Burk, Y.T. Wang, A. Fischer, and C. Dean. 2018. Synaptotagmin-3 drives AMPA receptor endocytosis, depression of synapse strength, and forgetting. Science. 363:eaav1483. doi:10.1126/science.aav1483.

Bellin, M., S. Casini, R.P. Davis, C. D’Aniello, J. Haas, D. Ward-van Oostwaard, L.G.J. Tertoolen, C.B. Jung, D.A. Elliott, A. Welling, K.-L. Laugwitz, A. Moretti, and C.L. Mummery. 2013. Isogenic human pluripotent stem cell pairs reveal the role of a KCNH2 mutation in long-QT syndrome. EMBO J. 32:3161–3175. doi:10.1038/emboj.2013.240.

Van Berlo, J.H., W.G. De Voogt, A.J. Van Der Kooi, J.P. Van Tintelen, G. Bonne, R. Ben Yaou, D. Duboc, T. Rossenbacker, H. Heidbüchel, M. De Visser, H.J.G.M. Crijns, and Y.M. Pinto. 2005. Meta-analysis of clinical characteristics of 299 carriers of LMNA gene mutations: Do lamin A/C mutations portend a high risk of sudden death? J. Mol. Med. 83:79–83. doi:10.1007/s00109-004-0589-1.

Bertrand, A.T., K. Chikhaoui, R. Ben Yaou, and G. Bonne. 2011. Clinical and genetic heterogeneity in laminopathies. Biochem. Soc. Trans. 39:1687–92. doi:10.1042/BST20110670.

Bodi, I., G. Mikala, S.E. Koch, S.A. Akhter, and A. Schwartz. 2005. The L-type calcium channel in the heart: the beat goes on. J. Clin. Invest. 115:3306–3317. doi:10.1172/JCI27167.

Bolzer, A., G. Kreth, I. Solovei, D. Koehler, K. Saracoglu, C. Fauth, S. Müller, R. Eils, C. Cremer, M.R. Speicher, and T. Cremer. 2005. Three-dimensional maps of all chromosomes in human male fibroblast nuclei and prometaphase rosettes. PLoS Biol. 3:e157. doi:10.1371/journal.pbio.0030157.

Bonne, G., K. Schwartz, M.R. Di Barletta, S. Varnous, H.-M. Bécane, E.-H. Hammouda, L. Merlini, F. Muntoni, C.R. Greenberg, F. Gary, J.-A. Urtizberea, D. Duboc, M. Fardeau, and D. Toniolo. 1999. Mutations in the gene encoding lamin A/C cause autosomal dominant Emery-Dreifuss muscular dystrophy. Nat. Genet. 21:285–288. doi:10.1038/6799.

Branco, M.R., and A. Pombo. 2006. Intermingling of chromosome territories in interphase suggests role in translocations and transcription-dependent associations. PLoS Biol. 4:e138. doi:10.1371/journal.pbio.0040138.

Buchwalter, A., J.M. Kaneshiro, and M.W. Hetzer. 2018. Coaching from the sidelines: the nuclear periphery in genome regulation. Nat. Rev. Genet. 20:39–50. doi:10.1038/s41576-018-0063-5.

Burridge, P.W., E. Matsa, P. Shukla, Z.C. Lin, J.M. Churko, A.D. Ebert, F. Lan, S. Diecke, B. Huber, N.M. Mordwinkin, J.R. Plews, O.J. Abilez, B. Cui, J.D. Gold, and J.C. Wu. 2014. Chemically defined generation of human cardiomyocytes. Nat. Methods. 11:855–60. doi:10.1038/nmeth.2999.

Capell, B.C., and F.S. Collins. 2006. Human laminopathies: nuclei gone genetically awry. Nat. Rev. Genet. 7:940–952. doi:10.1038/nrg1906.

Captur, G., E. Arbustini, G. Bonne, P. Syrris, K. Mills, K. Wahbi, S.A. Mohiddin, W.J. McKenna, S. Pettit, C.Y. Ho, A. Muchir, P. Gissen, P.M. Elliott, and J.C. Moon. 2018. Lamin and the heart. Heart. 104:468–479. doi:10.1136/heartjnl-2017-312338.

Carson, D., M. Hnilova, X. Yang, C.L. Nemeth, J.H. Tsui, A.S.T. Smith, A. Jiao, M. Regnier, C.E. Murry, C. Tamerler, and D.-H. Kim. 2016. Nanotopography-Induced Structural Anisotropy and Sarcomere Development in Human Cardiomyocytes Derived from Induced Pluripotent Stem Cells. ACS Appl. Mater. Interfaces. 8:21923–32. doi:10.1021/acsami.5b11671.

Catterall, W. a, E. Perez-Reyes, T.P. Snutch, and J. Striessnig. 2005. International Union of Pharmacology. XLVIII. Nomenclature and structure-function relationships of voltage-gated calcium channels. Pharmacol. Rev. 57:411–425. doi:10.1124/pr.57.4.5.units.

Cattin, M.-E., A. Muchir, and G. Bonne. 2013. “State-of-the-heart” of cardiac laminopathies. Curr Opin Cardiol. 28:297–304. doi:10.1097/HCO.0b013e32835f0c79.

Chen, E.Y., C.M. Tan, Y. Kou, Q. Duan, Z. Wang, G.V. Meirelles, N.R. Clark, and A. Ma’ayan. 2013. Enrichr: interactive and collaborative HTML5 gene list enrichment analysis tool. BMC Bioinformatics. 14:128. doi:10.1186/1471-2105-14-128.

Chen, W. V., and T. Maniatis. 2013. Clustered protocadherins. Development. 140:3297–3302. doi:10.1242/dev.090621.

Choi, J.C., A. Muchir, W. Wu, S. Iwata, S. Homma, J.P. Morrow, and H.J. Worman. 2012. Temsirolimus activates autophagy and ameliorates cardiomyopathy caused by lamin A/C gene mutation. Sci. Transl. Med. 4:144ra102. doi:10.1126/scitranslmed.3003875.

Constantinescu, D., H.L. Gray, P.J. Sammak, G.P. Schatten, and A.B. Csoka. 2006. Lamin A/C expression is a marker of mouse and human embryonic stem cell differentiation. Stem Cells. 24:177–185. doi:10.1634/stemcells.2004-0159.

Dekker, J., A.S. Belmont, M. Guttman, V.O. Leshyk, J.T. Lis, S. Lomvardas, L.A. Mirny, C.C. O’Shea, P.J. Park, B. Ren, J.C. Ritland Politz, J. Shendure, and S. Zhong. 2017. The 4D nucleome project. Nature. 549:219–226. doi:10.1038/nature23884.

Dixon, J.R., I. Jung, S. Selvaraj, Y. Shen, J.E. Antosiewicz-Bourget, A.Y. Lee, Z. Ye, A. Kim, N. Rajagopal, W. Xie, Y. Diao, J. Liang, H. Zhao, V. V. Lobanenkov, J.R. Ecker, J.A. Thomson, and B. Ren. 2015. Chromatin architecture reorganization during stem cell differentiation. Nature. 518:331–336. doi:10.1038/nature14222.

Dixon, J.R., S. Selvaraj, F. Yue, A. Kim, Y. Li, Y. Shen, M. Hu, J.S. Liu, and B. Ren. 2012. Topological domains in mammalian genomes identified by analysis of chromatin interactions. Nature. 485:376–380. doi:10.1038/nature11082.

Dobin, A., C.A. Davis, F. Schlesinger, J. Drenkow, C. Zaleski, S. Jha, P. Batut, M. Chaisson, and T.R. Gingeras. 2013. STAR: ultrafast universal RNA-seq aligner. Bioinformatics. 29:15–21. doi:10.1093/bioinformatics/bts635.

Dobrzynska, A., S. Gonzalo, C. Shanahan, and P. Askjaer. 2016. The nuclear lamina in health and disease. Nucleus. 7:233–248. doi:10.1080/19491034.2016.1183848.

Eisner, D.A., J.L. Caldwell, K. Kistamás, and A.W. Trafford. 2017. Calcium and Excitation-Contraction Coupling in the Heart. Circ. Res. 121:181–195. doi:10.1161/CIRCRESAHA.117.310230.

Fields, P.A., V. Ramani, G. Bonora, G.G. Yardimci, A. Bertero, H. Reinecke, L. Pabon, W.S. Noble, J. Shendure, and C. Murry. 2017. Dynamic reorganization of nuclear architecture during human cardiogenesis. bioRxiv. 222877. doi:10.1101/222877.

Fontana, M., A. Barison, N. Botto, L. Panchetti, G. Ricci, M. Milanesi, R. Poletti, V. Positano, G. Siciliano, C. Passino, M. Lombardi, M. Emdin, and P.G. Masci. 2013. CMR-Verified Interstitial Myocardial Fibrosis as a Marker of Subclinical Cardiac Involvement in LMNA Mutation Carriers. JACC Cardiovasc. Imaging. 6:124–126. doi:10.1016/j.jcmg.2012.06.013.

Fraser, J., C. Ferrai, A.M. Chiariello, M. Schueler, T. Rito, G. Laudanno, M. Barbieri, B.L. Moore, D.C. Kraemer, S. Aitken, S.Q. Xie, K.J. Morris, M. Itoh, H. Kawaji, I. Jaeger, Y. Hayashizaki, P. Carninci, A.R. Forrest, C.A. Semple, J. Dostie, A. Pombo, and M. Nicodemi. 2015. Hierarchical folding and reorganization of chromosomes are linked to transcriptional changes in cellular differentiation. Mol. Syst. Biol. 11:852–852. doi:10.15252/msb.20156492.

Guelen, L., L. Pagie, E. Brasset, W. Meuleman, M.B. Faza, W. Talhout, B.H. Eussen, A. de Klein, L. Wessels, W. de Laat, and B. van Steensel. 2008. Domain organization of human chromosomes revealed by mapping of nuclear lamina interactions. Nature. 453:948–51. doi:10.1038/nature06947.

Haas, J., K.S. Frese, B. Peil, W. Kloos, A. Keller, R. Nietsch, Z. Feng, S. Müller, E. Kayvanpour, B. Vogel, F. Sedaghat-Hamedani, W.K. Lim, X. Zhao, D. Fradkin, D. Köhler, S. Fischer, J. Franke, S. Marquart, I. Barb, D.T. Li, A. Amr, P. Ehlermann, D. Mereles, T. Weis, S. Hassel, A. Kremer, V. King, E. Wirsz, R. Isnard, M. Komajda, A. Serio, M. Grasso, P. Syrris, E. Wicks, V. Plagnol, L. Lopes, T. Gadgaard, H. Eiskjær, M. Jørgensen, D. Garcia-Giustiniani, M. Ortiz-Genga, M.G. Crespo-Leiro, R.H.L.D. Deprez, I. Christiaans, I.A. Van Rijsingen, A.A. Wilde, A. Waldenstrom, M. Bolognesi, R. Bellazzi, S. Mörner, J.L. Bermejo, L. Monserrat, E. Villard, J. Mogensen, Y.M. Pinto, P. Charron, P. Elliott, E. Arbustini, H.A. Katus, and B. Meder. 2015. Atlas of the clinical genetics of human dilated cardiomyopathy. Eur. Heart J. 36:1123–1135. doi:10.1093/eurheartj/ehu301.

Harr, J.C., T.R. Luperchio, X. Wong, E. Cohen, S.J. Wheelan, and K.L. Reddy. 2015. Directed targeting of chromatin to the nuclear lamina is mediated by chromatin state and A-type lamins. J. Cell Biol. 208:33–52. doi:10.1083/jcb.201405110.

Hasselberg, N.E., T.F. Haland, J. Saberniak, P.H. Brekke, K.E. Berge, T.P. Leren, T. Edvardsen, and K.H. Haugaa. 2018. Lamin A/C cardiomyopathy: Young onset, high penetrance, and frequent need for heart transplantation. Eur. Heart J. 39:853–860. doi:10.1093/eurheartj/ehx596.

Hayashi, S., and M. Takeichi. 2015. Emerging roles of protocadherins: from self-avoidance to enhancement of motility. J. Cell Sci. 128:1455–1464. doi:10.1242/jcs.166306.

Heinz, S., C. Benner, N. Spann, E. Bertolino, Y.C. Lin, P. Laslo, J.X. Cheng, C. Murre, H. Singh, and C.K. Glass. 2010. Simple combinations of lineage-determining transcription factors prime cis-regulatory elements required for macrophage and B cell identities. Mol. Cell. 38:576–89. doi:10.1016/j.molcel.2010.05.004.

Ho, J.C.Y., T. Zhou, W.H. Lai, Y. Huang, Y.C. Chan, X. Li, N.L.Y. Wong, Y. Li, K.W. Au, D. Guo, J. Xu, C.W. Siu, D. Pei, H.F. Tse, and M.A. Esteban. 2011. Generation of induced pluripotent stem cell lines from 3 distinct laminopathies bearing heterogeneous mutations in lamin A/C. Aging. 3:380–390. doi:10.18632/aging.100277.

Imakaev, M., G. Fudenberg, R.P. McCord, N. Naumova, A. Goloborodko, B.R. Lajoie, J. Dekker, and L.A. Mirny. 2012. Iterative correction of Hi-C data reveals hallmarks of chromosome organization. Nat. Methods. 9:999–1003. doi:10.1038/nmeth.2148.

Jakobs, P.M., E.L. Hanson, K.A. Crispell, W. Toy, H. Keegan, K. Schilling, T.B. Icenogle, M. Litt, and R.E. Hershberger. 2001. Novel lamin A/C mutations in two families with dilated cardiomyopathy and conduction system disease. J. Card. Fail. 7:249–256. doi:10.1054/jcaf.2001.26339.

Kalhor, R., H. Tjong, N. Jayathilaka, F. Alber, and L. Chen. 2012. Genome architectures revealed by tethered chromosome conformation capture and population-based modeling. Nat. Biotechnol. 30:90–98. doi:10.1038/nbt.2057.

Katainen, R., K. Dave, E. Pitkänen, K. Palin, T. Kivioja, N. Välimäki, A.E. Gylfe, H. Ristolainen, U.A. Hänninen, T. Cajuso, J. Kondelin, T. Tanskanen, J.-P. Mecklin, H. Järvinen, L. Renkonen-Sinisalo, A. Lepistö, E. Kaasinen, O. Kilpivaara, S. Tuupanen, M. Enge, J. Taipale, and L.A. Aaltonen. 2015. CTCF/cohesin-binding sites are frequently mutated in cancer. Nat. Genet. 47:818–21. doi:10.1038/ng.3335.

Kind, J., L. Pagie, S.S. De Vries, L. Nahidiazar, S.S. Dey, M. Bienko, Y. Zhan, B. Lajoie, C.A. De Graaf, M. Amendola, G. Fudenberg, M. Imakaev, L.A. Mirny, K. Jalink, J. Dekker, A. Van Oudenaarden, and B. Van Steensel. 2015. Genome-wide Maps of Nuclear Lamina Interactions in Single Human Cells. Cell. 163:134–147. doi:10.1016/j.cell.2015.08.040.

Kodo, K., S.-G. Ong, F. Jahanbani, V. Termglinchan, K. Hirono, K. InanlooRahatloo, A.D. Ebert, P. Shukla, O.J. Abilez, J.M. Churko, I. Karakikes, G. Jung, F. Ichida, S.M. Wu, M.P. Snyder, D. Bernstein, and J.C. Wu. 2016. iPSC-derived cardiomyocytes reveal abnormal TGF-β signalling in left ventricular non-compaction cardiomyopathy. Nat. Cell Biol. 18:1031–42. doi:10.1038/ncb3411.

Krumm, A., and Z. Duan. 2018. Understanding the 3D genome: Emerging impacts on human disease. Semin. Cell Dev. Biol. doi:10.1016/j.semcdb.2018.07.004.

Kumar, S., S.H. Baldinger, E. Gandjbakhch, P. Maury, J.M. Sellal, A.F.A. Androulakis, X. Waintraub, P. Charron, A. Rollin, P. Richard, W.G. Stevenson, C.J. Macintyre, C.Y. Ho, T. Thompson, J.K. Vohra, J.M. Kalman, K. Zeppenfeld, F. Sacher, U.B. Tedrow, and N.K. Lakdawala. 2016. Long-Term Arrhythmic and Nonarrhythmic Outcomes of Lamin A/C Mutation Carriers. J. Am. Coll. Cardiol. 68:2299–2307. doi:10.1016/j.jacc.2016.08.058.

Lee, P.J.H., D. Rudenko, M.A. Kuliszewski, C. Liao, M.G. Kabir, K.A. Connelly, and H. Leong-Poi. 2014. Survivin gene therapy attenuates left ventricular systolic dysfunction in doxorubicin cardiomyopathy by reducing apoptosis and fibrosis. Cardiovasc. Res. 101:423–433. doi:10.1093/cvr/cvu001.

Lee, Y., Y. Lau, Z. Cai, W. Lai, L. Wong, H. Tse, K. Ng, and C. Siu. 2017. Modeling Treatment Response for Lamin A/C Related Dilated Cardiomyopathy in Human Induced Pluripotent Stem Cells. J. Am. Heart Assoc. 6:e005677. doi:10.1161/JAHA.117.005677.

Leonard, A., A. Bertero, J.D. Powers, K.M. Beussman, S. Bhandari, M. Regnier, C.E. Murry, and N.J. Sniadecki. 2018. Afterload promotes maturation of human induced pluripotent stem cell derived cardiomyocytes in engineered heart tissues. J. Mol. Cell. Cardiol. 118:147–158. doi:10.1016/j.yjmcc.2018.03.016.

Lieberman-Aiden, E., N.L. van Berkum, L. Williams, M. Imakaev, T. Ragoczy, A. Telling, I. Amit, B.R. Lajoie, P.J. Sabo, M.O. Dorschner, R. Sandstrom, B. Bernstein, M.A. Bender, M. Groudine, A. Gnirke, J. Stamatoyannopoulos, L.A. Mirny, E.S. Lander, and J. Dekker. 2009. Comprehensive mapping of long-range interactions reveals folding principles of the human genome. Science. 326:289–93. doi:10.1126/science.1181369.

Liu, G.-H., B.Z. Barkho, S. Ruiz, D. Diep, J. Qu, S.-L. Yang, A.D. Panopoulos, K. Suzuki, L. Kurian, C. Walsh, J. Thompson, S. Boue, H.L. Fung, I. Sancho-Martinez, K. Zhang, J.Y. III, and J.C.I. Belmonte. 2011. Recapitulation of premature ageing with iPSCs from Hutchinson–Gilford progeria syndrome. Nature. 472:221–225. doi:10.1038/nature09879.

Luperchio, T.R., M.E.G. Sauria, X. Wong, M. Gaillard, P. Tsang, and K. Pekrun. 2017. Chromosome Conformation Paints Reveal the Role of Lamina Association in Genome Organization and Regulation. bioRxiv. doi:10.1101/122226.

Lupiáñez, D.G., K. Kraft, V. Heinrich, P. Krawitz, F. Brancati, E. Klopocki, D. Horn, H. Kayserili, J.M. Opitz, R. Laxova, F. Santos-Simarro, B. Gilbert-Dussardier, L. Wittler, M. Borschiwer, S.A. Haas, M. Osterwalder, M. Franke, B. Timmermann, J. Hecht, M. Spielmann, A. Visel, and S. Mundlos. 2015. Disruptions of topological chromatin domains cause pathogenic rewiring of gene-enhancer interactions. Cell. 161:1012–1025. doi:10.1016/j.cell.2015.04.004.

Macadangdang, J., X. Guan, A.S.T. Smith, R. Lucero, S. Czerniecki, M.K. Childers, D.L. Mack, and D.-H. Kim. 2015. Nanopatterned Human iPSC-based Model of a Dystrophin-Null Cardiomyopathic Phenotype. Cell. Mol. Bioeng. 8:320–332. doi:10.1007/s12195-015-0413-8.

Maruo, T., K. Mandai, M. Miyata, S. Sakakibara, S. Wang, K. Sai, Y. Itoh, A. Kaito, T. Fujiwara, A. Mizoguchi, and Y. Takai. 2017. NGL-3-induced presynaptic differentiation of hippocampal neurons in an afadin-dependent, nectin-1-independent manner. Genes Cells. 22:742–755. doi:10.1111/gtc.12510.

McDonough, S.I., K.J. Swartz, I.M. Mintz, L.M. Boland, and B.P. Bean. 1996. Inhibition of calcium channels in rat central and peripheral neurons by omega-conotoxin MVIIC. J Neurosci. 16:2612–2623. doi:10.1523/JNEUROSCI.16-08-02612.1996.

Meaburn, K.J., E. Cabuy, G. Bonne, N. Levy, G.E. Morris, G. Novelli, I.R. Kill, and J.M. Bridger. 2007. Primary laminopathy fibroblasts display altered genome organization and apoptosis. Aging Cell. 6:139–153. doi:10.1111/j.1474-9726.2007.00270.x.

Merideth, M.A., L.B. Gordon, S. Clauss, V. Sachdev, A.C.M. Smith, M.B. Perry, C.C. Brewer, C. Zalewski, H.J. Kim, B. Solomon, B.P. Brooks, L.H. Gerber, M.L. Turner, D.L. Domingo, T.C. Hart, J. Graf, J.C. Reynolds, A. Gropman, J.A. Yanovski, M. Gerhard-Herman, F.S. Collins, E.G. Nabel, R.O. Cannon, W.A. Gahl, and W.J. Introne. 2008. Phenotype and Course of Hutchinson–Gilford Progeria Syndrome. N. Engl. J. Med. 358:592–604. doi:10.1145/2503210.2503268.

Metra, M., and J.R. Teerlink. 2017. Heart failure. Lancet. 390:1981–1995. doi:10.1016/S0140-6736(17)31071-1.

Mewborn, S.K., M.J. Puckelwartz, F. Abuisneineh, J.P. Fahrenbach, Y. Zhang, H. MacLeod, L. Dellefave, P. Pytel, S. Selig, C.M. Labno, K. Reddy, H. Singh, and E. McNally. 2010. Altered chromosomal positioning, Compaction, and gene expression with a lamin A/C gene mutation. PLoS One. 5:e14342. doi:10.1371/journal.pone.0014342.

Molkentin, J.D., J.R. Lu, C.L. Antos, B. Markham, J. Richardson, J. Robbins, S.R. Grant, and E.N. Olson. 1998. A calcineurin-dependent transcriptional pathway for cardiac hypertrophy. Cell. 93:215–28. doi:10.1016/S0092-8674(00)81573-1.

Mosqueira, D., I. Mannhardt, J.R. Bhagwan, K. Lis-Slimak, P. Katili, E. Scott, M. Hassan, M. Prondzynski, S.C. Harmer, A. Tinker, J.G.W. Smith, L. Carrier, P.M. Williams, D. Gaffney, T. Eschenhagen, A. Hansen, and C. Denning. 2018. CRISPR/Cas9 editing in human pluripotent stem cell-cardiomyocytes highlights arrhythmias, hypocontractility, and energy depletion as potential therapeutic targets for hypertrophic cardiomyopathy. Eur. Heart J. 39:3879–3892. doi:10.1093/eurheartj/ehy249.

Muchir, A., P. Pavlidis, V. Decostre, A.J. Herron, T. Arimura, G. Bonne, and H.J. Worman. 2007. Activation of MAPK pathways links LMNA mutations to cardiomyopathy in Emery-Dreifuss muscular dystrophy. J. Clin. Invest. 117:1282–93. doi:10.1172/JCI29042.

Muchir, A., W. Wu, J.C. Choi, S. Iwata, J. Morrow, S. Homma, and H.J. Worman. 2012. Abnormal p38 mitogen-activated protein kinase signaling in dilated cardiomyopathy caused by lamin A/C gene mutation. Hum. Mol. Genet. 21:4325–4333. doi:10.1093/hmg/dds265.

Nakamura, T.Y., Y. Iwata, Y. Arai, K. Komamura, and S. Wakabayashi. 2008. Activation of Na+/H+ Exchanger 1 Is Sufficient to Generate Ca2+ Signals That Induce Cardiac Hypertrophy and Heart Failure. Circ. Res. 103:891–899. doi:10.1161/CIRCRESAHA.108.175141.

Nie, J., Q. Duan, M. He, X. Li, B. Wang, C. Zhou, L. Wu, Z. Wen, C. Chen, D.W. Wang, K.M. Alsina, X.H.T. Wehrens, D.W. Wang, and L. Ni. 2018. Ranolazine prevents pressure overload-induced cardiac hypertrophy and heart failure by restoring aberrant Na+ and Ca2+ handling. J. Cell. Physiol. doi:10.1002/jcp.27791.

Nimmrich, V., and G. Gross. 2012. P/Q-type calcium channel modulators. Br. J. Pharmacol. 167:741–759. doi:10.1111/j.1476-5381.2012.02069.x.

Nora, E.P., B.R. Lajoie, E.G. Schulz, L. Giorgetti, I. Okamoto, N. Servant, T. Piolot, N.L. van Berkum, J. Meisig, J. Sedat, J. Gribnau, E. Barillot, N. Blüthgen, J. Dekker, and E. Heard. 2012. Spatial partitioning of the regulatory landscape of the X-inactivation centre. Nature. 485:381–385. doi:10.1038/nature11049.

Ortmann, D., and L. Vallier. 2017. Variability of human pluripotent stem cell lines. Curr. Opin. Genet. Dev. 46:179–185. doi:10.1016/j.gde.2017.07.004.

Pan, H., A.A. Richards, X. Zhu, J.A. Joglar, H.L. Yin, and V. Garg. 2009. A novel mutation in LAMIN A/C is associated with isolated early-onset atrial fibrillation and progressive atrioventricular block followed by cardiomyopathy and sudden cardiac death. Hear. Rhythm. 6:707–710. doi:10.1016/j.hrthm.2009.01.037.

Peroz, D., N. Rodriguez, F. Choveau, I. Baró, J. Mérot, and G. Loussouarn. 2008. Kv7.1 (KCNQ1) properties and channelopathies. J. Physiol. 586:1785–1789. doi:10.1113/jphysiol.2007.148254.

Rajakulendran, S., D. Kaski, and M.G. Hanna. 2012. Neuronal P/Q-type calcium channel dysfunction in inherited disorders of the CNS. Nat. Rev. Neurol. 8:86–96. doi:10.1038/nrneurol.2011.228.

Ramani, V., D.A. Cusanovich, R.J. Hause, W. Ma, R. Qiu, X. Deng, C.A. Blau, C.M. Disteche, W.S. Noble, J. Shendure, and Z. Duan. 2016. Mapping 3D genome architecture through in situ DNase Hi-C. Nat Protoc. 11:2104–2121. doi:10.1038/nprot.2016.126.

Ramos, F.J., S.C. Chen, M.G. Garelick, D.-F. Dai, C.-Y. Liao, K.H. Schreiber, V.L. MacKay, E.H. An, R. Strong, W.C. Ladiges, P.S. Rabinovitch, M. Kaeberlein, and B.K. Kennedy. 2012. Rapamycin reverses elevated mTORC1 signaling in lamin A/C- deficient mice, rescues cardiac and skeletal muscle function, and extends survival. Sci. Transl. Med. 4:144ra103. doi:10.1126/scitranslmed.3003802.

Ran, F.A., P.D. Hsu, J. Wright, V. Agarwala, D.A. Scott, and F. Zhang. 2013. Genome engineering using the CRISPR-Cas9 system. Nat. Protoc. 8:2281–308. doi:10.1038/nprot.2013.143.

Rao, S.S.P., M.H. Huntley, N.C. Durand, E.K. Stamenova, I.D. Bochkov, J.T. Robinson, A.L. Sanborn, I. Machol, A.D. Omer, E.S. Lander, and E.L. Aiden. 2014. A 3D map of the human genome at kilobase resolution reveals principles of chromatin looping. Cell. 159:1665–1680. doi:10.1016/j.cell.2014.11.021.

Rast, G., U. Kraushaar, S. Buckenmaier, C. Ittrich, and B.D. Guth. 2016. Influence of field potential duration on spontaneous beating rate of human induced pluripotent stem cell-derived cardiomyocytes: Implications for data analysis and test system selection. J. Pharmacol. Toxicol. Methods. 82:74–82. doi:10.1016/j.vascn.2016.08.002.

Redin, C., H. Brand, R.L. Collins, T. Kammin, E. Mitchell, J.C. Hodge, C. Hanscom, V. Pillalamarri, C.M. Seabra, M.-A. Abbott, O.A. Abdul-Rahman, E. Aberg, R. Adley, S.L. Alcaraz-Estrada, F.S. Alkuraya, Y. An, M.-A. Anderson, C. Antolik, K. Anyane-Yeboa, J.F. Atkin, T. Bartell, J.A. Bernstein, E. Beyer, I. Blumenthal, E.M.H.F. Bongers, E.H. Brilstra, C.W. Brown, H.T. Brüggenwirth, B. Callewaert, C. Chiang, K. Corning, H. Cox, E. Cuppen, B.B. Currall, T. Cushing, D. David, M.A. Deardorff, A. Dheedene, M. D’Hooghe, B.B.A. de Vries, D.L. Earl, H.L. Ferguson, H. Fisher, D.R. FitzPatrick, P. Gerrol, D. Giachino, J.T. Glessner, T. Gliem, M. Grady, B.H. Graham, C. Griffis, K.W. Gripp, A.L. Gropman, A. Hanson-Kahn, D.J. Harris, M.A. Hayden, R. Hill, R. Hochstenbach, J.D. Hoffman, R.J. Hopkin, M.W. Hubshman, A.M. Innes, M. Irons, M. Irving, J.C. Jacobsen, S. Janssens, T. Jewett, J.P. Johnson, M.C. Jongmans, S.G. Kahler, D.A. Koolen, J. Korzelius, P.M. Kroisel, Y. Lacassie, W. Lawless, E. Lemyre, K. Leppig, A. V Levin, H. Li, H. Li, E.C. Liao, C. Lim, E.J. Lose, D. Lucente, M.J. Macera, P. Manavalan, G. Mandrile, C.L. Marcelis, L. Margolin, T. Mason, D. Masser-Frye, M.W. McClellan, C.J.Z. Mendoza, B. Menten, S. Middelkamp, L.R. Mikami, E. Moe, S. Mohammed, et al. 2017. The genomic landscape of balanced cytogenetic abnormalities associated with human congenital anomalies. Nat. Genet. 49:36–45. doi:10.1038/ng.3720.

Van Rijsingen, I.A.W., E. Arbustini, P.M. Elliott, J. Mogensen, J.F. Hermans-Van Ast, A.J. Van Der Kooi, J.P. Van Tintelen, M.P. Van Den Berg, A. Pilotto, M. Pasotti, S. Jenkins, C. Rowland, U. Aslam, A.A.M. Wilde, A. Perrot, S. Pankuweit, A.H. Zwinderman, P. Charron, and Y.M. Pinto. 2012. Risk factors for malignant ventricular arrhythmias in Lamin A/C mutation carriers: A European cohort study. J. Am. Coll. Cardiol. 59:493–500. doi:10.1016/j.jacc.2011.08.078.

Roncarati, R., C. Viviani Anselmi, P. Krawitz, G. Lattanzi, Y. von Kodolitsch, A. Perrot, E. di Pasquale, L. Papa, P. Portararo, M. Columbaro, A. Forni, G. Faggian, G. Condorelli, and P.N. Robinson. 2013. Doubly heterozygous LMNA and TTN mutations revealed by exome sequencing in a severe form of dilated cardiomyopathy. Eur. J. Hum. Genet. 21:1105–11. doi:10.1038/ejhg.2013.16.

Ruan, J.L., N.L. Tulloch, M. V. Razumova, M. Saiget, V. Muskheli, L. Pabon, H. Reinecke, M. Regnier, and C.E. Murry. 2016. Mechanical Stress Conditioning and Electrical Stimulation Promote Contractility and Force Maturation of Induced Pluripotent Stem Cell-Derived Human Cardiac Tissue. Circulation. 134:1557–1567. doi:10.1161/CIRCULATIONAHA.114.014998.

Russo, V., A. Rago, L. Politano, A.A. Papa, F. Di Meo, M.G. Russo, P. Golino, R. Calabrò, and G. Nigro. 2012. Increased dispersion of ventricular repolarization in emery dreifuss muscular dystrophy patients. Med Sci Monit. 18:643–647. doi:10.12659/MSM.883541.

Saga, A., A. Karibe, J. Otomo, K. Iwabuchi, T. Takahashi, H. Kanno, J. Kikuchi, M. Keitoku, T. Shinozaki, and H. Shimokawa. 2009. Lamin A/C gene mutations in familial cardiomyopathy with advanced atrioventricular block and arrhythmia. Tohoku J. Exp. Med. 218:309–16. doi:doi.org/10.1620/tjem.218.309.

Sala, L., M. Bellin, and C.L. Mummery. 2017. Integrating cardiomyocytes from human pluripotent stem cells in safety pharmacology: has the time come? Br. J. Pharmacol. 174:3749–3765. doi:10.1111/bph.13577.

Schmitt, A.D., M. Hu, I. Jung, Z. Xu, Y. Qiu, C.L. Tan, Y. Li, S. Lin, Y. Lin, C.L. Barr, and B. Ren. 2016. A Compendium of Chromatin Contact Maps Reveals Spatially Active Regions in the Human Genome. Cell Rep. 17:2042–2059. doi:10.1016/j.celrep.2016.10.061.

Servant, N., N. Varoquaux, B.R. Lajoie, E. Viara, C.-J. Chen, J.-P. Vert, E. Heard, J. Dekker, and E. Barillot. 2015. HiC-Pro: an optimized and flexible pipeline for Hi-C data processing. Genome Biol. 16:259. doi:10.1186/s13059-015-0831-x.

Sexton, T., E. Yaffe, E. Kenigsberg, F. Bantignies, B. Leblanc, M. Hoichman, H. Parrinello, A. Tanay, and G. Cavalli. 2012. Three-Dimensional Folding and Functional Organization Principles of the Drosophila Genome. Cell. 148:458–472. doi:10.1016/j.cell.2012.01.010.

Simonis, M., P. Klous, E. Splinter, Y. Moshkin, R. Willemsen, E. de Wit, B. van Steensel, and W. de Laat. 2006. Nuclear organization of active and inactive chromatin domains uncovered by chromosome conformation capture–on-chip (4C). Nat. Genet. 38:1348–1354. doi:10.1038/ng1896.

Siu, C.W., Y.K. Lee, J.C.Y. Ho, W.H. Lai, Y.C. Chan, K.M. Ng, L.Y. Wong, K.W. Au, Y.M. Lau, J. Zhang, K.W. Lay, A. Colman, and H.F. Tse. 2012. Modeling of lamin A/C mutation premature cardiac aging using patient-specific induced pluripotent stem cells. Aging. 4:803–822. doi:10.18632/aging.100503.

Slaymaker, I.M., L. Gao, B. Zetsche, D. a. Scott, W.X. Yan, and F. Zhang. 2015. Rationally engineered Cas9 nucleases with improved specificity. Science. 351:84–88. doi:10.1126/science.aad5227.

Sniadecki, N.J., and C.S. Chen. 2007. Microfabricated Silicone Elastomeric Post Arrays for Measuring Traction Forces of Adherent Cells. In Methods in Cell Biology. 313–328.

Solovei, I., A.S. Wang, K. Thanisch, C.S. Schmidt, S. Krebs, M. Zwerger, T. V. Cohen, D. Devys, R. Foisner, L. Peichl, H. Herrmann, H. Blum, D. Engelkamp, C.L. Stewart, H. Leonhardt, and B. Joffe. 2013. LBR and lamin A/C sequentially tether peripheral heterochromatin and inversely regulate differentiation. Cell. 152:584–598. doi:10.1016/j.cell.2013.01.009.

van Steensel, B., and A.S. Belmont. 2017. Lamina-Associated Domains: Links with Chromosome Architecture, Heterochromatin, and Gene Repression. Cell. 169:780–791. doi:10.1016/j.cell.2017.04.022.

Stevens, T.J., D. Lando, S. Basu, L.P. Atkinson, Y. Cao, S.F. Lee, M. Leeb, K.J. Wohlfahrt, W. Boucher, A. O’Shaughnessy-Kirwan, J. Cramard, A.J. Faure, M. Ralser, E. Blanco, L. Morey, M. Sansó, M.G.S. Palayret, B. Lehner, L. Di Croce, A. Wutz, B. Hendrich, D. Klenerman, and E.D. Laue. 2017. 3D structures of individual mammalian genomes studied by single-cell Hi-C. Nature. 544:59–64. doi:10.1038/nature21429.

Strate, I., F. Tessadori, and J. Bakkers. 2015. Glypican4 promotes cardiac specification and differentiation by attenuating canonical Wnt and Bmp signaling. Development. 142:1767–1776. doi:10.1242/dev.113894.

Sun, J.H., L. Zhou, D.J. Emerson, S.A. Phyo, K.R. Titus, W. Gong, T.G. Gilgenast, J.A. Beagan, B.L. Davidson, F. Tassone, and J.E. Phillips-Cremins. 2018. Disease-Associated Short Tandem Repeats Co-localize with Chromatin Domain Boundaries. Cell. 175:224–238.e15. doi:10.1016/j.cell.2018.08.005.

Taniura, H., C. Glass, and L. Gerace. 1995. A chromatin binding site in the tail domain of nuclear lamins that interacts with core histones. J. Cell Biol. 131:33–44. doi:10.1083/jcb.131.1.33.

Thorvaldsdóttir, H., J.T. Robinson, and J.P. Mesirov. 2013. Integrative Genomics Viewer (IGV): high-performance genomics data visualization and exploration. Brief. Bioinform. 14:178–92. doi:10.1093/bib/bbs017.

van Tintelen, J.P., R.M.W. Hofstra, H. Katerberg, T. Rossenbacker, A.C.P. Wiesfeld, G.J. du Marchie Sarvaas, A.A.M. Wilde, I.M. van Langen, E.A. Nannenberg, A.J. van der Kooi, M. Kraak, I.C. van Gelder, D.J. van Veldhuisen, Y. Vos, and M.P. van den Berg. 2007a. High yield of LMNA mutations in patients with dilated cardiomyopathy and/or conduction disease referred to cardiogenetics outpatient clinics. Am. Heart J. 154:1130–9. doi:10.1016/j.ahj.2007.07.038.

van Tintelen, J.P., R.A. Tio, W.S. Kerstjens-Frederikse, J.H. van Berlo, L.G. Boven, A.J.H. Suurmeijer, S.J. White, J.T. den Dunnen, G.J. te Meerman, Y.J. Vos, A.H. van der Hout, J. Osinga, M.P. van den Berg, D.J. van Veldhuisen, C.H.C.M. Buys, R.M.W. Hofstra, and Y.M. Pinto. 2007b. Severe myocardial fibrosis caused by a deletion of the 5’ end of the lamin A/C gene. J. Am. Coll. Cardiol. 49:2430–9. doi:10.1016/j.jacc.2007.02.063.

Tobita, T., S. Nomura, T. Fujita, H. Morita, Y. Asano, K. Onoue, M. Ito, Y. Imai, A. Suzuki, T. Ko, M. Satoh, K. Fujita, A.T. Naito, Y. Furutani, H. Toko, M. Harada, E. Amiya, M. Hatano, E. Takimoto, T. Shiga, T. Nakanishi, Y. Sakata, M. Ono, Y. Saito, S. Takashima, N. Hagiwara, H. Aburatani, and I. Komuro. 2018. Genetic basis of cardiomyopathy and the genotypes involved in prognosis and left ventricular reverse remodeling. Sci. Rep. 8:1–11. doi:10.1038/s41598-018-20114-9.

Trapnell, C., A. Roberts, L. Goff, G. Pertea, D. Kim, D.R. Kelley, H. Pimentel, S.L. Salzberg, J.L. Rinn, and L. Pachter. 2012. Differential gene and transcript expression analysis of RNA-seq experiments with TopHat and Cufflinks. Nat. Protoc. 7:562–578. doi:10.1038/nprot.2012.016.

Trapnell, C., B.A. Williams, G. Pertea, A. Mortazavi, G. Kwan, M.J. van Baren, S.L. Salzberg, B.J. Wold, and L. Pachter. 2010. Transcript assembly and quantification by RNA-Seq reveals unannotated transcripts and isoform switching during cell differentiation. Nat. Biotechnol. 28:511–515. doi:10.1038/nbt.1621.

Vandael, E., B. Vandenberk, J. Vandenberghe, R. Willems, and V. Foulon. 2017. Risk factors for QTc-prolongation: systematic review of the evidence. Int. J. Clin. Pharm. 39:16–25. doi:10.1007/s11096-016-0414-2.

Wang, Y., A. Khan, S. Heringer-Walther, H.-P. Schultheiss, M. da C. V Moreira, and T. Walther. 2013. Prognostic value of circulating levels of stem cell growth factor beta (SCGF beta) in patients with Chagas’ disease and idiopathic dilated cardiomyopathy. Cytokine. 61:728–31. doi:10.1016/j.cyto.2012.12.018.

Won, H., L. de la Torre-Ubieta, J.L. Stein, N.N. Parikshak, J. Huang, C.K. Opland, M.J. Gandal, G.J. Sutton, F. Hormozdiari, D. Lu, C. Lee, E. Eskin, I. Voineagu, J. Ernst, and D.H. Geschwind. 2016. Chromosome conformation elucidates regulatory relationships in developing human brain. Nature. 538:523–27. doi:10.1038/nature19847.

Worman, H.J., and J.-C. Courvalin. 2004. How do mutations in lamins A and C cause disease? J. Clin. Invest. 113:349–51. doi:10.1172/JCI20832.

Yang, T., F. Zhang, G.G. Yardımcı, F. Song, R.C. Hardison, W.S. Noble, F. Yue, and Q. Li. 2017. HiCRep: assessing the reproducibility of Hi-C data using a stratum-adjusted correlation coefficient. Genome Res. 27:1939–1949. doi:10.1101/gr.220640.117.

Yang, X., L. Pabon, and C.E. Murry. 2014. Engineering adolescence: Maturation of human pluripotent stem cell-derived cardiomyocytes. Circ. Res. 114:511–523. doi:10.1161/CIRCRESAHA.114.300558.

Ye, Q., and H.J. Worman. 1994. Primary structure analysis and lamin B and DNA binding of human LBR, an integral protein of the nuclear envelope inner membrane. J. Biol. Chem. 269:11306–11.

Ye, Q., and H.J. Worman. 1996. Interaction between an integral protein of the nuclear envelope inner membrane and human chromodomain proteins homologous to Drosophila HP1. J. Biol. Chem. 271:14653–6. doi:10.1074/jbc.271.25.14653.

Yu, M., and B. Ren. 2017. The Three-Dimensional Organization of Mammalian Genomes. Annu. Rev. Cell Dev. Biol. 33:265–289. doi:10.1146/annurev-cellbio-100616-060531.

Yuan, J., G. Simos, G. Blobel, and S.D. Georgatos. 1991. Binding of lamin A to polynucleosomes. J. Biol. Chem. 266:9211–5.

Yusa, K. 2013. Seamless genome editing in human pluripotent stem cells using custom endonuclease-based gene targeting and the piggyBac transposon. Nat. Protoc. 8:2061–78. doi:10.1038/nprot.2013.126.

Yusa, K., S.T. Rashid, H. Strick-Marchand, I. Varela, P.-Q. Liu, D.E. Paschon, E. Miranda, A. Ordóñez, N.R.F. Hannan, F.J. Rouhani, S. Darche, G. Alexander, S.J. Marciniak, N. Fusaki, M. Hasegawa, M.C. Holmes, J.P. Di Santo, D. a Lomas, A. Bradley, and L. Vallier. 2011. Targeted gene correction of α1-antitrypsin deficiency in induced pluripotent stem cells. Nature. 478:391–4. doi:10.1038/nature10424.

Zhang, X., X. Ai, H. Nakayama, B. Chen, D.M. Harris, M. Tang, Y. Xie, C. Szeto, Y. Li, Y. Li, H. Zhang, A.D. Eckhart, W.J. Koch, J.D. Molkentin, and X. Chen. 2016. Persistent increases in Ca2+ influx through Cav1.2 shortens action potential and causes Ca2+ overload-induced afterdepolarizations and arrhythmias. Basic Res. Cardiol. 111. doi:10.1007/s00395-015-0523-4.

Zheng, X., J. Hu, S. Yue, L. Kristiani, M. Kim, M. Sauria, J. Taylor, Y. Kim, and Y. Zheng. 2018. Lamins Organize the Global Three-Dimensional Genome from the Nuclear Periphery. Mol. Cell. 71:802–815.e7. doi:10.1016/j.molcel.2018.05.017.

Zheng, X., Y. Kim, and Y. Zheng. 2015. Identification of lamin B-regulated chromatin regions based on chromatin landscapes. Mol. Biol. Cell. 26:2685–2697. doi:10.1091/mbc.E15-04-0210.

