## Supplemental material for "Chromatin compartment dynamics in a haploinsufficient model of cardiac laminopathy"

Bertero et al.

### Supplemental material

**<u>Guide to content:</u>**

**Supplemental Figures and Figure Legends**

Supplemental Figure 1. Scarless correction of the *LMNA* R225X heterozygous mutation in hiPSCs.

Supplemental Figure 2. Characterization of lamin A/C haploinsufficient hiPSC-CMs.

Supplemental Figure 3. Global properties of chromatin topology in lamin A/C haploinsufficient hiPSC-CMs.

Supplemental Figure 4. Chromatin compartment transitions and associated gene expression changes in lamin A/C haploinsufficient hiPSC-CMs.

**Supplemental Tables**

Supplemental Table 1 (online). Differential gene expression analyses of RNA-seq data .

Supplemental Table 2 (online). Gene ontology analyses of RNA-seq data.

Supplemental Table 3 (online). QC metrics of DNase Hi-C experiments.

Supplemental Table 4 (online). A/B compartment analyses of Hi-C data.

Supplemental Table 5 (online). Ontology enrichment analyses of genes in lamin A/C-sensitive B compartments.

Supplemental Table 6. Genotyping strategies for the characterization of CRISPR/Cas9-edited hiPSCs.

Supplemental Table 7. RT-qPCR primers.

**Supplemental Videos and Supplemental Video Legends**

Supplemental Video 1 (online). Spontaneous contractions of mutant hiPSC-CMs.

Supplemental Video 2 (online). Spontaneous contractions of corrected hiPSC-CMs.

Supplemental Video 3 (online). Calcium fluxes indicated by Fluo-4 fluorescence in monolayers of mutant hiPSC-CMs electrically paced at 1 Hz.

Supplemental Video 4 (online). Calcium fluxes indicated by Fluo-4 fluorescence in monolayers of corrected hiPSC-CMs electrically paced at 1 Hz.

Supplemental Video 5 (online). Contraction correlation quantification (CCQ) analyses in monolayers of mutant hiPSC-CMs electrically paced at 1 Hz.

Supplemental Video 6 (online). Contraction correlation quantification (CCQ) analyses in monolayers of corrected hiPSC-CMs electrically paced at 1 Hz.

Supplemental Video 7. (online). Contractions of 3D-EHTs from mutant hiPSC-CMs.

Supplemental Video 8 (online). Contractions of 3D-EHTs from corrected hiPSC-CMs.

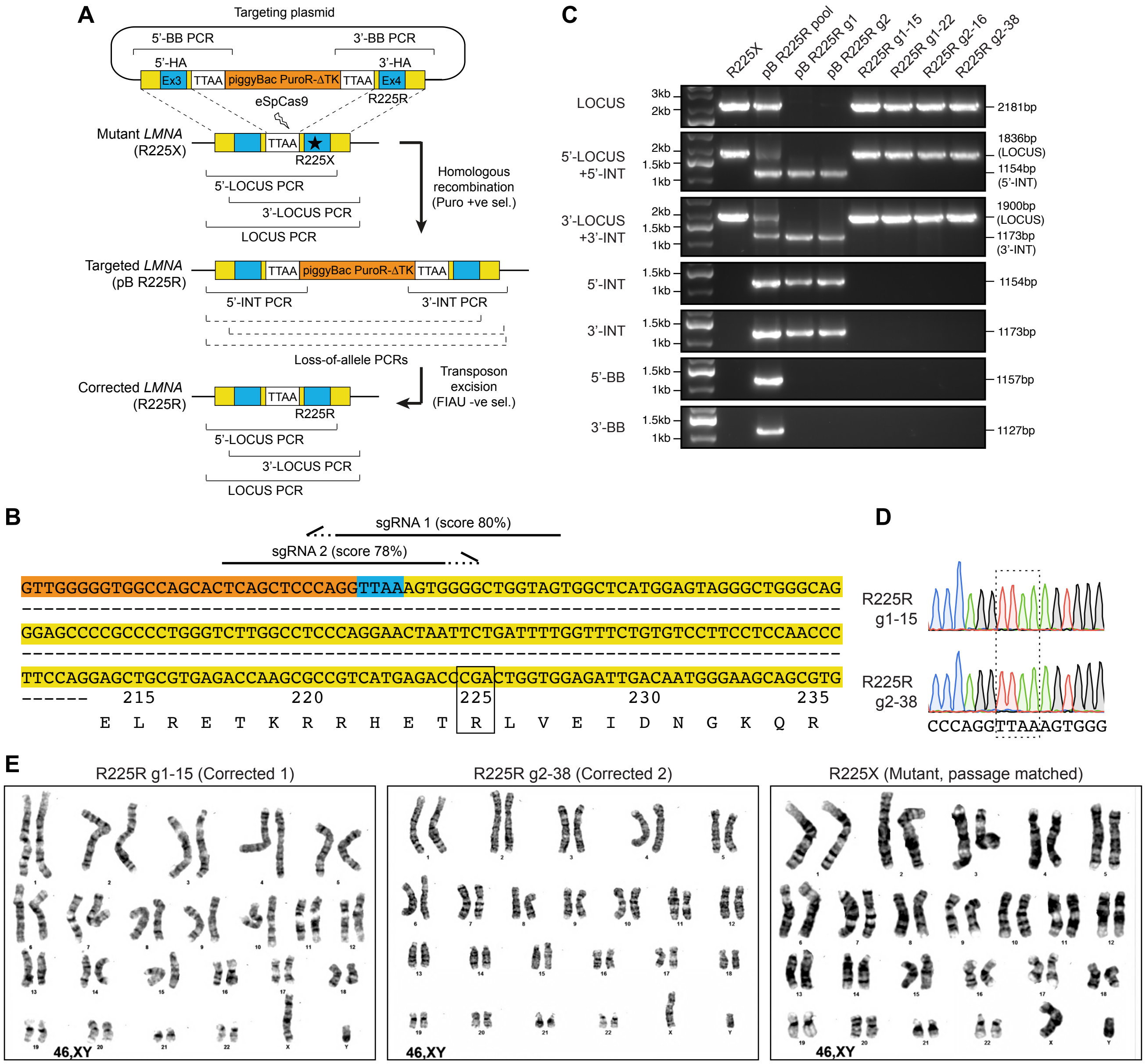


**Supplemental Figure 1. Scarless correction of the *LMNA* R225X heterozygous mutation in hiPSCs.**

(**A**) Schematics of the gene targeting and genotyping strategies. A targeting vector with wild type up- and downstream homology arms (5'- and 3'-HA; Ex: exon) and a polycistronic piggyBac cassette carrying puromycin N-acetyltransferase and Herpes Simplex truncated thymidine kinase (PuroR-ΔTK) was used in combination with enhanced specificity Cas9 (eSpCas9) to drive homologous recombination around an endogenous "TTAA" sequence proximal to the R225X mutation. Targeted clones were positively selected based on resistance to puromycin and screened by a panel of genomic PCR assays indicated by square brackets (PCR failing for gene-targeted alleles are indicated by dashed square brackets; see the Material and Methods for details). The piggyBac cassette was subsequently excised using transposase leaving behind only a "TTAA" scar identical to the original sequence, and clones were negatively selected based on sensitivity to fialuridine (FIAU). Corrected clones were screened by genomic PCR, Sanger sequencing, and karyotyping. (**B**) Genomic sequence of the *LMNA* locus around the R225X mutation (highlighted by the box). The amino acid translation of portion of *LMNA* exon 4 is also reported. The location and directionality of the sgRNAs used for CRISPR/Cas9 targeting are indicated by the half arrows (dashed portion: PAM site; specificity scores from: www.crispr.mit.edu). The 5’- and 3’-homology arms are shown with orange and yellow backgrounds, respectively. The “TTAA” sequence key to the scarless gene targeting strategy is shown with a blue background. (**C**) Selected genomic PCR results from the indicated hiPSC lines at different stages of the gene targeting procedure (see panel A). Pool: non-clonally selected hiPSCs after the first targeting step, providing a positive control for all PCR strategies. g1/g2: sgRNA 1/sgRNA 2 used for the first targeting step. (**D**) Sanger sequencing results for genomic PCR products (LOCUS PCR) from the two corrected hiPSC clones selected for further experiments. The “TTAA” sequence was faithfully reconstituted following transposase excision of the piggyBac cassette (see Fig. 1B for the R225R correction). (**E**) Representative G-banding karyotypes for the two corrected hiPSCs and passage-matched mutant hiPSCs, confirming a normal male 46 XY karyotype.

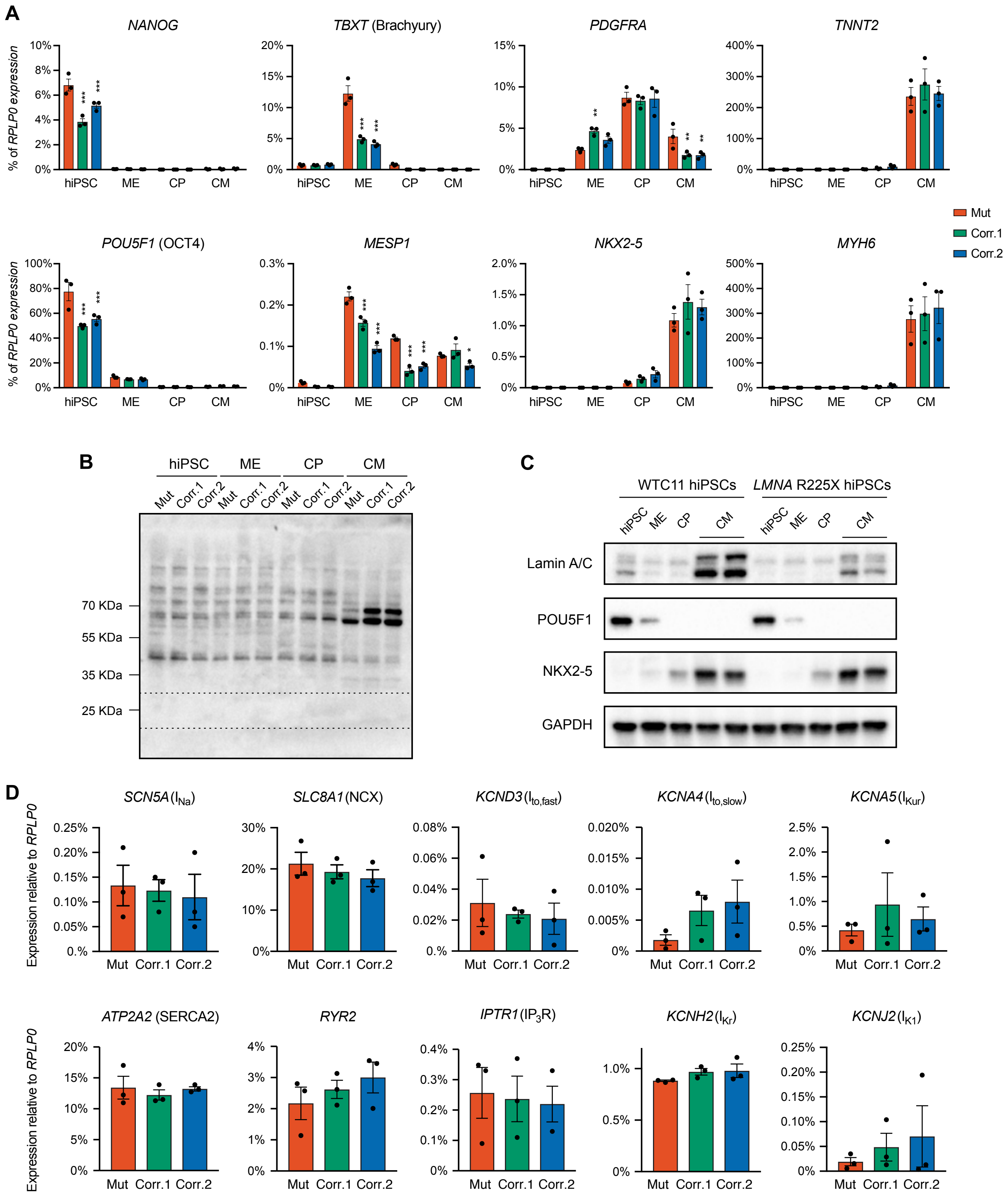


**Supplemental Figure 2. Characterization of lamin A/C haploinsufficient hiPSC-CMs.**

(**A**) RT-qPCR analyses for lineage-specific markers at the indicated stages of hiPSC-CM differentiation (ME: mesoderm; CP: cardiac progenitor; CM: cardiomyocyte; see Fig. 1C). Differences *versus* mutant were calculated by two-way ANOVA with post-hoc Holm-Sidak binary comparisons (* = $p < 0.05$, ** = $p < 0.05$, *** = $p < 0.001$; n = 3 differentiations; average ± SEM). (**B-C**) Representative Western blots for lamin A/C and differentiation markers during iPSC-CM differentiation. In B, the uncropped image shows a stronger exposure of the same lamin A/C Western blot presented in Fig. 1F. The expected location for putative truncated Lamin A and Lamin C due the R225X mutation is indicated by the dashed box (see Fig. 1A), but no such truncation product could be detected. (**D**) RT-qPCR in hiPSC-CMs at day 14 of differentiation. No significant differences were detected *versus* mutant (two-way ANOVA with post-hoc Holm-Sidak binary comparisons; n = 3 differentiations; average ± SEM). Throughout the figure (and in all other Supplemental figures), Mut or Mutant indicates *LMNA* R225X hiPSCs, and Corr.1/2 or Corrected 1/2 indicate the two isogenic corrected control *LMNA* R225R hiPSC.

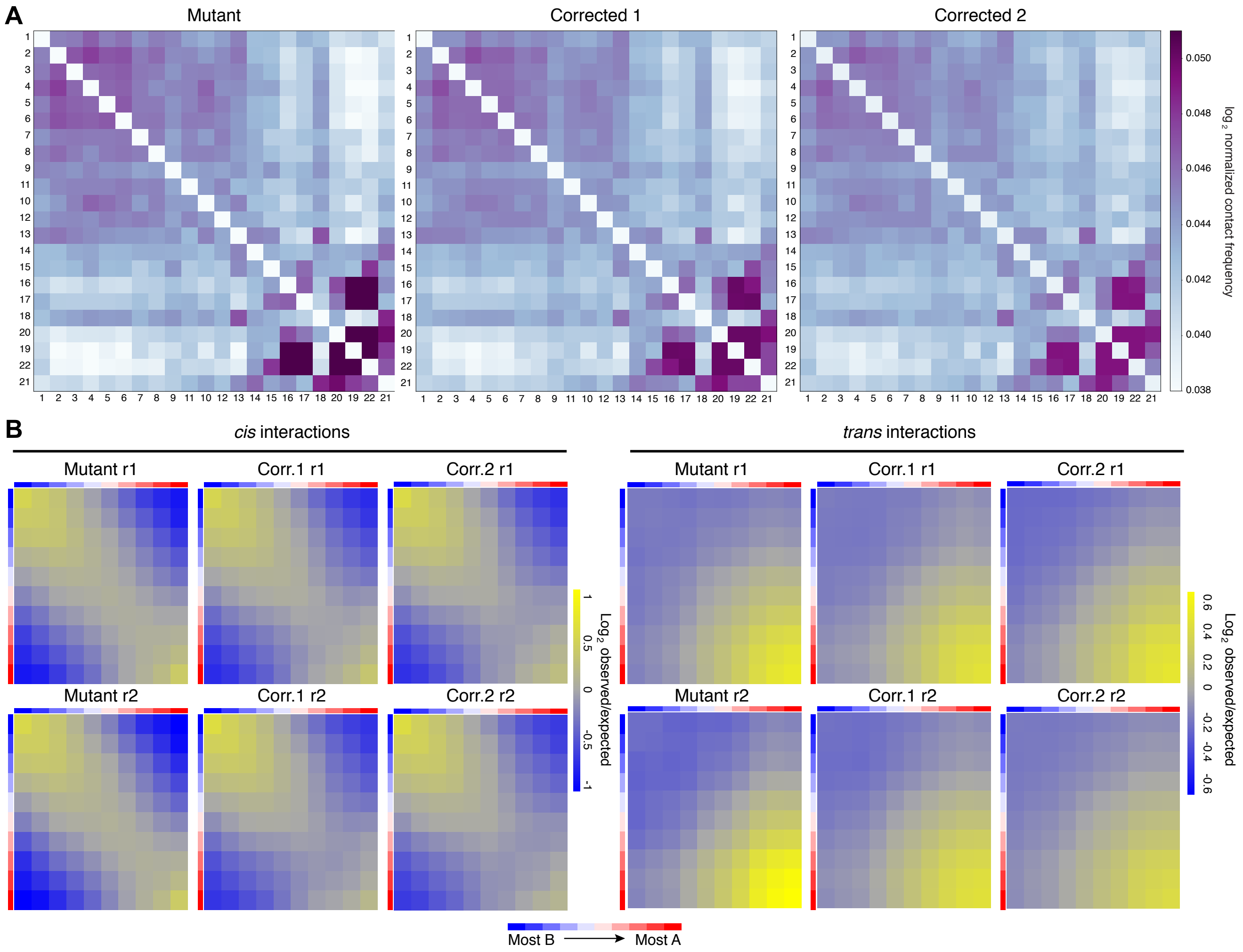


**Supplemental Figure 3. Global properties of chromatin topology in lamin A/C haploinsufficient hiPSC-CMs.** (**A**) Representative heatmaps of contact matrices between chromosomes. Autosomes are ranked based on their size from left to right and top to bottom. (**B**) Heatmaps of *cis* or *trans* interactions between active (A) and inactive (B) chromatin compartments. 500 Kb genomic bins were assigned to ten deciles based on their PC1 score from the linear dimensionality reduction of the Hi-C matrix (from most B to most A; Table S4), and average observed/expected distance normalized scores for each pair of deciles were calculated. r1/r2: biological replicate 1/2 (independent differentiations)

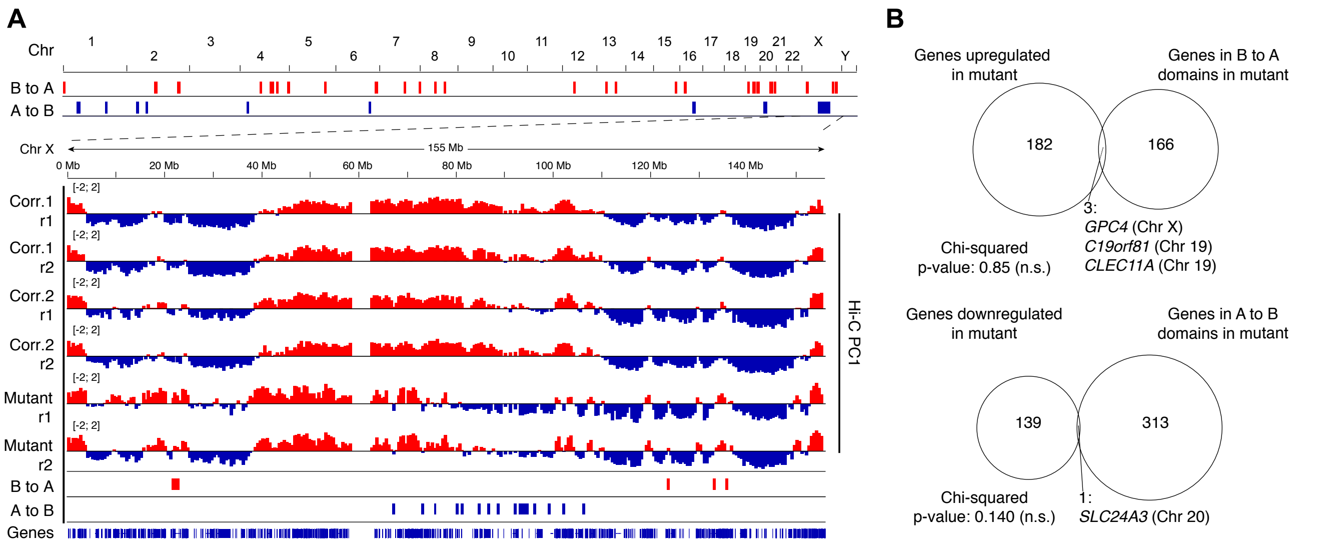


**C**

Day 14 hiPSC-CMs

| Gene | Genomic location | Expr. Mut. (RPKM) | Expr. Corr.1 (RPKM) | Expr. Corr.2 (RPKM) | Avg. log FC (Mut/Corr) |
|---|---|---|---|---|---|
| *PCDHGB4* | 5:141330570-141512981 | 2.78 | 0.13 | 0.07 | 4.86 |
| *CH16-140OK24.5* | 5:141181398-141195642 | 0.19 | 0.63 | 0.00 | 3.73 |
| *C19orf81* | 19:50621306-50719450 | 22.37 | 1.22 | 2.41 | 3.71 |
| *CLEC11A* | 19:50723328-50725718 | 4.97 | 0.16 | 0.94 | 3.67 |
| *FNDC1* | 6:159169396-159272109 | 0.61 | 0.04 | 0.07 | 3.57 |
| *SYT3* | 19:50621306-50719450 | 0.85 | 0.07 | 0.19 | 2.91 |
| *PCDHGA8* | 5:141330570-141512981 | 0.42 | 0.05 | 0.10 | 2.62 |
| *PCDHGB5* | 5:141330570-141512981 | 0.23 | 0.05 | 0.04 | 2.24 |
| *GPC4* | X:133300102-133415490 | 15.19 | 3.39 | 4.45 | 1.97 |
| *DNAH3* | 16:20933110-21180616 | 0.06 | 0.00 | 0.05 | 1.96 |
| *TMEM130* | 7:98846487-98870771 | 1.04 | 0.28 | 0.26 | 1.96 |
| *PCDHGB3* | 5:141330570-141512981 | 0.15 | 0.02 | 0.06 | 1.92 |
| *PCDHA10* | 5:140786135-141012347 | 0.90 | 0.27 | 0.21 | 1.90 |
| *PCDHGA2* | 5:141330570-141512981 | 0.37 | 0.20 | 0.05 | 1.82 |
| *PCDHGA5* | 5:141330570-141512981 | 0.14 | 0.04 | 0.05 | 1.80 |
| *PCDHGA7* | 5:141330570-141512981 | 0.16 | 0.05 | 0.05 | 1.70 |
| *PCDHGB2* | 5:141330570-141512981 | 0.31 | 0.09 | 0.10 | 1.68 |
| *ACSM3* | 16:20610242-20925006 | 2.92 | 0.83 | 1.07 | 1.63 |
| *CACNA1A* | 19:13206441-13623990 | 3.06 | 1.24 | 0.79 | 1.62 |
| *PCDHGA3* | 5:141330570-141512981 | 0.38 | 0.19 | 0.10 | 1.49 |
| *PCDHGA10* | 5:141330570-141512981 | 5.20 | 3.91 | 0.94 | 1.44 |
| *LRRC4B* | 19:50516891-50568045 | 4.23 | 1.39 | 2.04 | 1.33 |
| *ZNF534* | 19:52429186-52452315 | 0.25 | 0.13 | 0.09 | 1.25 |
| *NFIX* | 19:12995607-13098796 | 0.37 | 0.14 | 0.17 | 1.25 |
| *AOX1* | 2:200585867-200677064 | 0.32 | 0.14 | 0.13 | 1.21 |
| *PCDHA2* | 5:140786135-141012347 | 0.02 | 0.01 | 0.01 | 1.14 |
| *PCDHB15* | 5:141222931-141249365 | 4.50 | 2.41 | 1.96 | 1.05 |
| *CTD-2616J11.11* | 19:51345168-51369003 | 0.40 | 0.13 | 0.29 | 1.04 |
| *PCDHB5* | 5:141135217-141138625 | 0.52 | 0.29 | 0.22 | 1.02 |

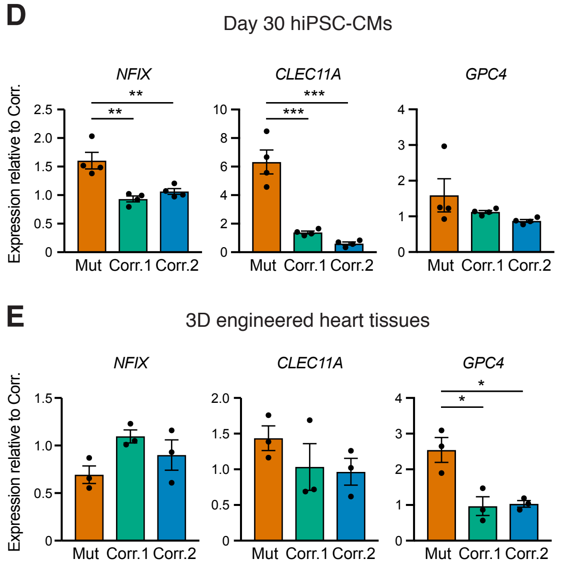


**Supplemental Figure 4. Chromatin compartment transitions and associated gene expression changes in lamin A/C haploinsufficient hiPSC-CMs.**

(**A**) On the top, genome-wide view of 500 Kb chromatin domains transitioning from B→A or A→B in corrected→mutant hiPSC-CMs (Table S4). On the bottom, genomic tracks of chromatin compartmentalization for chromosome X. Positive and negative Hi-C matrix PC1 scores are shown in red and blue, and indicate A and B compartments, respectively. (**B**) Overlap between genes up- or downregulated in mutant hiPSC-CMs (RNA-seq on 3 differentiations; fold-change > 2; q-value < 0.05) and genes located in lamin A/C-sensitive chromatin compartments (Table S4). p-values for the significance of observed overlaps were calculated by chi-squared test (n.s.: non-significant). (**C**) Average expression levels of genes found in lamin A/C-sensitive B chromatin compartments. Data is from the RNA-seq analyses of day 14 hiPSC-CMs (Table S1), and in contrast to panel B all genes with an average fold-change > 2 for mutant *versus* corrected are reported (no filter by q-value). The genomic location of all genes is indicated, and colored backgrounds highlight genes found in three genomic hotspots where lamin A/C haploinsufficiency leads to compartment transitions from B to A resulting in upregulation of multiple genes located within (19p13.13, 19q13.33, and 5q31.3; see Fig. 6E and Fig. 7B). (**D-E**) RT-qPCR validation of gene expression changes in hiPSC-

CMs matured by culture *in vitro* for 30 days (D) or by generation of 3D-EHTs (E). Differences *versus* mutant were calculated by one-way ANOVA with post-hoc Holm-Sidak binary comparisons (* = $p < 0.05$, ** = $p < 0.01$; *** = $p < 0.001$ n = 4 differentiations for panel D, and n = 3 3D-EHT batches for panel E; average ± SEM).

**Supplemental Table 6. Genotyping strategies for the characterization of CRISPR/Cas9-edited hiPSCs.**

| PCR | Primer | Sequence | Location | Band WT[1] (bp) | Band TARGET[2] (bp) | Band PLASMID[3] (bp) |
|---|---|---|---|---|---|---|
| LOCUS/LoA | LMNA_FW1 | CCCCTGCTCAAACATCCTCA | Left to 5' HAR | 2181 | 5260 or LoA | - |
| | LMNA_REV1 | TGCAATCAGAGCTTCCCCAG | Right to 3' HAR | | | |
| 5'-INT | LMNA_FW1 | CCCCTGCTCAAACATCCTCA | Left to 5' HAR | - | 1154 | - |
| | PB3-P2 | GCGACGGATTCGCGCTATTTAGAAAG | 5' ITR | | | |
| 5'-INT+ 5'-LOCUS | LMNA_FW1 | CCCCTGCTCAAACATCCTCA | Left to 5' HAR | 1836 | 1154 (4915) | - |
| | PB3-P2 | GCGACGGATTCGCGCTATTTAGAAAG | 5' ITR | | | |
| | LMNA_REV2 | CTGTGGTTGTGGGGACACTT | 3' HAR | | | |
| 3'-INT | PB5-P2 | CGTCAATTTTACGCATGATTATCTTTAAC | 3' ITR | - | 1173 | - |
| | LMNA_REV1 | TGCAATCAGAGCTTCCCCAG | Right to 3' HAR | | | |
| 3'-INT + 3'-LOCUS | LMNA_FW2 | CCCAAAAGTACCCCAGGCAT | 5' HAR | 1900 | 1173 (4979) | - |
| | PB5-P2 | CGTCAATTTTACGCATGATTATCTTTAAC | 3' ITR | | | |
| | LMNA_REV1 | TGCAATCAGAGCTTCCCCAG | Right to 3' HAR | | | |
| 5'-BB | M13-R49 | GAGCGGATAACAATTTCACACAGG | Plasmid 5' | - | - | 1157 |
| | PB3-P2 | GCGACGGATTCGCGCTATTTAGAAAG | 5' ITR | | | |
| 3'-BB | PB5-P2 | CGTCAATTTTACGCATGATTATCTTTAAC | 3' ITR | - | - | 1127 |
| | M13-F43 | AGGGTTTTCCCAGTCACGACGTT | Plasmid 3' | | | |

[1] Amplicon from the wild-type allele, or from the corrected allele following excision of the piggyBac cassette.

[2] Amplicon from the gene-targeted allele following integration of the piggyBac cassette.

[3] Amplicon from the targeting plasmid after its random integration.

**Supplemental Table 7. RT-qPCR primers.**

| Gene - isoform | Forward primer | Reverse primer | Amplicon size (bp) |
|---|---|---|---|
| *ATP2A2* | ATGGGGCTCCAACGAGTTAC | TTTCCTGCCATACACCCACAA | 224 |
| *C19orf81* | CCGAACGAGGAGGCCTGAC | AGTATCCACCCCCTCGAGCC | 147 |
| *CACNA1A* | AGGCATCCCTTTGATGGAGC | CTGAGTTTTGACCATGCGGC | 222 |
| *CACNA1C* | GCCGCTGCAGGAGAGTTTTA | CCCACATGTGCAAGACCACA | 151 |
| *CLEC11A* | TCCGGAATCTCCCTTCCCTT | TTATTGGCACCGGACCTAGC | 156 |
| *GPC4* | TGGACCGACTGGTTACTGATG | TGGTTTGCTGGTGTCAACCT | 227 |
| *IPTR1* | CGGAGCAGGGTATTGGAACA | GGTCCACTGAGGGCTGAAACT | 122 |
| *KCNA4* | GCCAAACCCGAGTGATTCCT | AAGCACTTCACCGTTCCCC | 156 |
| *KCNA5* | TAAGGAAGAGCAGGGCACTC | TTGCTCTTGGCCTTGACGTT | 174 |
| *KCND3* | ACCACACTGGGATACGGAGA | TTCGAACTGCCTGTTTTGGC | 221 |
| *KCNH2* | CAGCTCAACAGGCTGGAGAC | CTTCTTGGGGAAGCTCTGGG | 250 |
| *KCNJ2* | GTGCGAACCAACCGCTACA | CCAGCGAATGTCCACACAC | 234 |
| *KCNQ1* | AGCCTCACTCATTCAGACCG | TCTCCAGGAGTCACCCCATT | 192 |
| *LMNA* - Lamin A | CCATCACCACCACGGCTC | GTGACCAGATTGTCCCCGAA | 243 |
| *LMNA* - Lamin C | CAACTCCACTGGGGAAGAAGTG | CGGCGGCTACCACTCAC | 123 |
| *LRRC4B* | GGGCATTGCTCTTCCTCTGG | CGGATCACCTGGATGCCGTT | 241 |
| *MESP1* | TCGAAGTGGTTCCTTGGCAGAC | CCTCCTGCTTGCCTCAAAGTGTC | 163 |
| *MYH6* | GCCCTTTGACATTCGCACTG | GGTTTCAGCAATGACCTTGCC | 103 |
| *NANOG* | TTTGTGGGCCTGAAGAAAACT | AGGGCTGTCCTGAATAAGCAG | 116 |
| *NFIX* | ACAGCAGTCTCAGTCCTGGT | GGGCTGGGGGATTTTTCCAT | 119 |
| *NKX2-5* | GAGCCGAAAAGAAAGCCTGAA | CACCGACACGTCTCACTCAG | 149 |
| *PCDHA10* | GGAAGCTGCTGGATCGTGAA | AATCAGAAGCGTTGAGCCGT | 216 |
| *PCDHB15* | CTGACGGGAGGCTCTGAAAG | CAGACGGTCGGAAAGCTACA | 132 |
| *PCDHGB4* | AGCAGCACTGCACAGATACA | CCCGGGTACACGTTTTCTGA | 107 |
| *PDGFRA* | GCTCACTTCACTCTCCCCAAAG | CCGGCGTTCCTGGTCTTAG | 153 |
| *POU5F1* | GGGTTCTATTTGGGAAGGTAT | TTCATTGTTGTCAGCTTCCT | 131 |
| *RPLP0* | GGCGTCCTCGTGGAAGTGAC | GCCTTGCGCATCATGGTGTT | 255 |
| *RYR2* | ACAACAGAAGCTATGCTTGGC | GAGGAGTGTTCGATGACCACC | 250 |
| *SCN5A* | AACGGCACCTCTGATGTGTT | ACCTGAGGGTCTGCTGATAGA | 199 |
| *SLC8A1* | AGACCTGGCTTCCCACTTTG | TGGCAAATGTGTCTGGCACT | 101 |
| *SYT3* | GACTACGACTGCATCGGGCA | TGCTGCCTTTTGTGAAGCTG | 178 |
| *TBXT* | CAAATCCTCATCCTCAGTTTG | GTCAGAATAGGTTGGAGAATTG | 143 |
| *TNNT2* | TTCACCAAAGATCTGCTCCTCGCT | TTATTACTGGTGTGGAGTGGGTGTGG | 166 |

**Supplemental Video 1 (online). Spontaneous contractions of mutant hiPSC-CMs.**
Spontaneous contractions of mutant hiPSC-CM monolayers at day 14 of differentiation. Phase-contrast images were acquired at 30 frames per second (fps) and are shown at 30 fps.

**Supplemental Video 2 (online). Spontaneous contractions of corrected hiPSC-CMs.**
Spontaneous contractions of corrected control hiPSC-CM monolayers (from Corr.1 hiPSCs) at day 14 of differentiation. Phase-contrast images were acquired at 30 fps and are shown at 30 fps.

**Supplemental Video 3 (online). Calcium fluxes indicated by Fluo-4 fluorescence in monolayers of mutant hiPSC-CMs electrically paced at 1 Hz.**
Calcium fluxes in electrically-paced (1 Hz) mutant hiPSC-CM monolayers at day 30 of differentiation. Fluo-4 epifluorescence images were acquired at 20 fps and are shown at 20 fps.

**Supplemental Video 4 (online). Calcium fluxes indicated by Fluo-4 fluorescence in monolayers of corrected hiPSC-CMs electrically paced at 1 Hz.**
Calcium fluxes in electrically-paced (1 Hz) corrected control hiPSC-CM monolayers (from Corr.1 hiPSCs) at day 30 of differentiation. Fluo-4 epifluorescence images were acquired at 20 fps and are shown at 20 fps.

**Supplemental Video 5 (online). Contraction correlation quantification (CCQ) analyses in monolayers of mutant hiPSC-CMs electrically paced at 1 Hz.**
Contractions of electrically-paced (1 Hz) mutant hiPSC-CM monolayers at day 30 of differentiation. Phase-contrast images were acquired at 30 fps and are shown at 30 fps. Superimposed white arrows indicate the displacement vectors calculated by CCQ analysis.

**Supplemental Video 6 (online). Contraction correlation quantification (CCQ) analyses in monolayers of corrected hiPSC-CMs electrically paced at 1 Hz.**
Contractions of electrically-paced (1 Hz) corrected control hiPSC-CM monolayers (from Corr.1 hiPSCs) at day 30 of differentiation. Phase-contrast images were acquired at 30 fps and are shown at 30 fps. Superimposed white arrows indicate the displacement vectors calculated by CCQ analysis.

**Supplemental Video 7. (online). Contractions of 3D-EHTs from mutant hiPSC-CMs.**
Contractions of electrically-paced (1 Hz) mutant 3D-EHTs analyzed 4 weeks after casting. Phase-contrast images were acquired at 65 fps and are shown at 65 fps.

**Supplemental Video 8 (online). Contractions of 3D-EHTs from corrected hiPSC-CMs.**
Contractions of electrically-paced (1 Hz) corrected control 3D-EHTs (from Corr.1 hiPSCs) analyzed 4 weeks after casting. Phase-contrast images were acquired at 65 fps and are shown at 65 fps.
